# Mathematical Modeling of AA Amyloidosis: Coupling SAA–HDL Binding Dynamics with Path-Dependent Renal Aging

**DOI:** 10.64898/2026.02.19.706923

**Authors:** Andrey V. Kuznetsov

## Abstract

Why does chronic inflammation cause AA amyloidosis in only some patients, even when serum amyloid A (SAA) remains elevated in many? This study proposes that disease progression is governed by two linked nonlinear processes: saturation of the finite SAA-binding capacity of high-density lipoprotein (HDL), followed by autocatalytic aggregation of amyloid A (AA) in the kidney. A hypothesis-generating mathematical model is developed that couples reversible SAA–HDL binding to renal filtration of free SAA, lysosomal cleavage to AA fragments, Finke–Watzky aggregation, oligomer deposition, and fibril turnover. As total SAA rises beyond the effective HDL binding capacity, the model predicts a disproportionate increase in free, filterable SAA available for renal amyloid formation. A dimensionless reduction reveals two kinetic regimes separated by a threshold at which autocatalytic oligomer production balances oligomer loss. Below this threshold, aggregation is nucleation-limited; above it, the system becomes supply-limited and is controlled mainly by upstream SAA delivery. In the zero-primary-nucleation limit, the two steady-state branches exchange stability at this threshold, giving a transcritical bifurcation that finite primary nucleation smooths into a continuous transition. The baseline parameter set lies in the supply-limited regime, explaining why total SAA concentration and effective HDL binding-site capacity have the largest local sensitivities, with SAA–HDL affinity and lysosomal cleavage next in rank, whereas the aggregation rate constants have small local effects but can still determine which regime applies. The model further predicts that the temporal pattern of inflammation matters: intermittent SAA flares can produce several-fold more renal damage than sustained inflammation with the same mean SAA, and earlier SAA suppression leaves a lower renal biological-age index, a calibrated measure of cumulative damage, than later suppression. Published SAA–outcome data are consistent with the predicted SAA dependence. By linking HDL buffering to renal SAA delivery, nonlinear aggregation, and cumulative injury, the model provides a quantitative framework for identifying amyloidogenic transitions and testing how SAA control can alter renal disease progression.

## 1. Introduction

Systemic amyloid A (AA) amyloidosis is a severe, often fatal complication of chronic inflammatory conditions, including rheumatoid arthritis, familial Mediterranean fever, and chronic infections (Lachmann and Hawkins, 2006). The disease involves extracellular deposition of insoluble fibrillar aggregates in major organs, predominantly the kidneys, where amyloid accumulation can lead to progressive organ dysfunction and, in advanced disease, organ failure (Lachmann et al., 2007), (Westermark and Westermark, 2009). Although chronic inflammation affects many patients, clinically significant AA amyloidosis develops in only a minority of them (Kluve-Beckerman et al., 2022). The clinical course of renal amyloidosis is well characterized, but the molecular thresholds that govern the transition from chronic inflammation to pathological amyloid deposition remain unclear.

The amyloid deposits originate from serum amyloid A (SAA), an acute-phase apolipoprotein synthesized by the liver. Under normal physiological conditions and mild inflammation, SAA circulates predominantly in association with high-density lipoprotein (HDL) particles (Frame and Gursky, 2016), (Webb, 2021), (Ji et al., 2020). Structural studies suggest that HDL binding protects SAA from misfolding by sequestering the amyloidogenic N-terminal region against the lipoprotein surface (Frame and Gursky, 2016). Thus, HDL serves as a molecular chaperone, maintaining SAA in a soluble, non-aggregating state.

This protective mechanism, however, has a finite capacity. Biophysical studies show that SAA binds to HDL through a rigid hydrophobic interface, creating a saturable interaction with stoichiometric constraints on SAA occupancy of HDL particles (Frame and Gursky, 2016). When severe or prolonged inflammation increases SAA production beyond the binding capacity of HDL, saturation occurs, and excess SAA remains free in plasma without lipid protection (Webb, 2021). These lipid-poor SAA species are no longer protected from misfolding.

While the large SAA–HDL complex cannot pass through the glomerular filtration barrier and remains in the bloodstream, free SAA (∼12 kDa) is sufficiently small to undergo renal filtration (Röcken and Shakespeare, 2002). Once filtered into the renal compartment and taken up by renal cells, unbound SAA becomes locally concentrated and undergoes proteolytic cleavage, creating conditions that thermodynamically favor AA protein aggregation (Röcken and Shakespeare, 2002), (Gillmore et al., 2001).

Despite these mechanistic insights, predicting amyloidosis development in individual patients remains difficult. Serum SAA levels correlate poorly with amyloid burden; many patients with chronically elevated SAA never develop disease, while others progress rapidly. This suggests that a threshold may exist at which the protective capacity of HDL is exceeded and autocatalytic fibril growth becomes self-sustaining (Knowles et al., 2009).

Building on previously developed models of transthyretin and AL amyloidosis (Kuznetsov, 2025a), (Kuznetsov, 2026a), this work develops a novel mathematical model that couples SAA–HDL binding dynamics with amyloid aggregation kinetics. The model calculates the concentration of free, filterable SAA from HDL binding capacity and dissociation constant (*K*_*d*_) determined under the assumption of reversible binding equilibrium between free SAA monomers (*S*_*free*_) and available binding sites on HDL particles (Artl et al., 2000). The computed free SAA concentration serves as a parameter in a system of differential equations governing renal filtration, mesangial accumulation, and conversion to fibrillar aggregates through primary nucleation, autocatalytic conversion, and oligomer deposition into fibrils (Cohen et al., 2013). The model quantifies how HDL saturation triggers pathological amyloid accumulation, providing a framework for evaluating potential therapeutic targets.

The framework developed here is intended as a hypothesis-generating, mechanistic exploration of how SAA–HDL buffering couples to renal amyloid kinetics; it is not a patient-specific predictive tool. Several kinetic parameters, in particular the two Finke–Watzky rate constants, cannot presently be determined for AA protein from published data and are therefore varied over representative ranges drawn from other amyloidogenic systems (Table 2, footnotes c and d). Accordingly, quantitative outputs should be interpreted primarily as comparisons among scenarios computed under a common set of assumptions.

Renal biological age is used as a calibrated damage index rather than as a directly measured physiological clock, and the index is allowed to fall when fibrillar deposits regress, but never below a floor set by chronological age and accumulated oligomer exposure (Section 2.3). Section 2.5 further shows that combinations of parameters, rather than individual parameter values alone, determine the aggregation regime.

## 2. Materials and models

### 2.1. Governing equations

#### 2.1.1. Equations describing SAA–HDL association

In the model, the interaction between SAA and HDL is treated as a saturable binding event. Because HDL particles can accommodate several SAA monomers under acute inflammatory conditions (Frame and Gursky, 2016), (Thomas and Sorci-Thomas, 2015), the model represents this multivalent binding through the total concentration of effective SAA-binding sites, [*H*]_*tot*_, rather than through the HDL particle concentration itself. Since each effective site is assumed to bind at most one SAA molecule, a particle that carries several SAA molecules contributes several sites to [*H*]_*tot*_. Site heterogeneity and cooperative binding are not represented. Thus, [*H*]_*tot*_ equals the HDL particle concentration multiplied by the effective number of SAA-binding sites per particle and is an inferred model quantity rather than a directly measured concentration. At the mean HDL particle concentration of 13.4 µM measured by calibrated ion mobility analysis (Hutchins et al., 2014), the baseline [*H*]_*tot*_ = 30 µM (Table 2) corresponds to approximately 2.2 effective sites per HDL particle, consistent with the multivalent binding noted above.

The second binding parameter is the dissociation constant, *K*_*d*_. This constant is not taken directly from *in vitro* binding measurements; instead, it is calibrated for the chosen [*H*]_*tot*_ so that Eq. (13) reproduces the measured fraction of lipid-free SAA in serum (see footnote f of Table 2). A reversible binding equilibrium is assumed between free SAA monomers (*S* _*free*_) and available binding sites on HDL particles (*H* _*free*_)

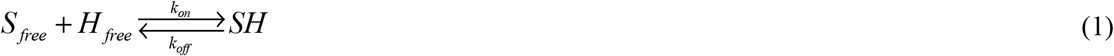

where *SH* represents the SAA–HDL complex.

The equilibrium dissociation constant for *SH* is determined as:

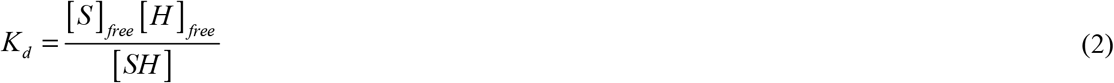

The total SAA concentration (HDL-bound plus free) is then given by:

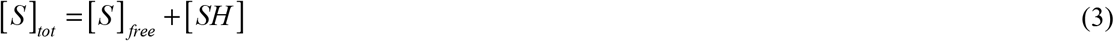

and the total concentration of SAA-binding sites on HDL particles, comprising both occupied and unoccupied sites, is calculated as:

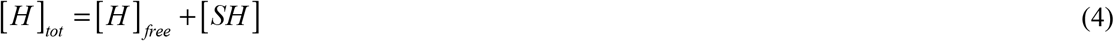

Substituting Eq. (4) into Eq. (2) yields the following expression for the equilibrium dissociation constant:

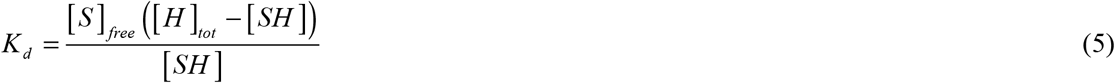

Solving Eq. (5) for the concentration of the SAA–HDL complex yields:

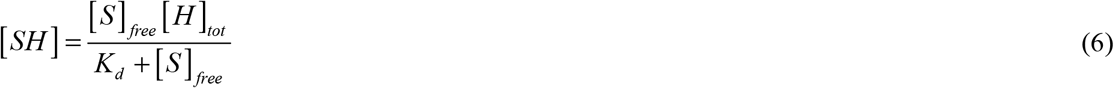

Substituting the expression for [*SH*] from Eq. (3) into Eq. (6) results in:

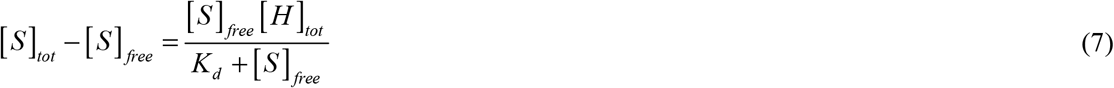

Solving for [*S*]_*free*_ yields a quadratic equation:

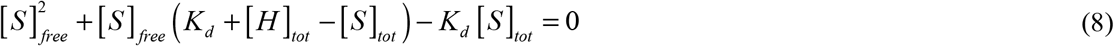

Because concentrations must be non-negative, the physically meaningful root is selected:

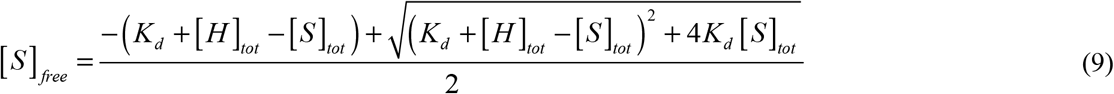

The fraction of free SAA can then be calculated as:

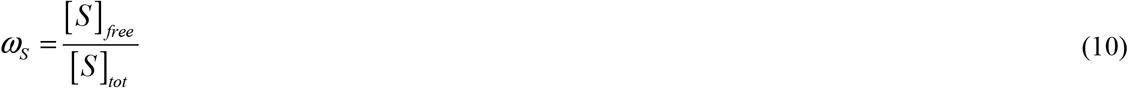

Similarly, the fraction of unoccupied HDL-associated binding sites can be calculated as:

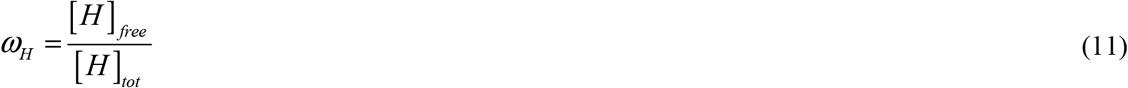

Where

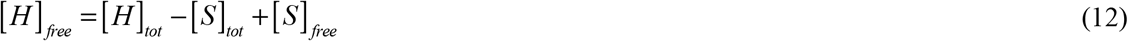

Substituting [*S*]_*free*_ from Eq. (10) and [*SH*] from Eq. (3) into Eq. (5) yields the following expression for the equilibrium dissociation constant:

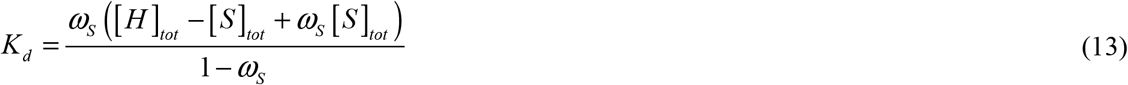

#### 2.1.2. Development of conservation equations describing SAA deposition in the kidneys and AA amyloid fibril formation

The SAA–HDL complex is assumed to be unable to pass through the glomerular filtration barrier into the renal compartment and is retained in the blood. Free SAA monomers, however, are assumed to undergo renal filtration, after which they are internalized by mesangial and proximal tubular cells (Kon et al., 2021). Within the acidic environment of the lysosome (pH 3.5–4.5), these unbound monomers may escape complete degradation and instead contribute to oligomer formation (Jayaraman et al., 2017).

Subsequent proteolytic cleavage sheds the C-terminus, generating the amyloidogenic AA fragment (Röcken and Shakespeare, 2002), which is assumed to nucleate within the supersaturated renal interstitium to form the insoluble fibrils characteristic of the disease.

Upon renal uptake, internalized SAA is processed in endo-lysosomal compartments by cathepsins. Cathepsin B trims the SAA C-terminus to generate N-terminal AA-like fragments, while cathepsin D and other lysosomal proteases promote more complete degradation. Limited C-terminal proteolysis destabilizes the SAA structure, exposes the amyloidogenic N-terminal region, and promotes formation of AA-type aggregates that accumulate in tissues (Lu et al., 2014). This cleavage process is simulated by the following equation:

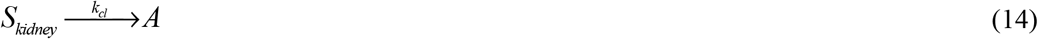

The mathematical model comprises conservation equations for SAA monomers entering the kidneys from the blood, AA monomers derived from the lysosomal cleavage of SAA precursors, free AA oligomers, and AA oligomers deposited into fibrils. A lumped capacitance formulation is employed, assuming spatial homogeneity of all AA species within the kidneys (a simplifying assumption that neglects regional variations in renal AA processing). Time is the sole independent variable and is defined as the patient’s calendar age minus the age at diagnosis of AA amyloidosis, which is identified with the disease-onset time *t*_*AA*,*onset*_ of Eq. (49). Dependent variables are defined in Table 1, and model parameters are summarized in Table 2.

**Table 1.** Model-dependent variables.

| Symbol | Definition | Unit |
|---|---|---|
| [*A*] | Molar concentration of AA fragments (monomers) in kidneys | µM |
| [*B*] | Molar concentration of free (non-deposited) AA oligomers in kidneys | µM |
| [*I*] | Molar concentration of AA oligomers deposited into AA amyloid fibrils in kidneys | µM |
| [*S*]<sub>*free*</sub> | Molar concentration of free (unbound) SAA in blood plasma | µM |
| [*S*]<sub>*kidney*</sub> | Molar concentration of SAA in kidneys | µM |
| [*H*]<sub>*free*</sub> | Molar concentration of free (available) binding sites on HDL | µM |
| [*SH*] | Molar concentration of bound SAA (SAA–HDL complex) | µM |

Formulating the conservation equation for SAA monomers entering the kidneys from the blood and normalizing by *V*_*kidneys*_ yields the following equation:

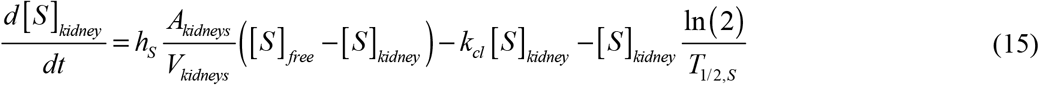

In Eq. (15), the first term on the right-hand side represents the retention rate of filtered SAA monomers, where *h*_*S*_ *A*_*kidneys*_ ([*S*]_*free*_ − [*S*]_*kidney*_) denotes the total rate of renal deposition of monomers. The second term represents the loss of SAA monomers deposited in the kidneys through their cleavage into N-terminal AA fragments susceptible to aggregation (see Eq. (14)). The third term accounts for the clearance of SAA monomers by proteolytic enzymes within the lysosomes of proximal tubular cells (Miura et al., 1985).

The numerical solution demonstrates that [*S*]_*kidney*_ rapidly approaches a constant steady-state value that is maintained for most of the disease course (Fig. S1), validating the use of the steady-state approximation. Applying steady-state conditions to Eq. (15) yields:

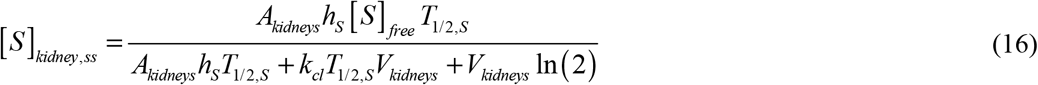

Cleavage of SAA (see Eq. (14)) releases the N-terminal AA fragment, which is assumed to undergo rapid self-association into oligomers and eventually form insoluble amyloid fibrils. Soluble prefibrillar AA oligomers act as pathogenic intermediates that bridge elevated SAA levels to tissue AA fibril deposition (Jayaraman et al., 2017). To simulate the oligomerization of the AA protein, the Finke–Watzky (F–W) model was used. This minimal two-step model, which captures both nucleation and autocatalytic growth, has been widely used to characterize aggregation kinetics of various amyloidogenic proteins (Morris et al., 2008), (Iashchishyn et al., 2017), (Finke et al., 2020). Within this framework, the first pseudo-elementary step corresponds to the primary nucleation of AA oligomers. The second step represents autocatalytic conversion; pre-existing oligomers act as autocatalytic species that accelerate the conversion of free AA monomers (Aragonès Pedrola et al., 2025).

The first pseudo-elementary step describes primary nucleation of AA protein oligomers:

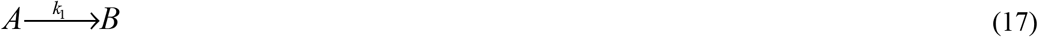

In the F–W model, the oligomeric product is treated as an autocatalytic species capable of promoting further aggregation. The model does not explicitly distinguish between monomeric and oligomeric species with respect to molecular size or mass. The second pseudo-elementary step represents the autocatalytic conversion of monomeric AA protein into oligomeric forms:

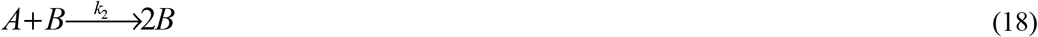

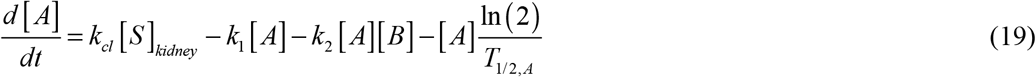

In Eq. (19), the first term on the right-hand side represents the generation of AA fragments (monomers) via the cleavage of SAA (as defined in Eq. (14)). The second and third terms account for the depletion of AA monomers due to their conversion into oligomers through nucleation and autocatalytic processes, respectively. Finally, the fourth term describes the degradation of AA monomers by proteases.

The formation of AA fibrils in the kidneys from oligomers is modeled analogously to colloidal suspension coagulation (Boltachev and Ivanov, 2020). In this model, free AA oligomers (*B*) are assigned a half-deposition time *θ*_1/2, *B*_, defined as the time required for half of the free oligomers to be incorporated into AA fibrils. Applying the principle of conservation of the number of free AA oligomers within the kidneys and normalizing the resulting equation by *V*_*kidneys*_

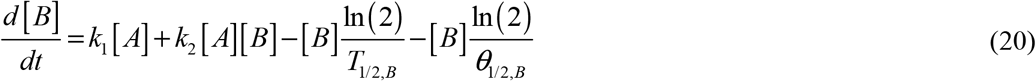

The first two terms on the right-hand side of Eq. (20) represent the generation of free AA oligomers from AA monomers through nucleation and autocatalytic processes, respectively. These terms are analogous to the second and third terms in Eq. (19) but with opposite signs. The third term accounts for the clearance of oligomers due to their finite half-life. The fourth term represents the loss of free oligomers through their incorporation into fibrils.

As with [*S*]_*kidney*_, the concentrations of AA monomers and free oligomers ([*A*] and [*B*]) rapidly approach steady-state values that are maintained for most of the disease course (Figs. S2 and S3). This justifies applying the steady-state approximation, under which Eqs. (19) and (20) become:

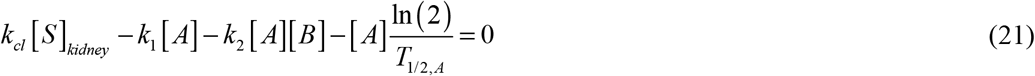

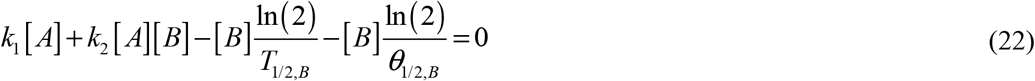

The summation of Eqs. (21) and (22) yields:

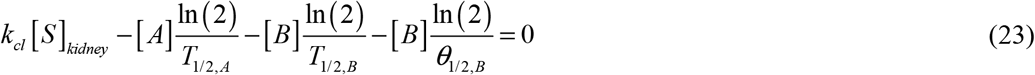

Solving Eq. (23) for [*A*] yields:

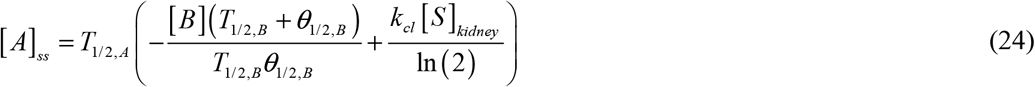

Substituting Eq. (24) into Eq. (22) and solving for [*B*], while selecting the positive root of the quadratic equation, yields:

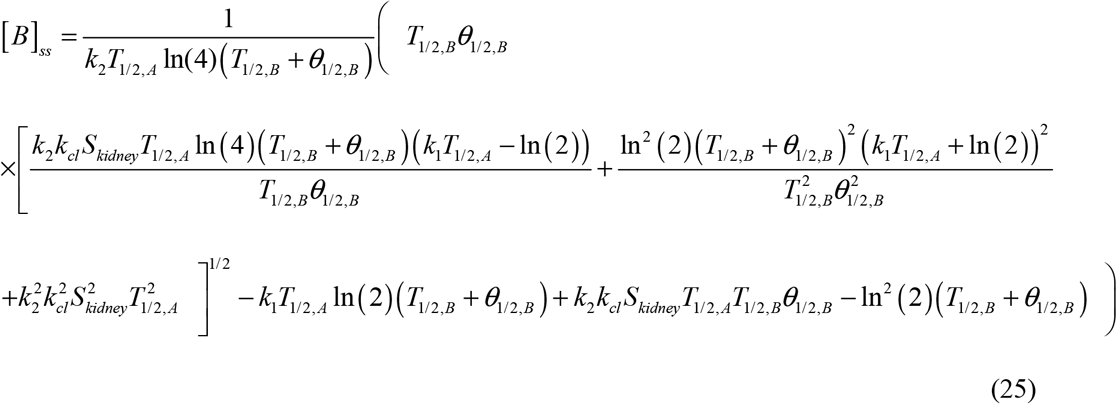

Applying the conservation principle to AA oligomers sequestered into fibrillar structures within the kidneys and normalizing by *V*_*kidneys*_ yields the following governing equation:

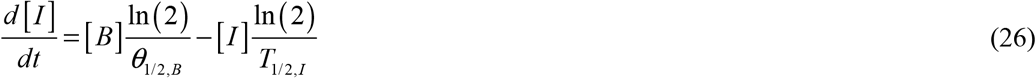

where [*I*] is the concentration of deposited oligomers.

In Eq. (26), the first term on the right-hand side corresponds to the fourth term in Eq. (20) with opposite sign, representing deposition into fibrils. The second term accounts for the clearance of deposited AA oligomers from fibrils based on their half-life.

Since [*B*](*t*) rapidly attains its steady-state value [*B*]_*ss*_, as confirmed by the numerical solution in Fig. S3, solving Eq. (26) subject to the initial condition (32d) yields:

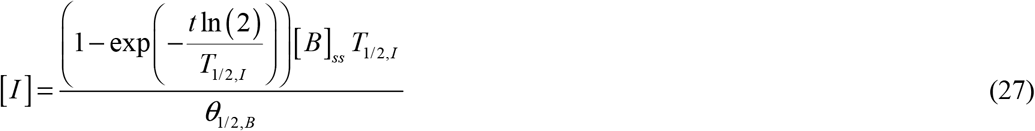

Eq. (27) shows that as time approaches infinity, the fibril concentration approaches a steady-state value:

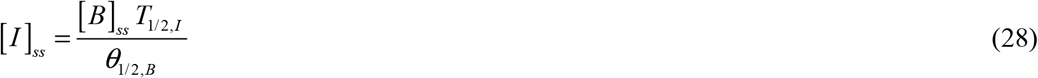

The steady-state fibril burden given by Eq. (28) is determined by the balance between fibril formation and fibril clearance. AA amyloid fibrils are highly stable structures with long half-lives; their burden falls appreciably only when deposition is curtailed for a period comparable to the fibril half-life, as is observed clinically when circulating SAA is suppressed (footnote m to Table 2). The time required for the fibril concentration to reach 90% of its steady-state value can be derived from Eq. (27) as:

**Table 2.**
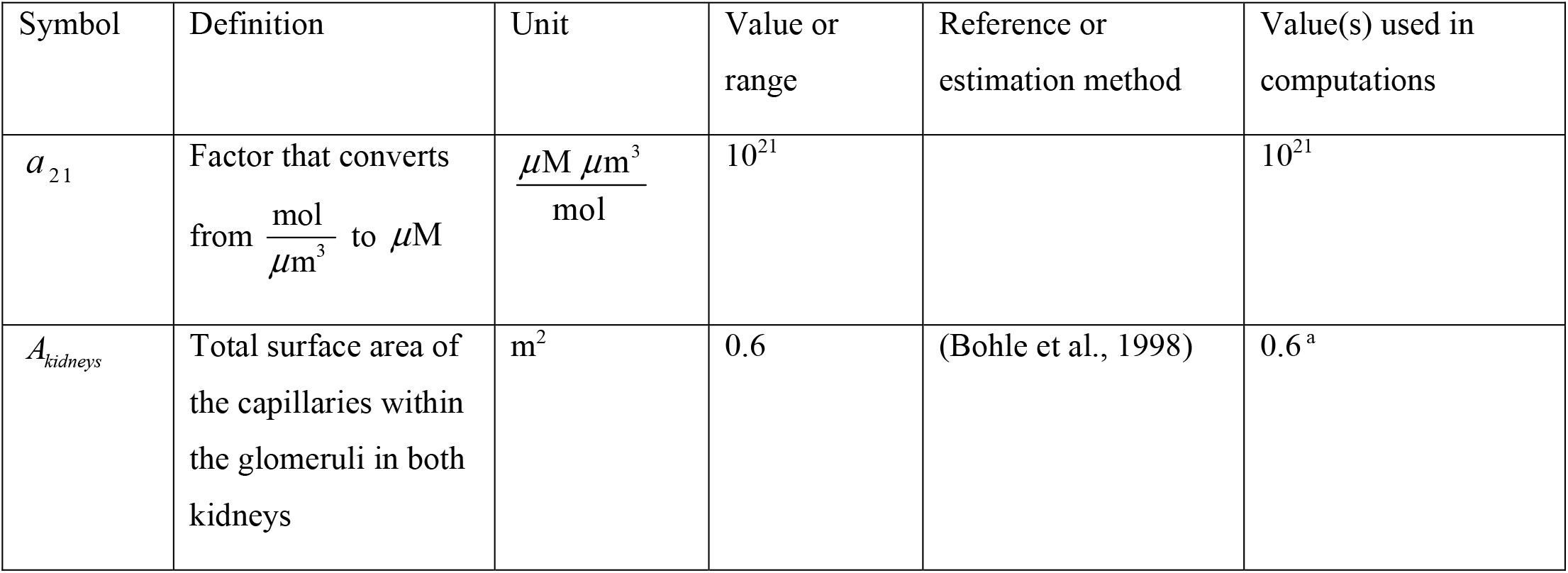
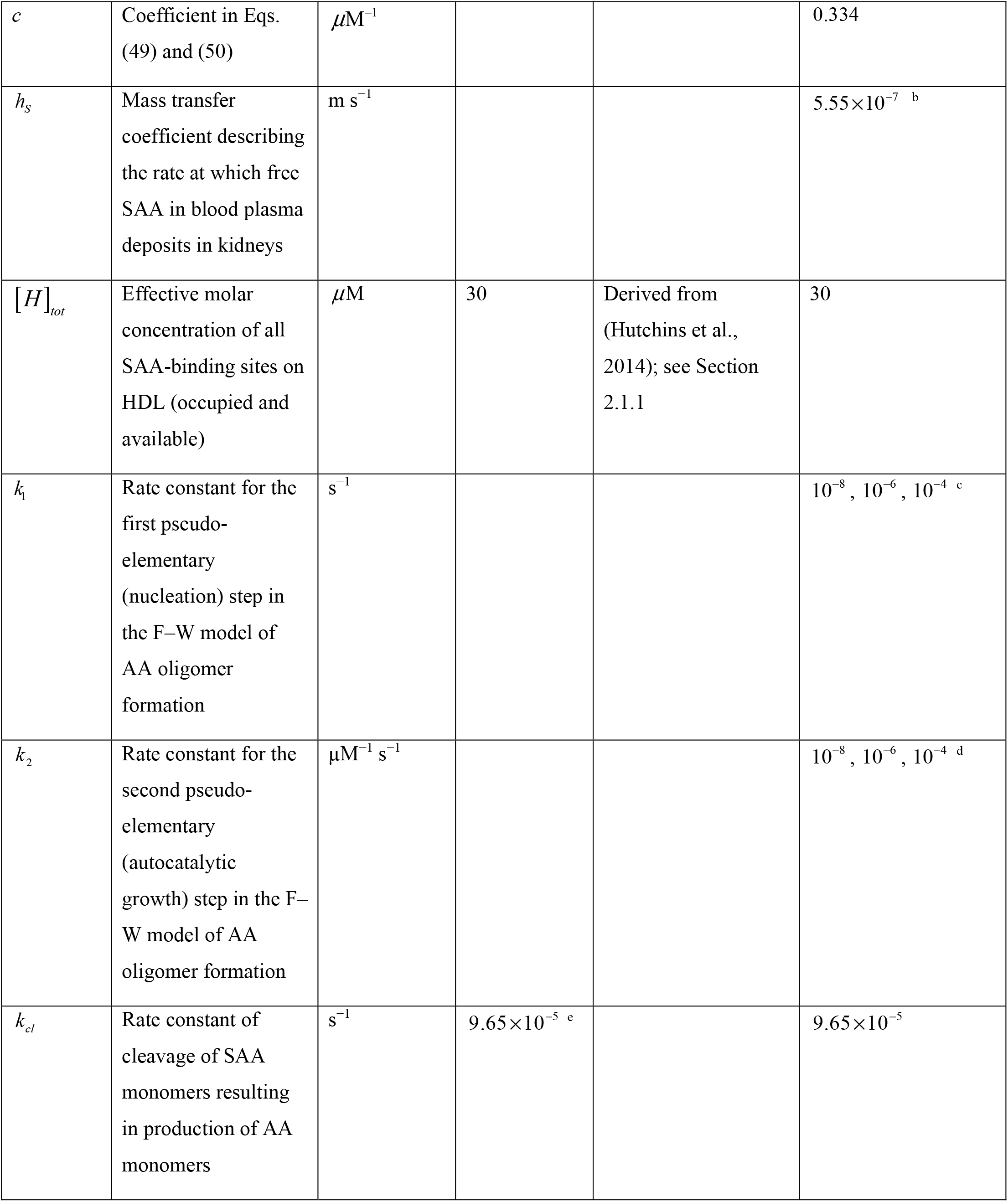
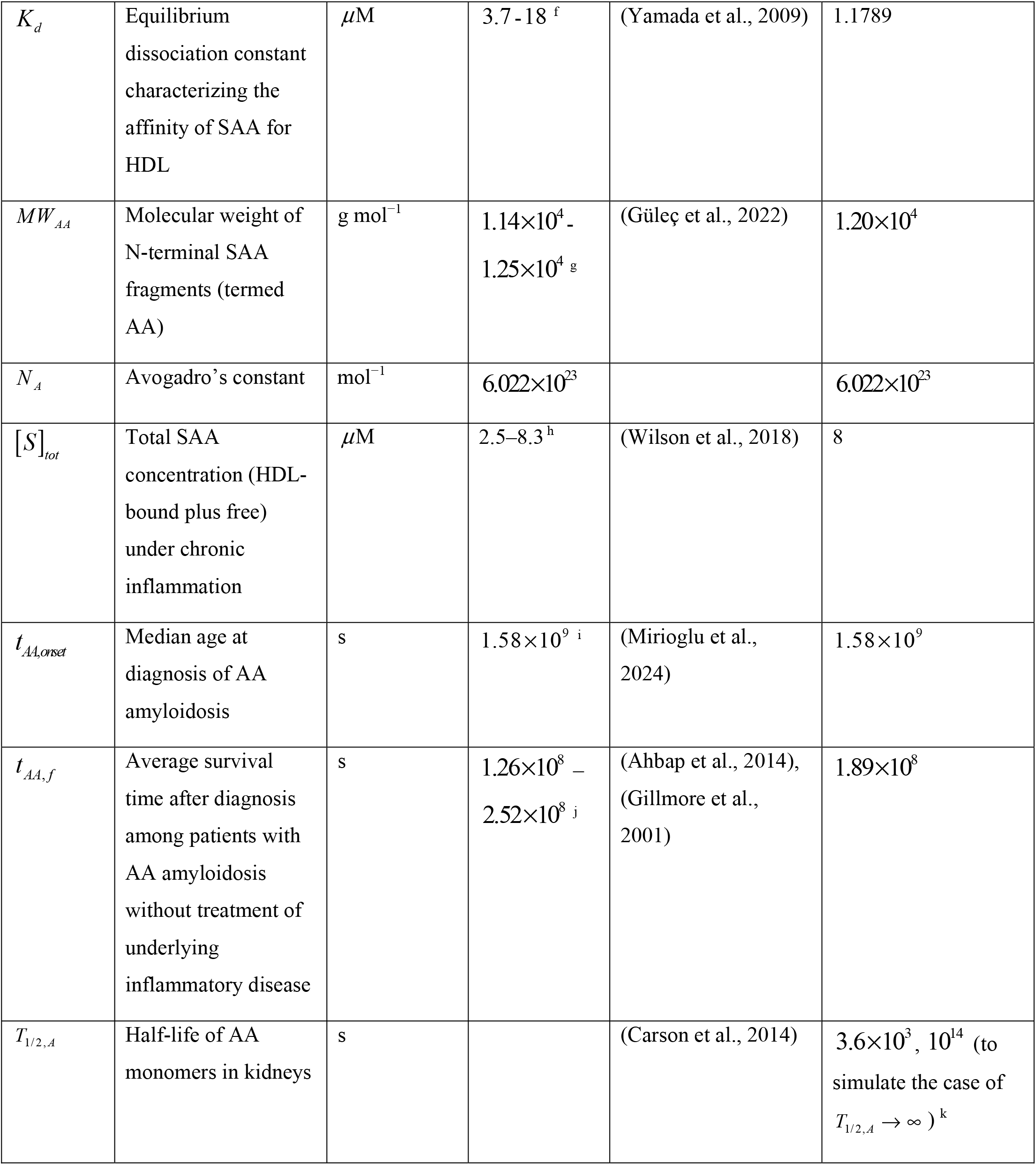
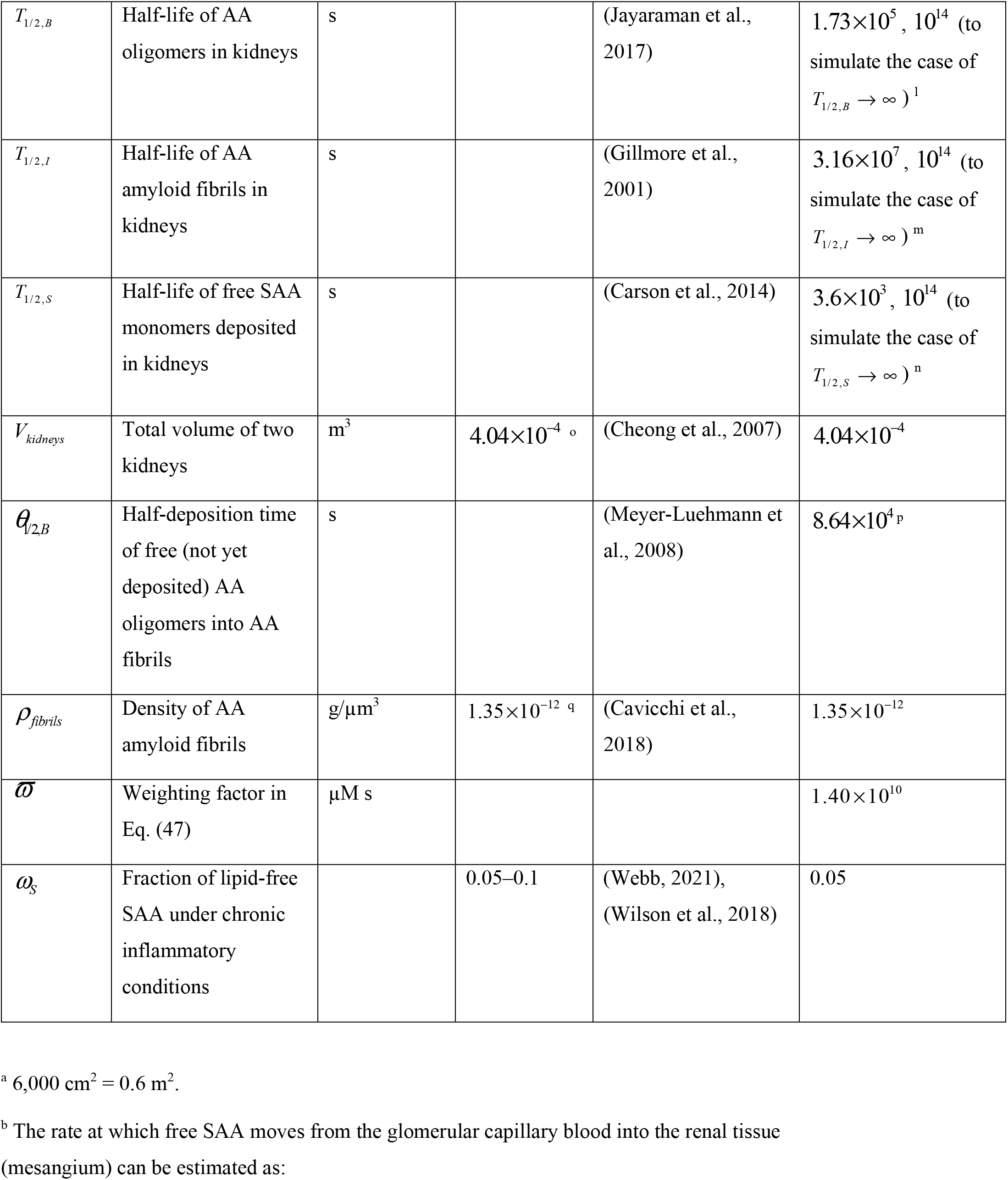
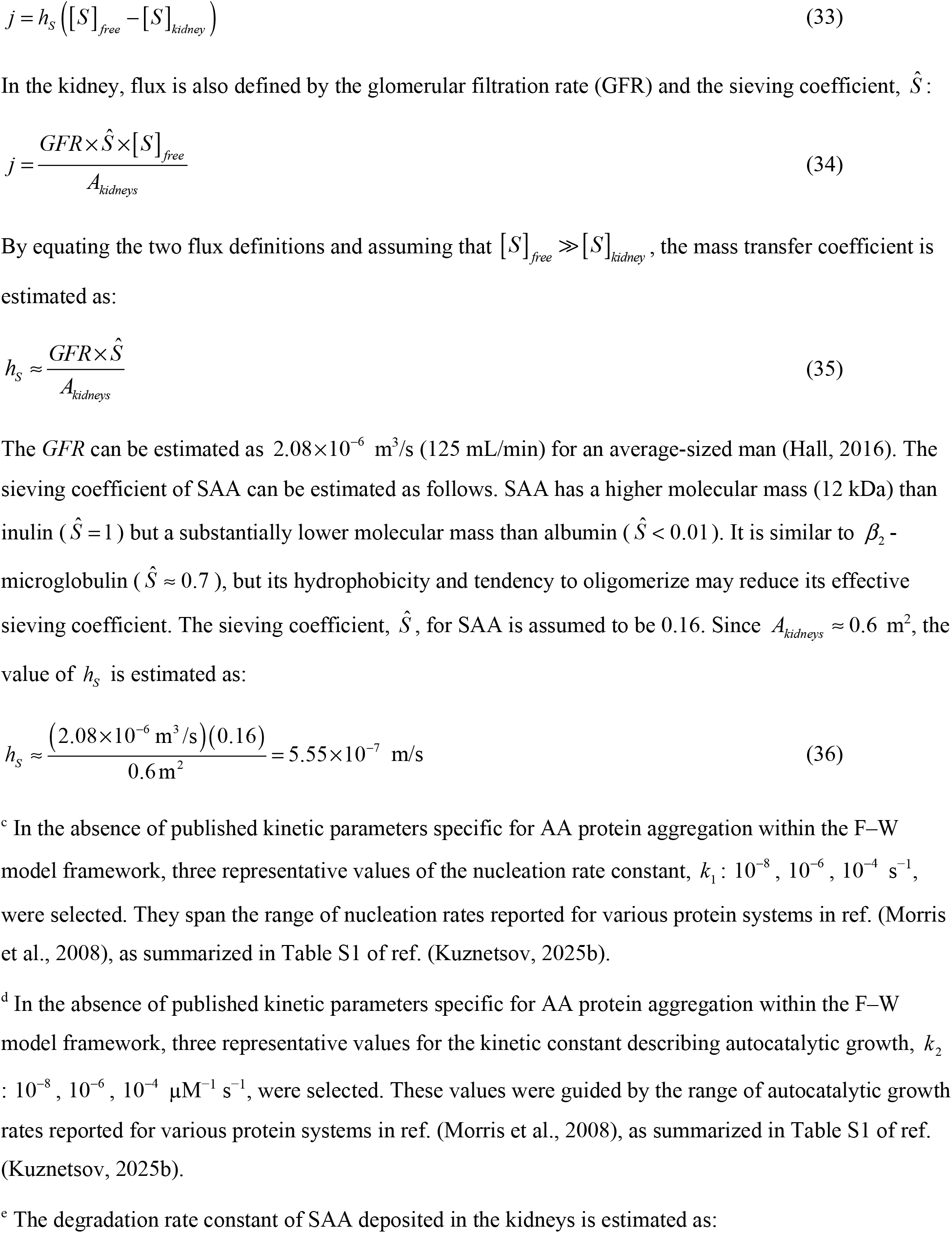
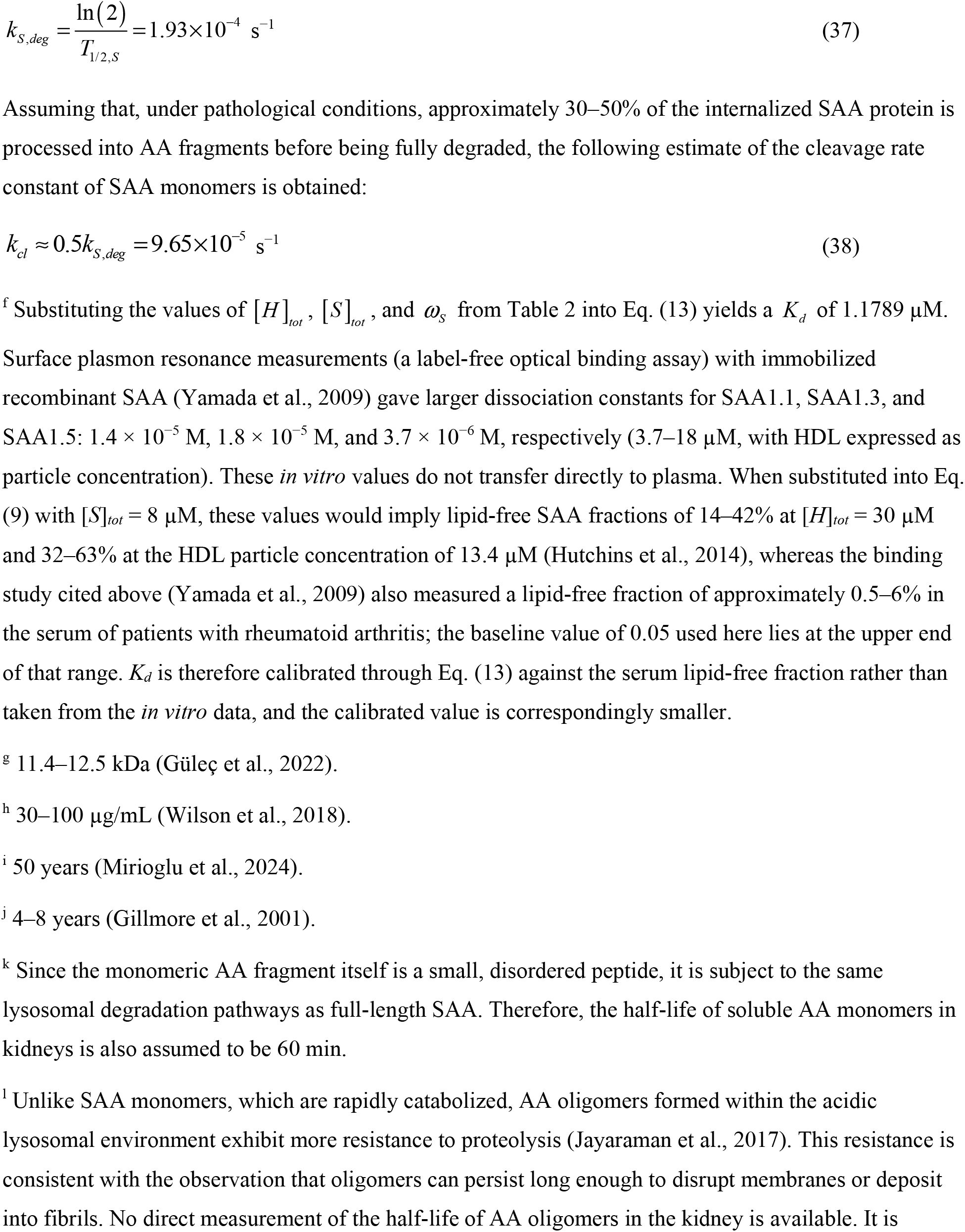
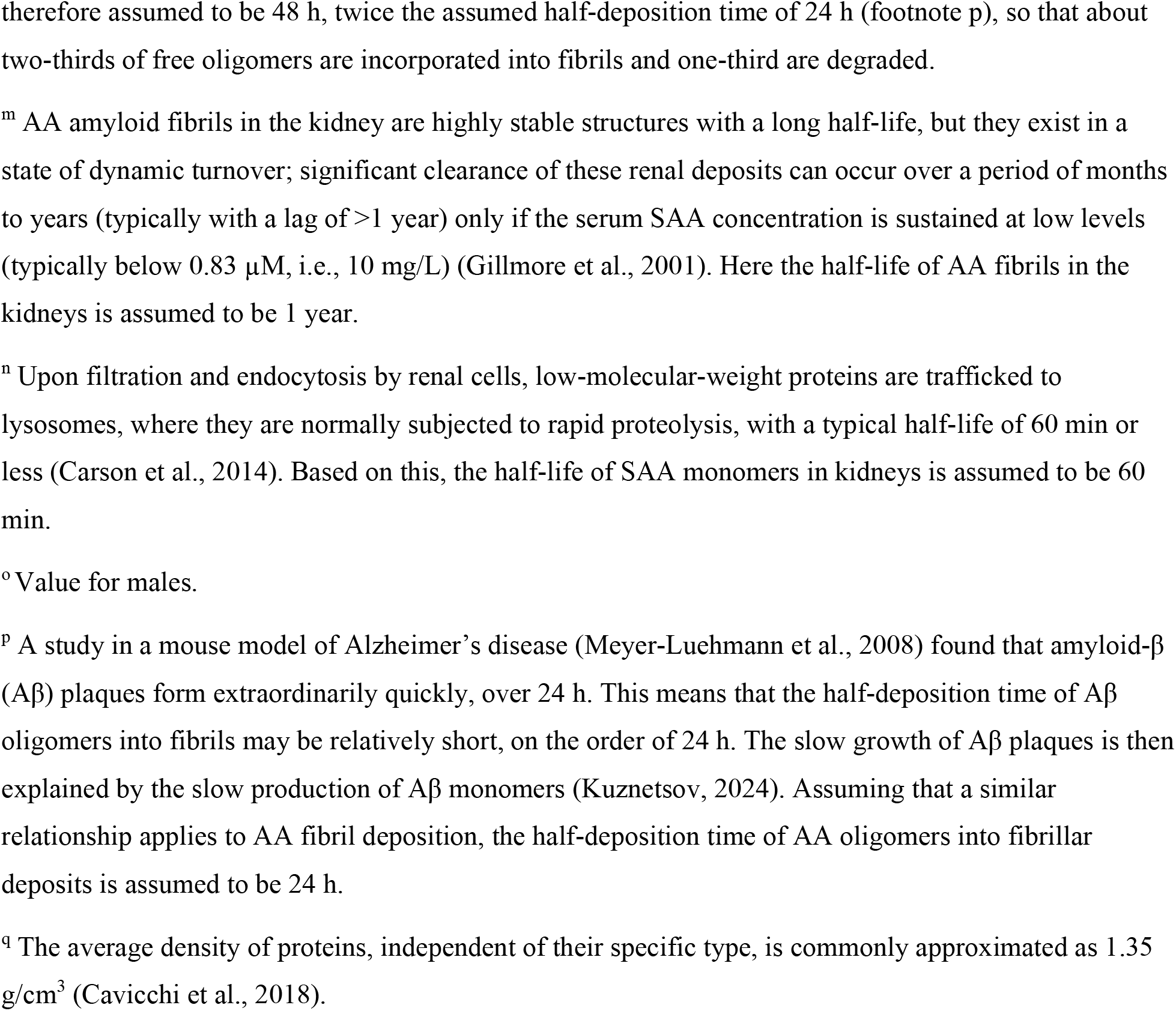
Model parameters and their estimated values.

| Symbol | Definition | Unit | Value or range | Reference or estimation method | Value(s) used in computations |
|---|---|---|---|---|---|
| *a*<sub>21</sub> | Factor that converts from $\frac{\mathrm{mol}}{\mu\mathrm{m}^3}$ to *µ*M | $\frac{\mu\mathrm{M}\ \mu\mathrm{m}^3}{\mathrm{mol}}$ | 10<sup>21</sup> |  | 10<sup>21</sup> |
| *A*<sub>*kidneys*</sub> | Total surface area of the capillaries within the glomeruli in both kidneys | m<sup>2</sup> | 0.6 | (Bohle et al., 1998) | 0.6<sup>a</sup> |
| *c* | Coefficient in Eqs. (49) and (50) | *µ*M<sup>−1</sup> |  |  | 0.334 |
| *h*<sub>*S*</sub> | Mass transfer coefficient describing the rate at which free SAA in blood plasma deposits in kidneys | m s<sup>−1</sup> |  |  | 5.55×10<sup>−7</sup> <sup>b</sup> |
| [*H*]<sub>*tot*</sub> | Effective molar concentration of all SAA-binding sites on HDL (occupied and available) | *µ*M | 30 | Derived from (Hutchins et al., 2014); see Section 2.1.1 | 30 |
| *k*<sub>1</sub> | Rate constant for the first pseudo-elementary (nucleation) step in the F–W model of AA oligomer formation | s<sup>−1</sup> |  |  | 10<sup>−8</sup>, 10<sup>−6</sup>, 10<sup>−4</sup> <sup>c</sup> |
| *k*<sub>2</sub> | Rate constant for the second pseudo-elementary (autocatalytic growth) step in the F–W model of AA oligomer formation | µM<sup>−1</sup> s<sup>−1</sup> |  |  | 10<sup>−8</sup>, 10<sup>−6</sup>, 10<sup>−4</sup> <sup>d</sup> |
| *k*<sub>*cl*</sub> | Rate constant of cleavage of SAA monomers resulting in production of AA monomers | s<sup>−1</sup> | 9.65×10<sup>−5</sup> <sup>e</sup> |  | 9.65×10<sup>−5</sup> |
| *K*<sub>*d*</sub> | Equilibrium dissociation constant characterizing the affinity of SAA for HDL | *µ*M | 3.7 - 18<sup>f</sup> | (Yamada et al., 2009) | 1.1789 |
| *MW*<sub>*AA*</sub> | Molecular weight of N-terminal SAA fragments (termed AA) | g mol<sup>−1</sup> | 1.14×10<sup>4</sup> - 1.25×10<sup>4</sup> <sup>g</sup> | (Güleç et al., 2022) | 1.20×10<sup>4</sup> |
| *N*<sub>*A*</sub> | Avogadro’s constant | mol<sup>−1</sup> | 6.022×10<sup>23</sup> |  | 6.022×10<sup>23</sup> |
| [*S*]<sub>*tot*</sub> | Total SAA concentration (HDL-bound plus free) under chronic inflammation | *µ*M | 2.5–8.3<sup>h</sup> | (Wilson et al., 2018) | 8 |
| *t*<sub>*AA,onset*</sub> | Median age at diagnosis of AA amyloidosis | s | 1.58×10<sup>9</sup> <sup>i</sup> | (Mirioglu et al., 2024) | 1.58×10<sup>9</sup> |
| *t*<sub>*AA,f*</sub> | Average survival time after diagnosis among patients with AA amyloidosis without treatment of underlying inflammatory disease | s | 1.26×10<sup>8</sup> – 2.52×10<sup>8</sup> <sup>j</sup> | (Ahbap et al., 2014), (Gillmore et al., 2001) | 1.89×10<sup>8</sup> |
| *T*<sub>1/2,*A*</sub> | Half-life of AA monomers in kidneys | s |  | (Carson et al., 2014) | 3.6×10<sup>3</sup>, 10<sup>14</sup> (to simulate the case of *T*<sub>1/2,*A*</sub> → ∞ )<sup>k</sup> |
| *T*<sub>1/2,*B*</sub> | Half-life of AA oligomers in kidneys | s |  | (Jayaraman et al., 2017) | 1.73×10<sup>5</sup>, 10<sup>14</sup> (to simulate the case of *T*<sub>1/2,*B*</sub> → ∞ )<sup>l</sup> |
| *T*<sub>1/2,*I*</sub> | Half-life of AA amyloid fibrils in kidneys | s |  | (Gillmore et al., 2001) | 3.16×10<sup>7</sup>, 10<sup>14</sup> (to simulate the case of *T*<sub>1/2,*I*</sub> → ∞ )<sup>m</sup> |
| *T*<sub>1/2,*S*</sub> | Half-life of free SAA monomers deposited in kidneys | s |  | (Carson et al., 2014) | 3.6×10<sup>3</sup>, 10<sup>14</sup> (to simulate the case of *T*<sub>1/2,*S*</sub> → ∞ )<sup>n</sup> |
| *V*<sub>*kidneys*</sub> | Total volume of two kidneys | m<sup>3</sup> | 4.04×10<sup>−4</sup> <sup>o</sup> | (Cheong et al., 2007) | 4.04×10<sup>−4</sup> |
| *θ*<sub>1/2,*B*</sub> | Half-deposition time of free (not yet deposited) AA oligomers into AA fibrils | s |  | (Meyer-Luehmann et al., 2008) | 8.64×10<sup>4</sup> <sup>p</sup> |
| *ρ*<sub>*fibrils*</sub> | Density of AA amyloid fibrils | g/µm<sup>3</sup> | 1.35×10<sup>−12</sup> <sup>q</sup> | (Cavicchi et al., 2018) | 1.35×10<sup>−12</sup> |
| *ϖ* | Weighting factor in Eq. (47) | µM s |  |  | 1.40×10<sup>10</sup> |
| *ω*<sub>*S*</sub> | Fraction of lipid-free SAA under chronic inflammatory conditions |  | 0.05–0.1 | (Webb, 2021), (Wilson et al., 2018) | 0.05 |

<sup>a</sup> 6,000 cm<sup>2</sup> = 0.6 m<sup>2</sup>.
<sup>b</sup> The rate at which free SAA moves from the glomerular capillary blood into the renal tissue (mesangium) can be estimated as:
$j = h_S\left(\left[S\right]_{free} - \left[S\right]_{kidney}\right)$ (33)
In the kidney, flux is also defined by the glomerular filtration rate (GFR) and the sieving coefficient, *Ŝ*:
$j = \frac{GFR \times \hat{S} \times \left[S\right]_{free}}{A_{kidneys}}$ (34)
By equating the two flux definitions and assuming that [*S*]<sub>*free*</sub> ≫ [*S*]<sub>*kidney*</sub>, the mass transfer coefficient is estimated as:
$h_S \approx \frac{GFR \times \hat{S}}{A_{kidneys}}$ (35)
The *GFR* can be estimated as 2.08×10<sup>−6</sup> m<sup>3</sup>/s (125 mL/min) for an average-sized man (Hall, 2016). The sieving coefficient of SAA can be estimated as follows. SAA has a higher molecular mass (12 kDa) than inulin ( *Ŝ* = 1 ) but a substantially lower molecular mass than albumin ( *Ŝ* < 0.01 ). It is similar to *β*<sub>2</sub>-microglobulin ( *Ŝ* ≈ 0.7 ), but its hydrophobicity and tendency to oligomerize may reduce its effective sieving coefficient. The sieving coefficient, *Ŝ*, for SAA is assumed to be 0.16. Since *A*<sub>*kidneys*</sub> ≈ 0.6 m<sup>2</sup>, the value of *h*<sub>*S*</sub> is estimated as:
$h_S \approx \frac{\left(2.08\times10^{-6}\ \mathrm{m}^3/\mathrm{s}\right)\left(0.16\right)}{0.6\,\mathrm{m}^2} = 5.55\times10^{-7}\ \mathrm{m/s}$ (36)
<sup>c</sup> In the absence of published kinetic parameters specific for AA protein aggregation within the F–W model framework, three representative values of the nucleation rate constant, *k*<sub>1</sub>: 10<sup>−8</sup> , 10<sup>−6</sup> , 10<sup>−4</sup> s<sup>−1</sup>, were selected. They span the range of nucleation rates reported for various protein systems in ref. (Morris et al., 2008), as summarized in Table S1 of ref. (Kuznetsov, 2025b).
<sup>d</sup> In the absence of published kinetic parameters specific for AA protein aggregation within the F–W model framework, three representative values for the kinetic constant describing autocatalytic growth, *k*<sub>2</sub> : 10<sup>−8</sup> , 10<sup>−6</sup> , 10<sup>−4</sup> µM<sup>−1</sup> s<sup>−1</sup>, were selected. These values were guided by the range of autocatalytic growth rates reported for various protein systems in ref. (Morris et al., 2008), as summarized in Table S1 of ref. (Kuznetsov, 2025b).
<sup>e</sup> The degradation rate constant of SAA deposited in the kidneys is estimated as:
$k_{S,deg} = \frac{\ln\left(2\right)}{T_{1/2,S}} = 1.93\times10^{-4}\ \mathrm{s}^{-1}$ (37)
Assuming that, under pathological conditions, approximately 30–50% of the internalized SAA protein is processed into AA fragments before being fully degraded, the following estimate of the cleavage rate constant of SAA monomers is obtained:
$k_{cl} \approx 0.5k_{S,deg} = 9.65\times10^{-5}\ \mathrm{s}^{-1}$ (38)
<sup>f</sup> Substituting the values of [*H*]<sub>*tot*</sub> , [*S*]<sub>*tot*</sub> , and *ω*<sub>*S*</sub> from Table 2 into Eq. (13) yields a *K*<sub>*d*</sub> of 1.1789 µM. Surface plasmon resonance measurements (a label-free optical binding assay) with immobilized recombinant SAA (Yamada et al., 2009) gave larger dissociation constants for SAA1.1, SAA1.3, and SAA1.5: 1.4 × 10<sup>−5</sup> M, 1.8 × 10<sup>−5</sup> M, and 3.7 × 10<sup>−6</sup> M, respectively (3.7–18 µM, with HDL expressed as particle concentration). These *in vitro* values do not transfer directly to plasma. When substituted into Eq. (9) with [*S*]<sub>*tot*</sub> = 8 µM, these values would imply lipid-free SAA fractions of 14–42% at [*H*]<sub>*tot*</sub> = 30 µM and 32–63% at the HDL particle concentration of 13.4 µM (Hutchins et al., 2014), whereas the binding study cited above (Yamada et al., 2009) also measured a lipid-free fraction of approximately 0.5–6% in the serum of patients with rheumatoid arthritis; the baseline value of 0.05 used here lies at the upper end of that range. *K*<sub>*d*</sub> is therefore calibrated through Eq. (13) against the serum lipid-free fraction rather than taken from the *in vitro* data, and the calibrated value is correspondingly smaller.
<sup>g</sup> 11.4–12.5 kDa (Güleç et al., 2022).
<sup>h</sup> 30–100 µg/mL (Wilson et al., 2018).
<sup>i</sup> 50 years (Mirioglu et al., 2024).
<sup>j</sup> 4–8 years (Gillmore et al., 2001).
<sup>k</sup> Since the monomeric AA fragment itself is a small, disordered peptide, it is subject to the same lysosomal degradation pathways as full-length SAA. Therefore, the half-life of soluble AA monomers in kidneys is also assumed to be 60 min.
<sup>l</sup> Unlike SAA monomers, which are rapidly catabolized, AA oligomers formed within the acidic lysosomal environment exhibit more resistance to proteolysis (Jayaraman et al., 2017). This resistance is consistent with the observation that oligomers can persist long enough to disrupt membranes or deposit into fibrils. No direct measurement of the half-life of AA oligomers in the kidney is available. It is therefore assumed to be 48 h, twice the assumed half-deposition time of 24 h (footnote p), so that about two-thirds of free oligomers are incorporated into fibrils and one-third are degraded.
<sup>m</sup> AA amyloid fibrils in the kidney are highly stable structures with a long half-life, but they exist in a state of dynamic turnover; significant clearance of these renal deposits can occur over a period of months to years (typically with a lag of >1 year) only if the serum SAA concentration is sustained at low levels (typically below 0.83 µM, i.e., 10 mg/L) (Gillmore et al., 2001). Here the half-life of AA fibrils in the kidneys is assumed to be 1 year.
<sup>n</sup> Upon filtration and endocytosis by renal cells, low-molecular-weight proteins are trafficked to lysosomes, where they are normally subjected to rapid proteolysis, with a typical half-life of 60 min or less (Carson et al., 2014). Based on this, the half-life of SAA monomers in kidneys is assumed to be 60 min.
<sup>o</sup> Value for males.
<sup>p</sup> A study in a mouse model of Alzheimer’s disease (Meyer-Luehmann et al., 2008) found that amyloid-β (Aβ) plaques form extraordinarily quickly, over 24 h. This means that the half-deposition time of Aβ oligomers into fibrils may be relatively short, on the order of 24 h. The slow growth of Aβ plaques is then explained by the slow production of Aβ monomers (Kuznetsov, 2024). Assuming that a similar relationship applies to AA fibril deposition, the half-deposition time of AA oligomers into fibrillar deposits is assumed to be 24 h.
<sup>q</sup> The average density of proteins, independent of their specific type, is commonly approximated as 1.35 g/cm<sup>3</sup> (Cavicchi et al., 2018).

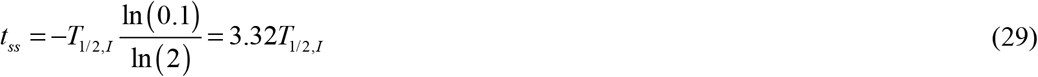

In the limit of *T*_1/2,*I*_ →∞, Eq. (26) reduces to:

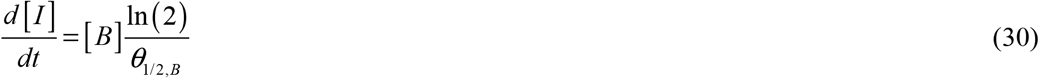

Under the assumption that the steady-state value [*B*]_*ss*_ is achieved rapidly, application of the initial condition (32d) to Eq. (30) gives:

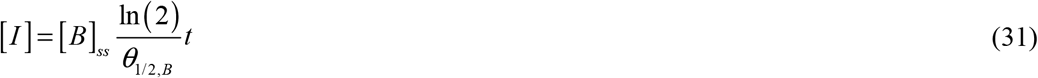

Eq. (31) predicts unbounded growth of [*I*], consistent with the assumption that fibril degradation is negligible.

Eqs. (15), (19), (20), and (26) are solved subject to the following initial conditions:

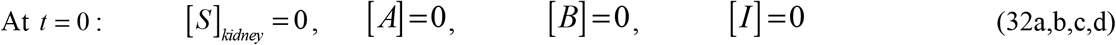

The simulation spans several years, consistent with the clinical timescale of renal AA fibril deposition (Ahbap et al., 2014). Because the long-term behavior of the system is insensitive to the initial conditions over this time interval, all four species in Eq. (32) are initialized at zero.

The numerical solution methodology is documented in Section S1 of the Supplemental Materials.

Provenance of the model parameters. Because the model combines quantities measured for SAA with quantities transferred from other amyloidogenic systems, the parameters of Table 2 are grouped here according to their origin. (i) Directly measured: the total SAA concentration under chronic inflammation and the lipid-free SAA fraction (Wilson et al., 2018), (Webb, 2021); the molecular weight of the AA fragment (Güleç et al., 2022); the *in vitro* SAA–HDL dissociation constants of three SAA1 isotypes (Yamada et al., 2009), which are not used directly (footnote f of Table 2); the HDL particle concentration (Hutchins et al., 2014), from which the effective concentration of SAA-binding sites on HDL is derived (Section 2.1.1); and the half-life of filtered low-molecular-weight proteins in the renal lysosome (Carson et al., 2014). (ii) Human renal physiology, independent of amyloidosis: glomerular capillary surface area (Bohle et al., 1998), kidney volume (Cheong et al., 2007), and the glomerular filtration rate and sieving coefficient used in footnote b (Hall, 2016). (iii) Clinical epidemiology of AA amyloidosis: median age at diagnosis (Mirioglu et al., 2024), untreated survival after diagnosis (Gillmore et al., 2001), (Ahbap et al., 2014), and the fibril half-life inferred from the regression of deposits under sustained SAA suppression (Gillmore et al., 2001). (iv) Transferred from other amyloidogenic systems and therefore not AA-specific: the two Finke–Watzky rate constants, which span the range reported for unrelated protein systems (Morris et al., 2008) and are varied over three orders of magnitude throughout this study; the half-deposition time of oligomers into fibrils, taken from amyloid-β plaque growth (Meyer-Luehmann et al., 2008); the oligomer half-life, assumed on the basis of the proteolytic resistance of SAA oligomers at lysosomal pH (Jayaraman et al., 2017); and the generic protein density (Cavicchi et al., 2018). (v) Fixed by back-calibration rather than measurement: the SAA–HDL dissociation constant, calibrated through Eq. (13) against the measured lipid-free SAA fraction; the weighting factor of Eq. (48) and the coefficient *c* of Eq. (51); see Section 2.3 and Fig. S28. Baseline local sensitivities to several transferred parameters are small, but this should not be used to discount provenance uncertainty: Section 2.5 shows that variation of the borrowed autocatalytic rate constant can move the model between different aggregation regimes. Local sensitivity at one baseline point and global uncertainty about the dynamical regime therefore answer different questions.

### 2.2. Calculating the volume occupied by AA fibrils in the kidneys

Renal AA fibril volume growth is determined by the total number of AA oligomers incorporated into the fibrils over time *t, N*_*mon in fibrils*_ (*t*) . Since the F–W model assumes equivalent molecular weights for monomeric and oligomeric species (Eq. (17)), the number of deposited monomer equivalents provides a measure of fibril volume. Adapting the approach from (Watzky et al., 2008) yields:

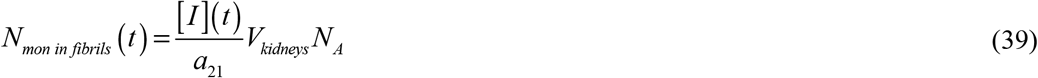

Where *N*_*A*_ denotes Avogadro’s constant.

Alternatively, the number of deposited oligomers can be computed using the volume-based expression from (Watzky et al., 2008):

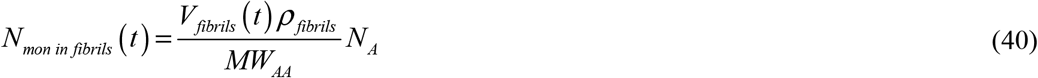

In this context, *MW*_*AA*_ represents the molecular weight of an AA monomer, *ρ* _*fibrils*_ is the density of the fibrils, and *V* _*fibrils*_ (*t*) denotes the time-dependent volume of renal AA amyloid deposits.

By equating the right-hand sides of Eqs. (39) and (40) and solving for the fraction of the baseline kidney volume occupied by fibrils, the following is obtained:

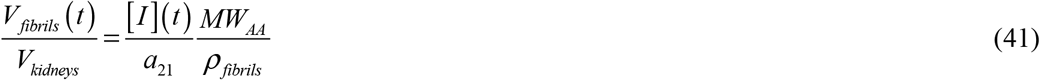

Substituting the analytical solution (27) into Eq. (41) yields:

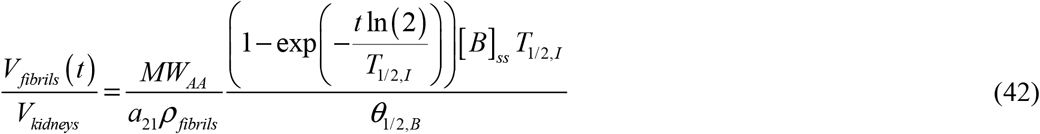

This implies that as time approaches infinity, the fraction of baseline kidney volume occupied by fibrils approaches the following steady-state value:

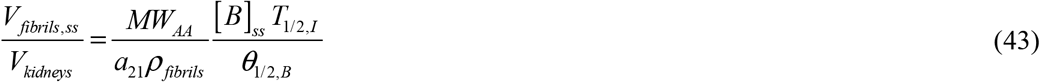

Employing the analytical solution (31) (valid for *T*_1/2,*I*_ →∞) in Eq. (41) gives:

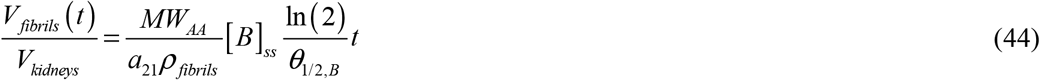

Note that in this case fibrillar volume continues to increase without bound due to the absence of fibril degradation.

The total volume of two kidneys at time *t* is given by *V*_*fibrils*_ (*t*) + *V*_*kidneys*_ .

Despite extensive histopathological characterization of AA amyloidosis, the terminal fibril burden in end-stage renal disease has not been systematically quantified in the literature. One autopsy study found 7– 11% kidney replacement by AA deposits in advanced disease, but the patient died of pancreatitis, not renal failure (Kuroda et al., 2013). A maximum renal amyloid fraction of 15% at end-stage AA amyloidosis is assumed, as higher burdens would likely be incompatible with sustained glomerular filtration.

### 2.3 Accumulated nephrotoxicity from AA oligomers, combined nephrotoxic and fibrillar effects, and a proposed index of kidney biological age

Persistently elevated circulating SAA is associated with higher whole-body AA amyloid load and worse clinical outcome (Gillmore et al., 2001).

The pathogenesis of AA amyloidosis is increasingly attributed to stable, protease-resistant SAA oligomers formed at lysosomal pH (3.5–4.5). Unlike soluble SAA or mature fibrils, these pH-stabilized intermediates have a strong capacity to bind to and permeabilize lipid membranes. A study (Srinivasan et al., 2013) found that during the aggregation phase of the pathogenic SAA1.1 isoform, there is a prolonged lag phase heavily populated by spherical oligomers. *In vitro* experiments confirmed that these prefibrillar oligomers can permeabilize synthetic membranes, resulting in membrane-associated cytotoxic effects before the formation of mature fibrils, which appear to exhibit less direct cytotoxicity than the oligomers. Within the renal environment, SAA internalization likely triggers this oligomerization, leading to lysosomal and plasma membrane disruption. Consequently, kidney damage in AA amyloidosis may be driven not only by the physical accumulation of insoluble fibrils, but also by the membrane-lytic activity of these toxic oligomers (Jayaraman et al., 2017).

The proposed criterion for characterizing nephrotoxicity follows approaches previously developed to quantify accumulated neurotoxicity from Aβ oligomers in Alzheimer’s disease (Kuznetsov, 2025c) and α- synuclein oligomers in Parkinson’s disease (Kuznetsov, 2025d), as well as accumulated cardiotoxicity in TTR amyloidosis (Kuznetsov, 2025a) and AL amyloidosis (Kuznetsov, 2026a).

Accumulated nephrotoxicity is defined as:

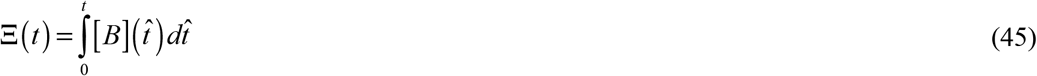

Substituting the analytical solution (25) into Eq. (45) yields:

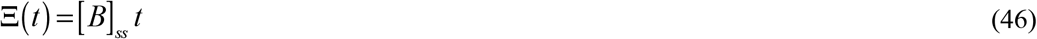

The model represents nephrotoxic injury in AA amyloidosis as arising from two mechanisms: accumulated cytotoxicity of AA oligomers and aggregation of insoluble fibrils. The combined extent of nephrotoxic damage is quantified by the following parameter:

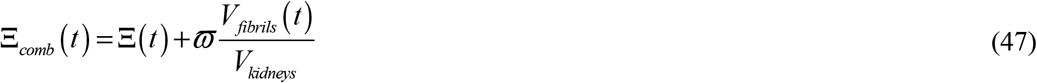

The first term of Eq. (47), the accumulated toxicity of Eq. (45), is a time integral and therefore cannot decrease; the second is proportional to the current fibril volume and decreases when fibrils are cleared faster than they are deposited. The weighting factor *ϖ* is estimated by assuming that nephrotoxicity from AA oligomers and physical damage from fibril deposition contribute approximately equally to total renal injury in AA amyloidosis. Under conditions of impaired protein degradation with [*H*]_*tot*_ = 15 µM, the maximum accumulated oligomer toxicity is 2.08 × 10^9^ µM s (Fig. S6b). Under the same conditions, fibrillar amyloid occupies 14.9% of the total kidney volume (Fig. S5b). Setting the weighting factor such that these two factors contribute equally to Ξ_*comb*_ yields:

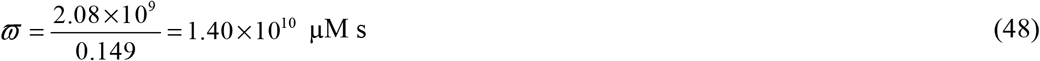

A previous study (Dormann and Lemke, 2024) proposed aging biomarkers based on the aggregation kinetics of intrinsically disordered proteins, including Aβ, tau, α-synuclein, and TDP-43. The present study applies this framework to AA amyloidosis, with important distinctions: aggregation occurs primarily in kidneys rather than brain tissue, and SAA must undergo proteolytic cleavage to generate aggregation-competent AA fragments. Accordingly, the renal biological-age index (or simply biological age) is defined as:

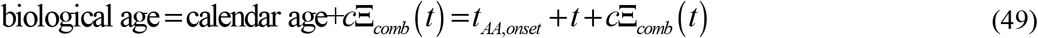

Because the fibrillar contribution to Eq. (47) decreases when deposits are cleared faster than they are formed, Eq. (49) allows the biological-age index to decrease during successful therapy. This is the intended behavior: in a prospective study, amyloidotic organ function stabilized or improved in 39 of the 42 patients whose median SAA remained below 10 mg/L (Gillmore et al., 2001), and an index that could not fall would be unable to represent that recovery. The decrease is nevertheless bounded. The accumulated toxicity Ξ of Eq. (45) is the time integral of a non-negative oligomer concentration and therefore cannot decrease, and the fibrillar term of Eq. (47) cannot be negative. The biological-age index therefore satisfies:

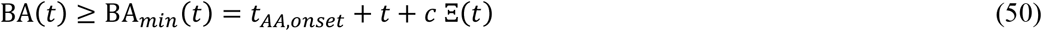

Eq. (50) follows from Eqs. (45), (47), and (49) rather than being imposed on them. The floor contains only chronological time and the accumulated oligomer toxicity, so oligomer-mediated injury is treated as permanent, whereas the part of the index above the floor, *c* times the fibrillar term of Eq. (47), is recovered as deposits are cleared. How far and how fast the index falls toward the floor after SAA is controlled is therefore set by the fibril clearance kinetics, not by the definition of the index. Whenever circulating SAA does not fall, Ξ_*comb*_ is non-decreasing and the index rises at least as fast as chronological age; this is the case for the simulations in Figs. 2 and 4–7 and Figs. S1–S26. When fibrils are resorbed, as between flares in Fig. 9e, after SAA suppression in Fig. 10a, and after the early inflammatory episode in Fig. S27b, the index rises more slowly than chronological age and, if clearance is fast enough, decreases.

**Fig. 1.**
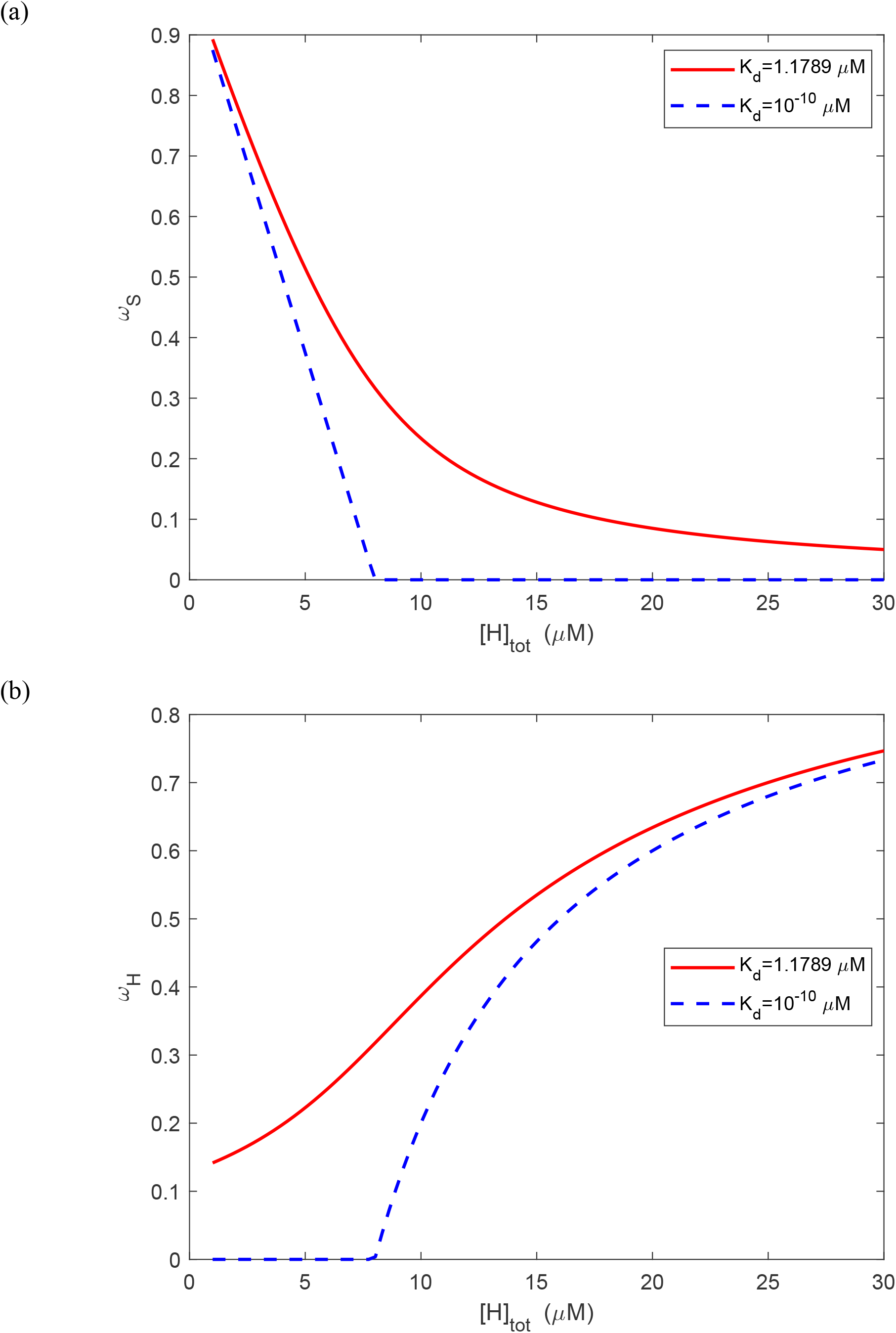
(a) Fraction of lipid-free SAA, *ω*_*S*_ . (b) Fraction of SAA-binding sites on HDL that are unoccupied, *ω*_*H*_ . The cases correspond to the baseline value of the equilibrium dissociation constant, *K*_*d*_ = 1.1789 *µ*M, and to a very small value, *K*_*d*_ = 10^−10^ *µ*M . The second case is an idealized tight-binding limit retained only to expose the stoichiometric structure of Eq. (9); it is not a physiologically attainable value. In that limit, the break in panel (a) occurs at the point of stoichiometric equivalence between HDL binding sites and total SAA.

**Fig. 2.**
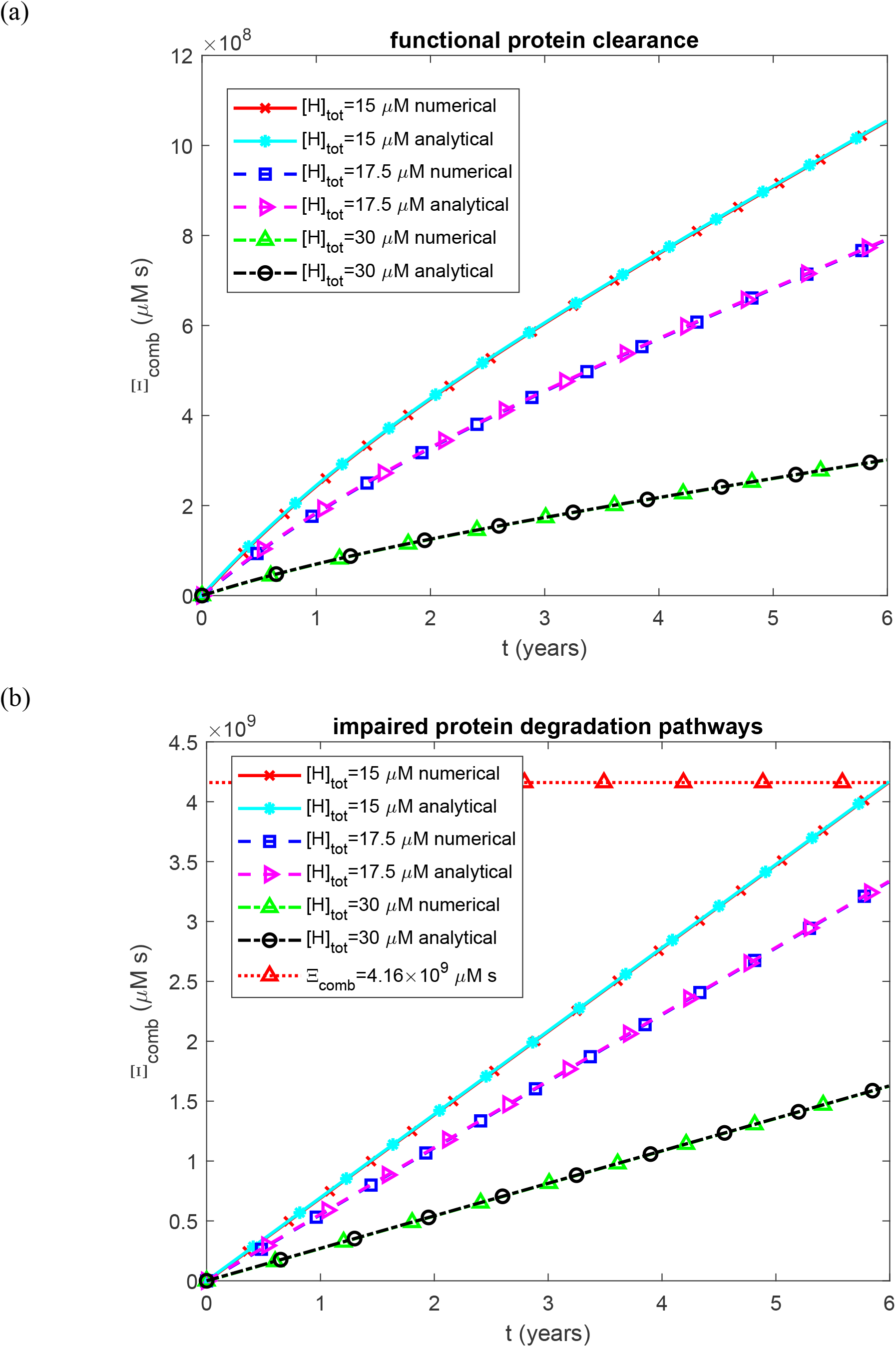
Combined renal damage from accumulated AA oligomer cytotoxicity and insoluble fibril burden as a function of time. (a) Biologically relevant half-lives *T*_1/2, *A*_, *T*_1/2,*B*_, *T*_1/2,*I*_, and *T*_1/2,*S*_, representing functional protein degradation machinery. (b) Infinite half-lives *T*_1/2, *A*_, *T*_1/2,*B*_, *T*_1/2,*I*_, and *T*_1/2,*S*_, representing complete impairment of protein degradation. Each panel shows results for three values of the effective concentration of SAA-binding sites on HDL in plasma. The horizontal line in (b) marks Ξ_*comb*_ = 4.16 × 10^9^ µM s, the combined damage that corresponds to the calibrated terminal-damage threshold of BA = 100 years (Section 2.3). Baseline parameters: *k*_1_ =10^−4^ s^−1^, *k*_2_ = 10^−4^ µM^−1^ s^−1^, *k*_*cl*_ = 9.65×10^−5^ s^−1^, *K*_*d*_ = 1.1789 *µ*M, *h*_*S*_ = 5.55×10^−7^ m s^−1^, *S*_*tot*_ = 8 *µ*M, and *θ*_1/2,*B*_ = 8.64 ×10^4^ s; remaining parameters as listed in Table 2.

**Fig. 3.**
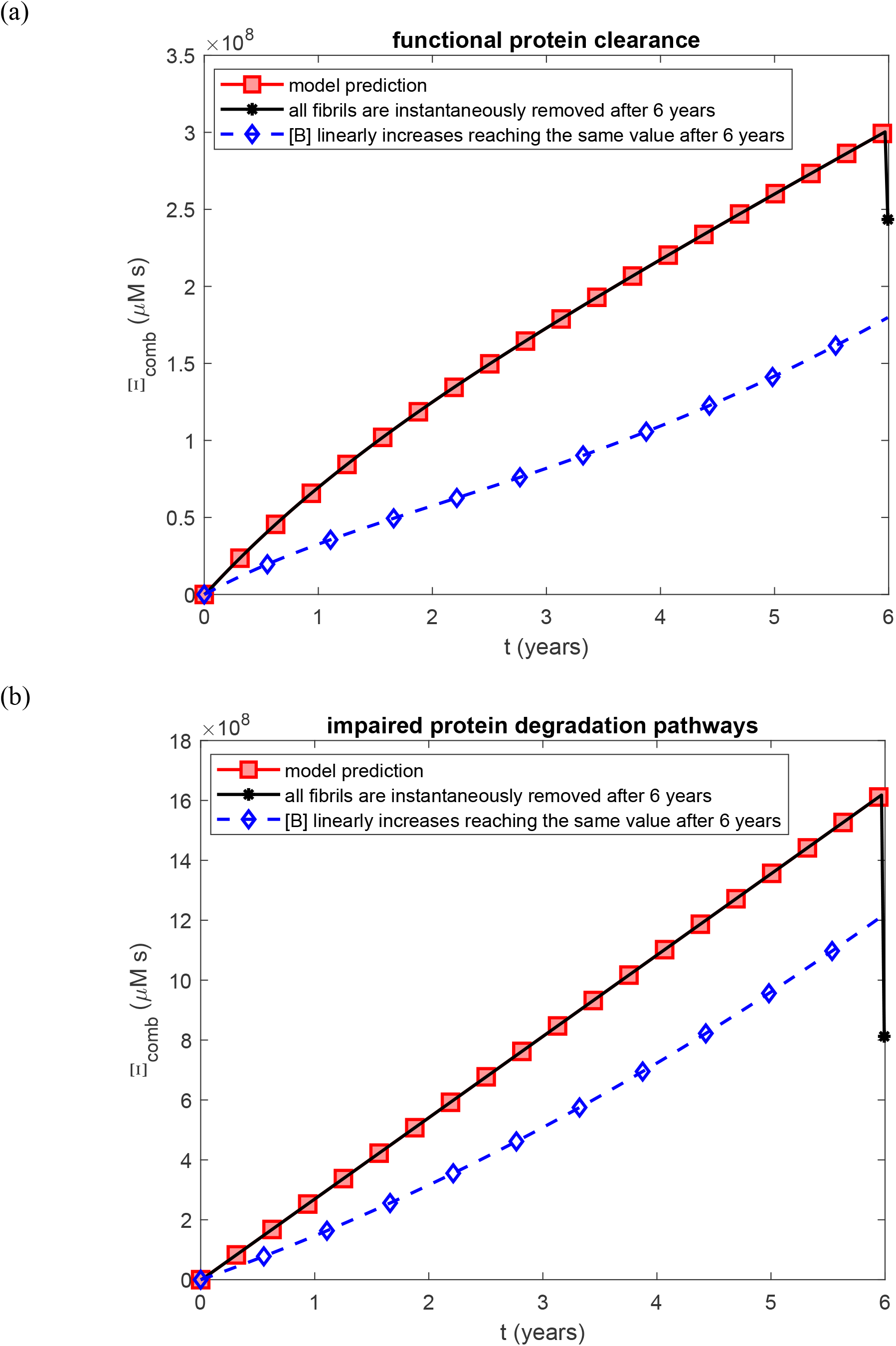
Combined renal damage from accumulated AA oligomer cytotoxicity and insoluble fibril burden as a function of time. Three scenarios are compared: the model prediction, in which [*B*] rapidly attains a constant steady-state value early in the disease course; a modified scenario following model predictions for [*B*] but with instantaneous removal of all fibrillar deposits 6 years after onset; and a hypothetical scenario in which [*B*] increases linearly throughout disease progression (see Fig. S7). (a) Biologically relevant half-lives *T*_1/2, *A*_, *T*_1/2,*B*_, *T*_1/2,*I*_, and *T*_1/2,*S*_, representing functional protein degradation machinery. (b) Infinite half-lives *T*_1/2, *A*_, *T*_1/2,*B*_, *T*_1/2,*I*_, and *T*_1/2,*S*_, representing complete impairment of protein degradation. Baseline parameters: *k*_1_ =10^−4^ s^−1^, *k*_2_ = 10^−4^ µM^−1^ s^−1^, *k*_*cl*_ = 9.65×10^−5^ s^−1^, *K*_*d*_ = 1.1789 *µ*M, *h*_*S*_ = 5.55×10^−7^ m s^−1^, *H*_*tot*_ = 30 *µ*M, *S*_*tot*_ = 8 *µ*M, and *θ*_1/2,*B*_ = 8.64 ×10^4^ s; remaining parameters as listed in Table 2.

**Fig. 4.**
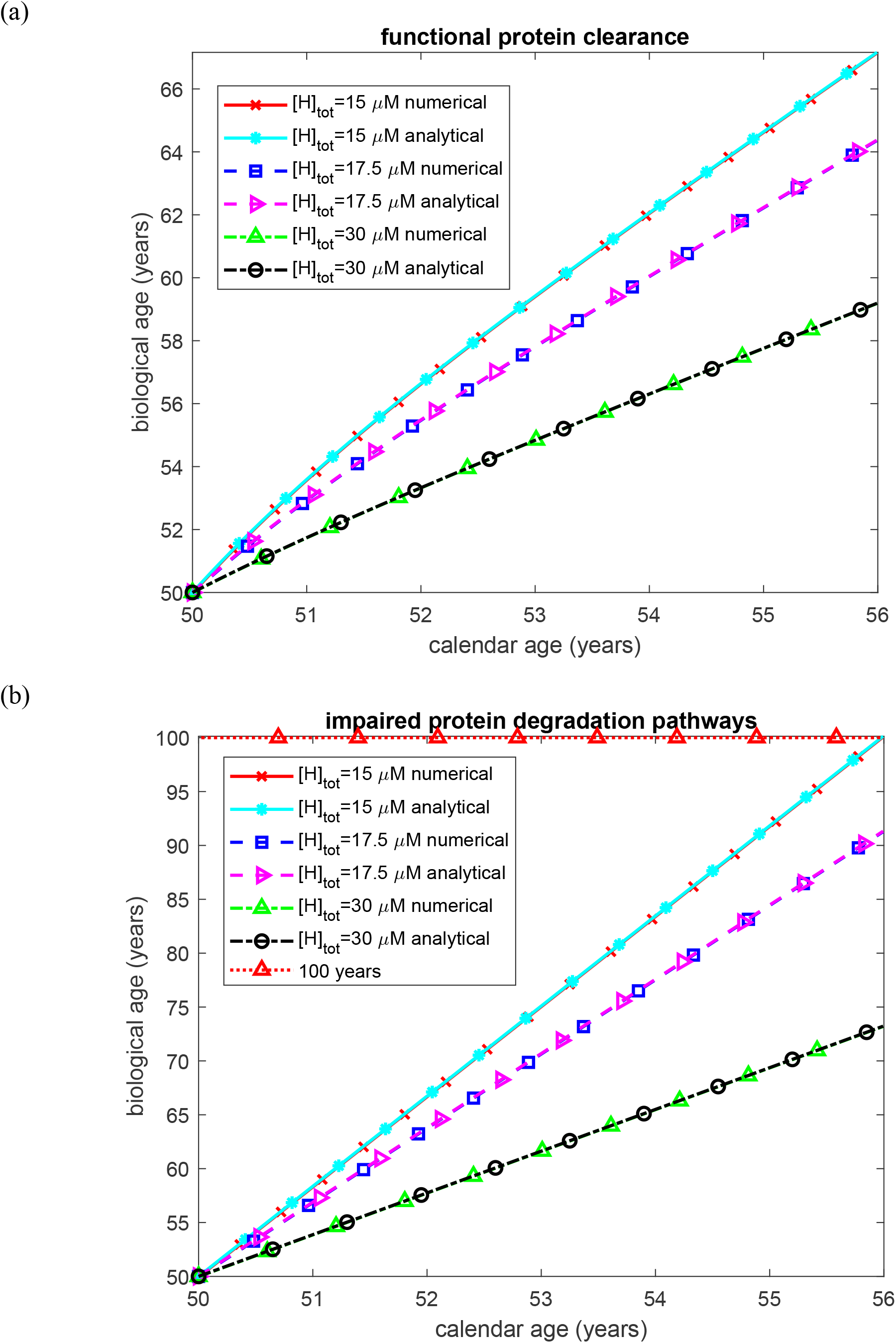
Biological age of the kidney as a function of time. (a) Biologically relevant half-lives *T*_1/2, *A*_, *T*_1/2,*B*_, *T*_1/2,*I*_, and *T*_1/2,*S*_, representing functional protein degradation machinery. (b) Infinite half-lives *T*_1/2, *A*_, *T*_1/2,*B*_, *T*_1/2,*I*_, and *T*_1/2,*S*_, representing complete impairment of protein degradation. Each panel shows results for three values of the effective concentration of SAA-binding sites on HDL in plasma. The horizontal line in (b) marks the calibrated terminal-damage threshold of BA = 100 years. Baseline parameters: *k*_1_ =10^−4^ s^−1^, *k*_2_ = 10^−4^ µM^−1^ s^−1^, *k*_*cl*_ = 9.65×10^−5^ s^−1^, *K*_*d*_ = 1.1789 *µ*M, *h*_*S*_ = 5.55×10^−7^ m s^−1^, *S*_*tot*_ = 8 *µ*M, and *θ*_1/2,*B*_ = 8.64 ×10^4^ s; remaining parameters as listed in Table 2.

**Fig. 5.**
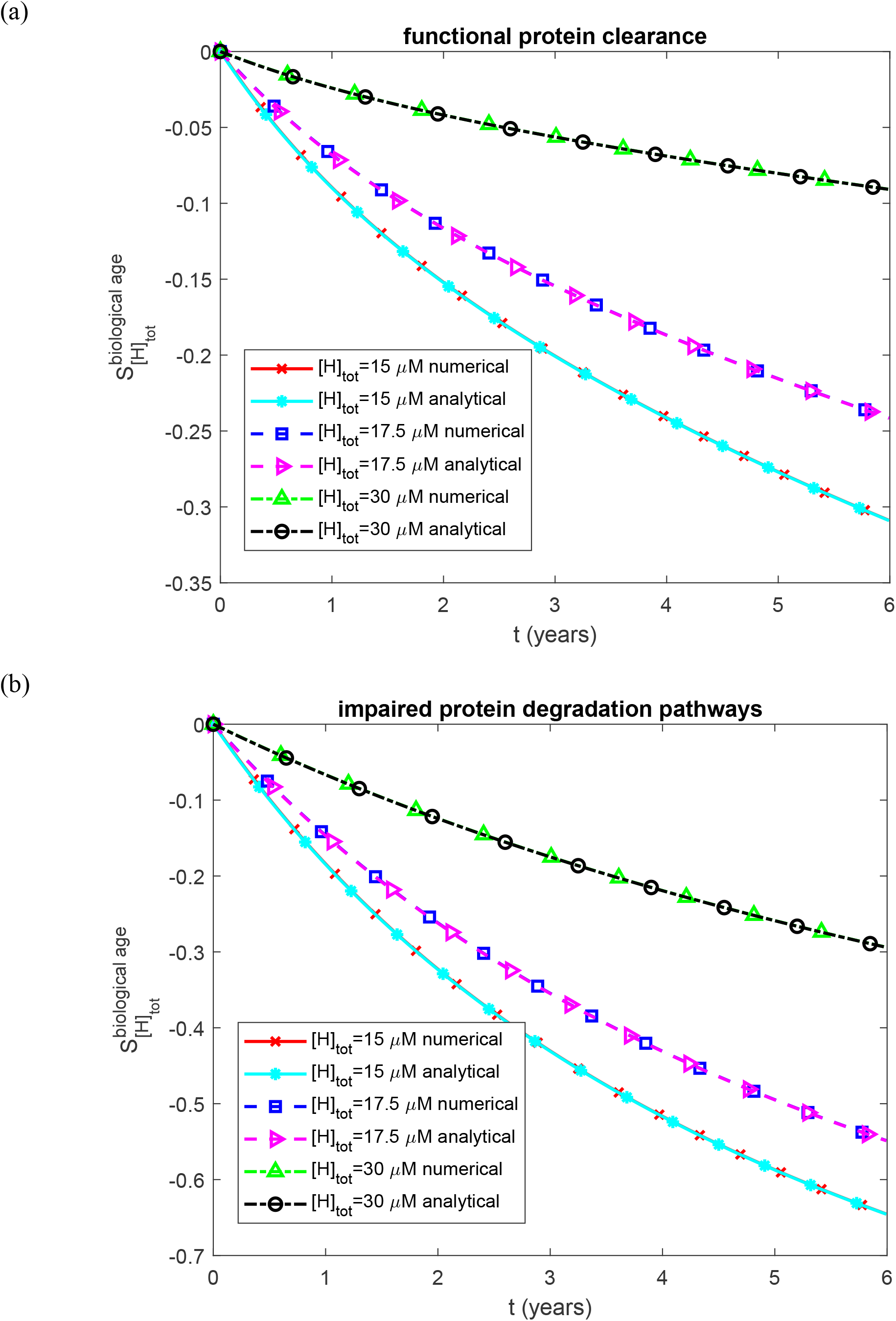
Dimensionless sensitivity of renal biological age to the effective concentration of SAA-binding sites on HDL in plasma, as a function of time. (a) Biologically relevant half-lives *T*_1/2, *A*_, *T*_1/2,*B*_, *T*_1/2,*I*_, and *T*_1/2,*S*_, representing functional protein degradation machinery. (b) Infinite half-lives *T*_1/2, *A*_, *T*_1/2,*B*_, *T*_1/2,*I*_, and *T*_1/2,*S*_, representing complete impairment of protein degradation. Each panel shows results for three values of the effective concentration of SAA-binding sites on HDL in plasma. Baseline parameters: *k*_1_ =10^−4^ s^−1^, *k*_2_ = 10^−4^ µM^−1^ s^−1^, *k*_*cl*_ = 9.65×10^−5^ s^−1^, *K*_*d*_ = 1.1789 *µ*M, *h*_*S*_ = 5.55×10^−7^ m s^−1^, *S*_*tot*_ = 8 *µ*M, and *θ*_1/2,*B*_ = 8.64 ×10^4^ s; remaining parameters as listed in Table 2.

**Fig. 6.**
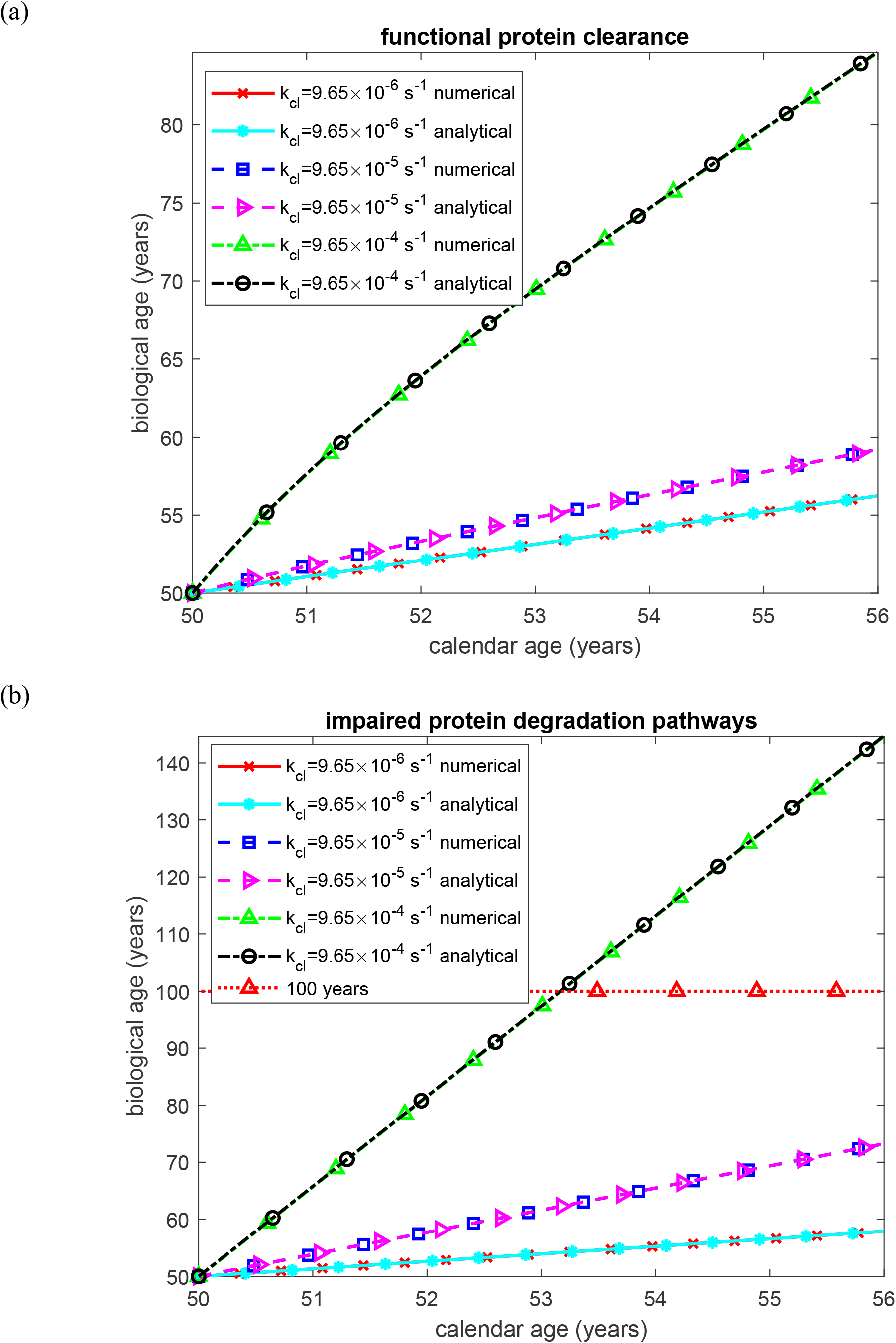
Biological age of the kidney as a function of time. (a) Biologically relevant half-lives *T*_1/2, *A*_, *T*_1/2,*B*_, *T*_1/2,*I*_, and *T*_1/2,*S*_, representing functional protein degradation machinery. (b) Infinite half-lives *T*_1/2, *A*_, *T*_1/2,*B*_, *T*_1/2,*I*_, and *T*_1/2,*S*_, representing complete impairment of protein degradation. Each panel shows results for three values of the rate constant for proteolytic cleavage of SAA that generates AA monomers. The horizontal line in (b) marks the calibrated terminal-damage threshold of BA = 100 years. Baseline parameters: *k*_1_ =10^−4^ s^−1^, *k*_2_ = 10^−4^ µM^−1^ s^−1^, *K*_*d*_ = 1.1789 *µ*M, *h*_*S*_ = 5.55×10^−7^ m s^−1^, *H*_*tot*_ = 30 *µ*M, *S*_*tot*_ = 8 *µ*M, and *θ*_1/2,*B*_ = 8.64 ×10^4^ s; remaining parameters as listed in Table 2.

Simulations using the parameter values *h*_*S*_ = 5.55×10^−7^ m s^−1^, *k*_1_ = 10^−4^ s^−1^, *k*_2_ = 10^−4^ µM^−1^ s^−1^, *k*_*cl*_ = 9.65×10^−5^ s^−1^, *T*_1/2,*B*_ = 10^14^ s, *T*_1/2,*S*_ = 10^14^ s, *T*_1/2,*A*_ = 10^14^ s, *T*_1/2,*I*_ = 10^14^ s, *θ*_1/2,*B*_ = 8.64 ×10^4^ s, *K*_*d*_ = 1.1789 *µ*M, *S*_*tot*_ = 8 *µ*M, [*H*]_*tot*_ = 15 µM, and the remaining values from Table 2 yielded a combined damage criterion of Ξ_*comb*_ (*t*_*AA, f*_) = 4.16 ×10^9^ µM s. For a clinical scenario with disease onset at age 50, progression over 6 years, and a reference terminal-damage threshold (biological age = 100 years) at age 56, parameter *c* in Eq. (49) becomes:

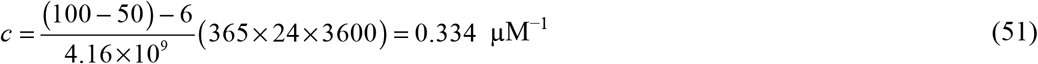

Eq. (51) determines *c* by back-solving the biological-age mapping from a single reference scenario; *c* is therefore a calibration constant, not an independently measured physiological parameter. The reference scenario maps a six-year untreated disease course beginning at age 50 onto BA = 100 years. The six-year interval is motivated by the reported median age at diagnosis and the reported 4–8-year untreated survival range, but survival time is not itself a direct measurement of time to renal failure. Accordingly, BA = 100 years should be read only as the calibrated terminal-damage threshold of this model. As shown in Fig. S28, varying the assumed reference interval from 4 to 8 years changes *c* from 0.522 to 0.238 µM^−1^, a factor of 2.2. The magnitudes of the dimensionless sensitivity coefficients change with this calibration, while their signs and the order of the parameters from most to least influential remain unchanged over the tested range. Thus the ranking of influential parameters is robust within this calibration analysis, whereas absolute values expressed in ‘years of biological age’ are not independently predicted.

### 2.4. Sensitivity analysis of renal biological age

The sensitivity of renal biological age to the model parameters was assessed using local sensitivity coefficients, defined as the first-order partial derivatives of biological age with respect to individual parameters (Beck and Arnold, 1977), (Zadeh and Montas, 2010), (Zi, 2011), (Kuznetsov and Kuznetsov, 2019). For example, the sensitivity of biological age to the effective concentration of SAA-binding sites on HDL in plasma, [*H*]_*tot*_, was computed via finite differences:

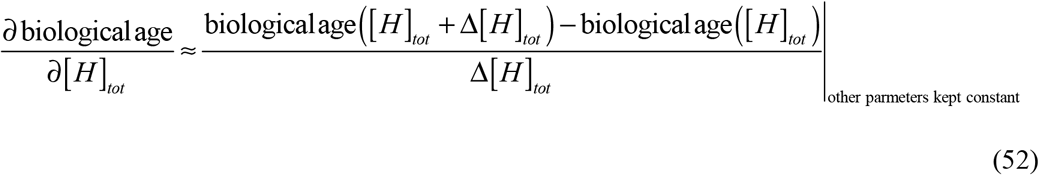

where Δ[*H*]_*tot*_ = 10^−3^ [*H*]_*tot*_ denotes the step size for numerical differentiation. Multiple step sizes were tested to verify that the calculated sensitivity coefficients were insensitive to the choice of step size.

Dimensionless relative sensitivity coefficients were additionally computed to enable direct comparison across parameters differing in units and magnitude (Zadeh and Montas, 2010), (Kuznetsov and Kuznetsov, 2019):

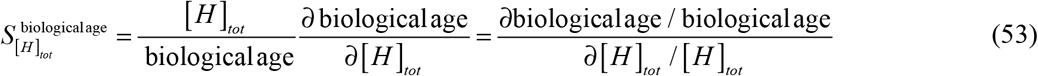

The dimensionless coefficient 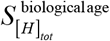 represents the fractional change in biological age per unit fractional change in [*H*]_*tot*_, such that a value of 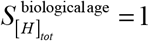 indicates that a 1% increase in [*H*]_*tot*_ produces a 1% increase in biological age.

### 2.5. Dimensionless formulation of the aggregation step and the autocatalytic threshold

The Introduction noted the expectation that a threshold exists at which autocatalytic conversion becomes self-sustaining (Knowles et al., 2009). The steady-state equations derived above make that threshold explicit. Let [*B*]* denote the supply-limited oligomer concentration, that is, the concentration that would be reached if every AA monomer generated by cleavage were converted into oligomers:

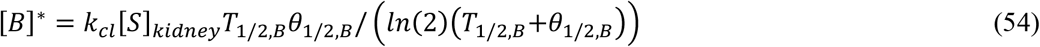

We also introduce the two dimensionless groups:

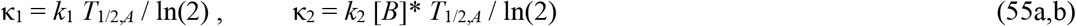

With *b* = [*B*]/[*B*]* the normalized oligomer burden, Eqs. (21) and (22) collapse to the single quadratic equation

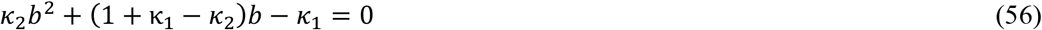

whose physically meaningful root is

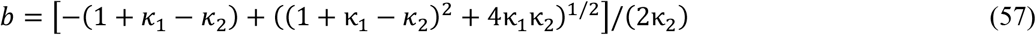

The group κ_1_ compares the rate of primary nucleation with the rate of proteolytic degradation of AA monomers, while κ_2_ compares the rate of autocatalytic conversion evaluated at the supply-limited oligomer concentration [*B*]* with that same degradation rate; κ_2_ is therefore an autocatalytic gain. In the limit of vanishing primary nucleation (κ_1_ → 0), Eq. (57) reduces to

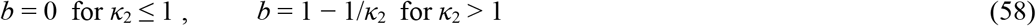

At κ_1_ = 0, Eq. (56) has two steady-state branches that cross at κ_2_ = 1: *b* = 0, which exists for every κ_2_, and *b* = 1 − 1/κ_2_, which is non-negative only for κ_2_ ≥ 1. Thus κ_2_ = 1 marks an autocatalytic threshold separating a zero-oligomer branch from a positive branch that approaches the supply limit as κ_2_ increases. The stability of the two branches follows from linearizing Eqs. (19) and (20) at fixed renal SAA concentration, which is legitimate because Eq. (15) contains neither [*A*] nor [*B*]. For κ_1_ = 0, the Jacobian on the zero-oligomer branch is triangular, with eigenvalues −ln(2)/*T*_1/2,*A*_ and *λ*_*B*_(κ_2_ − 1), where *λ*_*B*_ = ln(2)/*T*_1/2,*B*_ + ln(2)/*θ*_1/2,*B*_ is the total first-order loss rate of free oligomers. The zero-oligomer branch is therefore stable for κ_2_ < 1 and unstable for κ_2_ > 1. On the positive branch, which is physical only for κ_2_ > 1, the Jacobian has negative trace and positive determinant, so that branch is stable wherever it is physical. κ_2_ = 1 is thus a transcritical bifurcation of the monomer–oligomer subsystem, at which the two branches exchange stability. The oligomer burden nevertheless changes continuously at the threshold, rising from zero as *b* = 1 − 1/κ_2_ (*b* = 0.09 at κ_2_ = 1.1 and 0.5 at κ_2_ = 2); what changes abruptly is whether an oligomer population can sustain itself without primary nucleation. For κ_1_ > 0 the non-negative steady state is unique and, by the same argument, linearly stable for every κ_2_, so primary nucleation unfolds the bifurcation into a continuous crossover; these eigenvalues are verified numerically in AA_tipping_point.m (Kuznetsov, 2026b). The sharpness of the transition is set by κ_1_. At the baseline nucleation rate constant, *k*_1_ = 10^−4^ s^−1^, κ_1_ = 0.52 under functional clearance and the transition is strongly smoothed; a switch-like transition requires κ_1_ ≪ 1, which within the range considered in Table 2 is reached for *k*_1_ = 10^−6^ and 10^−8^ s^−1^ (κ_1_ ≈ 5 × 10^−3^ and 5 × 10^−5^). Because [*B*]* is proportional to renal SAA concentration through Eq. (16), and hence to the free plasma SAA concentration set by HDL buffering through Eq. (9), κ_2_ depends on the effective HDL binding-site concentration; the condition κ_2_ = 1 therefore defines a critical binding-site concentration for each value of *k*_2_. Eqs. (54)–(57) are an exact restatement of the steady-state system rather than an additional approximation, and they reproduce the numerical solution of Eqs. (15), (19), and (20) to a relative error of order 10^−15^ (Fig. 8b).

## 3. Results

The results are presented in four parts: the SAA–HDL binding equilibrium (Fig. 1); the steady-state course of the renal AA species, combined damage, and biological age under constant inflammation (Figs. 2–4 and S1–S7); the dependence of biological age on, and its sensitivity to, the model parameters (Figs. 5–7, S8–S26, and Table 3); and the autocatalytic threshold, time-varying inflammation, treatment timing, and comparison with clinical data (Figs. 8–11, S27, and S28). Because the effective concentration of SAA-binding sites on HDL is the principal disease variable in the first two parts, the direction and approximate magnitude of inflammatory HDL depletion are summarized first.

**Table 3.** Dimensionless sensitivity of biological age to model parameters evaluated 6 years after disease onset. Simulations employed the baseline values *k*_1_ = 10^−4^ s^−1^, *k*_2_ = 10^−4^ µM^−1^ s^−1^, *k*_*cl*_ = 9.65 × 10^−5^ s^−1^, *K*_*d*_ = 1.1789 µM, *h*_*S*_ = 5.55 × 10^−7^ m s^−1^, [*H*]_*tot*_ = 30 µM, *S*_*tot*_ = 8 µM, and *θ*_1/2,*B*_ = 8.64 × 10^4^ s, with remaining parameters taken from Table 2. Within each descriptive group, parameters are listed in descending order of the magnitude of their baseline sensitivity coefficients. The labels ‘largest’, ‘intermediate’, and ‘smallest’ are rankings for this baseline calculation rather than universal sensitivity thresholds; Section 2.5 and Fig. 8 show that the ranking of aggregation parameters can change when the model moves to a different dynamical regime.

| Parameter | Sensitivity for the case of functional protein clearance | Sensitivity for the case of impaired protein degradation pathways |
|---|---|---|
| Parameters with the largest baseline sensitivities |  |  |
| [*S*]<sub>*tot*</sub> | 0.0958 | 0.310 |
| [*H*]<sub>*tot*</sub> | −0.0908 | −0.294 |
| Parameters with intermediate baseline sensitivities |  |  |
| *K*<sub>*d*</sub> | 0.0679 | 0.220 |
| *k*<sub>*cl*</sub> | 0.0664 | 0.211 |
| *θ*<sub>1/2,*B*</sub> | 0.0381 | 0.118 |
| Parameters with the smallest baseline sensitivities |  |  |
| *h*<sub>*S*</sub> | 0.0189 | 0.0246 |
| *k*<sub>2</sub> | 0.0187 | 1.90 × 10<sup>−6</sup> |
| *k*<sub>1</sub> | 0.0145 | 4.42 × 10<sup>−7</sup> |

In human studies, HDL cholesterol levels typically decrease by approximately twofold during strong inflammatory acute-phase responses, while in chronic inflammatory diseases HDL is more often reduced by roughly 1.3–1.5-fold compared with healthy controls (Jahangiri et al., 2009).

A small equilibrium dissociation constant (*K*_*d*_) indicates strong binding between SAA and HDL, whereas a large *K*_*d*_ value corresponds to relatively weak binding. Fig. 1 illustrates these effects. For the relatively large baseline *K*_*d*_ value (1.1789 *µ*M), which represents weaker binding, the fraction of lipid-free SAA, *ω*_*S*_, decreases monotonically as the effective SAA-binding-site concentration increases (Fig. 1a). Conversely, the fraction of unoccupied HDL-associated binding sites, *ω*_*H*_, increases with the effective SAA-binding-site concentration (Fig. 1b).

In contrast, for the very small *K*_*d*_ value of 10^−10^ *µ*M (corresponding to tight binding), the fraction of lipid-free SAA decreases linearly until the effective SAA-binding-site concentration reaches a threshold of 8 *µ*M . This threshold corresponds to the SAA concentration of 8 *µ* M used in the simulation. Once the effective SAA-binding-site concentration exceeds 8 *µ*M, the free SAA fraction remains at zero, as all SAA molecules are sequestered by HDL (Fig. 1a). This second case is an idealized tight-binding limit introduced only to expose the stoichiometric structure of Eq. (9), and not a physiologically attainable affinity: the dissociation constants measured for SAA binding to HDL by surface plasmon resonance are 3.7–18 µM (Yamada et al., 2009). The apparent sharp transition in this limit is therefore not a dynamical tipping point of the disease; it is simply the point of stoichiometric equivalence at which the number of HDL binding sites equals the total SAA concentration. A genuine dynamical threshold does exist in the model, but it arises in the aggregation step rather than in the binding step, and it is analyzed in Section 2.5 and Fig. 8.

Figs. S1–S3 demonstrate that the molar concentrations of SAA, AA monomers, and free AA oligomers in the kidneys ([*S*]_*kidney*_, [*A*], and [*B*], respectively) rapidly attain constant steady-state values. This validates the steady-state approximation underlying the analytical solutions given by Eqs. (16), (24), and (25). Furthermore, all three concentrations increase as the effective HDL binding-site concentration decreases, in agreement with published evidence showing that inflammatory depletion of HDL reduces available SAA binding capacity, thereby enabling liver-derived SAA to saturate HDL particles and generate an excess of free, lipid-poor SAA in the bloodstream (Webb, 2021), (Schuchardt et al., 2019). Lipid-poor SAA promotes renal inflammation and can participate in amyloid formation (Simons et al., 2013).

The molar concentration of AA oligomers incorporated into amyloid fibrils in the kidneys, [*I*], increases over time. For physiologically relevant half-lives of AA species, the rate of fibril accumulation progressively decreases, and the fibril burden approaches the steady-state burden predicted by Eq. (28) (Fig. S4a). By contrast, when fibril degradation is absent (infinite half-life scenario), fibril burden increases linearly and indefinitely (Fig. S4b). A reduction in the effective HDL binding-site concentration leads to greater fibril accumulation in the kidneys. The fraction of kidney volume occupied by fibrils displays similar behavior, asymptotically approaching a steady-state value in Fig. S5a. A lower effective HDL binding-site concentration likewise drives increased fibril accumulation within the renal parenchyma (Fig. S5a,b).

Accumulated AA oligomer nephrotoxicity increases linearly with time under both finite- and infinite-half-life scenarios for AA species (Fig. S6a,b). This linear increase follows directly from the definition of accumulated nephrotoxicity as the time integral of free oligomer concentration [*B*] (Eq. (45)): since [*B*] rapidly attains and maintains a constant steady-state value for most of the disease course (Fig. S3), its time integral increases linearly. Furthermore, a reduced effective HDL binding-site concentration leads to greater accumulated nephrotoxicity (Fig. S6a,b).

The combined nephrotoxic injury in AA amyloidosis, Ξ_*comb*_, which combines two mechanisms of renal damage—accumulated cytotoxicity of AA oligomers and aggregation of insoluble fibrils—increases with time. This increase slows down over time when biologically relevant half-lives of AA species are used (Fig. 2a) and remains linear when the half-lives are infinite (no degradation of AA species) (Fig. 2b). The combined nephrotoxic injury increases as the effective HDL binding-site concentration decreases.

The combined nephrotoxic damage criterion Ξ_*comb*_ cannot be returned to zero by fibril clearance alone. Even if a treatment capable of clearing all fibrillar deposits, such as fibril-specific monoclonal antibodies (Thorne et al., 2022), were developed and administered 6 years after onset, it would not return Ξ_*comb*_ to zero. This residual arises because Ξ_*comb*_ comprises two contributions: fibril burden, which is amenable to therapeutic clearance, and accumulated oligomer cytotoxicity, which integrates continuously over time (Eq. (45)) and so cannot decrease. As Fig. 3 shows (black line with star symbols), such therapy would reduce Ξ_*comb*_ only by its fibrillar part. The existence of this residual follows directly from defining the oligomer contribution as a time integral (Eq. (45)) and is not an independent finding about renal injury; what the model adds is the size of the residual relative to the fibrillar term, which depends on the binding and aggregation dynamics. The instantaneous criterion therefore does fall when fibrils are removed, and the biological-age index of Eq. (49) falls with it, although never below the floor of Eq. (50).

The combined nephrotoxic injury is also path-dependent rather than state-dependent. In a hypothetical scenario in which [*B*] increases linearly throughout disease progression (blue diamond line, Fig. S7), Ξ_*comb*_ attains a lower value at 6 years than predicted by the model, where [*B*] maintains a constant steady-state value for most of the disease course (red square line, Fig. S7). This occurs even though both *V*_*fibrils*_ (*t*) scenarios produce identical values of [*B*] and 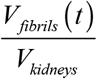 at *t* = 6 years, demonstrating that the history of *V*_*kidneys*_ oligomer exposure, not merely its current level, determines the extent of nephrotoxic damage.

Renal biological age at various effective HDL binding-site concentrations (Fig. 4) exhibits behavior analogous to that of Ξ_*comb*_ in Fig. 2. This similarity follows directly from the assumed linear relation between biological age and Ξ_*comb*_ (Eq. (49)). Biological age increases more rapidly when impaired protein degradation pathways are assumed (Fig. 4b) and increases as the effective HDL binding-site concentration decreases.

The sensitivity of biological age to effective HDL binding-site concentration is negative (Fig. 5), indicating that biological age decreases with increasing effective HDL binding-site concentration, consistent with the trends observed in Fig. 4. The magnitude of the dimensionless sensitivity decreases as the effective HDL binding-site concentration increases.

Increasing the SAA cleavage rate constant *k*_*cl*_ decreases renal SAA monomer concentration (Fig. S8) while increasing AA monomer and free oligomer concentrations (Figs. S9 and S10). A counterintuitive result emerges when comparing the half-life scenarios: although oligomer concentrations are approximately twice as high in the infinite-half-life scenario (Fig. S10b) as under physiologically relevant conditions (Fig. S10a), AA monomer concentrations are lower (Fig. S9b vs. S9a). This behavior can be explained by accelerated autocatalytic conversion of monomers to oligomers driven by the elevated oligomer concentration (Eq. (18)).

The concentration of AA oligomers incorporated into fibrillar deposits in the kidneys (Fig. S11), the fractional kidney volume occupied by fibrils (Fig. S12), and accumulated nephrotoxicity (Fig. S13) all increase with the SAA cleavage rate constant. For physiologically relevant half-lives, [*I*] and fibril volume fraction 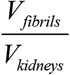 increase at a decelerating rate, whereas infinite half-lives produce linear growth (Figs. S11 and S12). In contrast, accumulated nephrotoxicity increases linearly with time under both finite- and infinite-half-life scenarios (Fig. S13). This linear behavior follows directly from the definition of accumulated nephrotoxicity as the time integral of [*B*] (Eq. (45)): since [*B*] attains and maintains a constant steady-state value for most of the disease course (Fig. S10), its integral increases linearly with time.

Combined renal damage from accumulated AA oligomer cytotoxicity and insoluble fibril burden increases with the SAA cleavage rate constant (Fig. S14), with renal biological age exhibiting the same trend (Fig. 6). For the largest cleavage rate constant considered, *k*_*cl*_ = 9.65×10^−4^ s^−1^, the calibrated terminal-damage threshold (biological age = 100 years) is reached after approximately 3.5 years of disease progression when degradation of AA species is absent (infinite half-lives; Fig. 6b).

The sensitivity of biological age to the SAA cleavage rate constant is positive (Fig. 7), indicating that biological age increases with increasing *k*_*cl*_ . The magnitude of the dimensionless sensitivity increases with the cleavage rate constant.

**Fig. 7.**
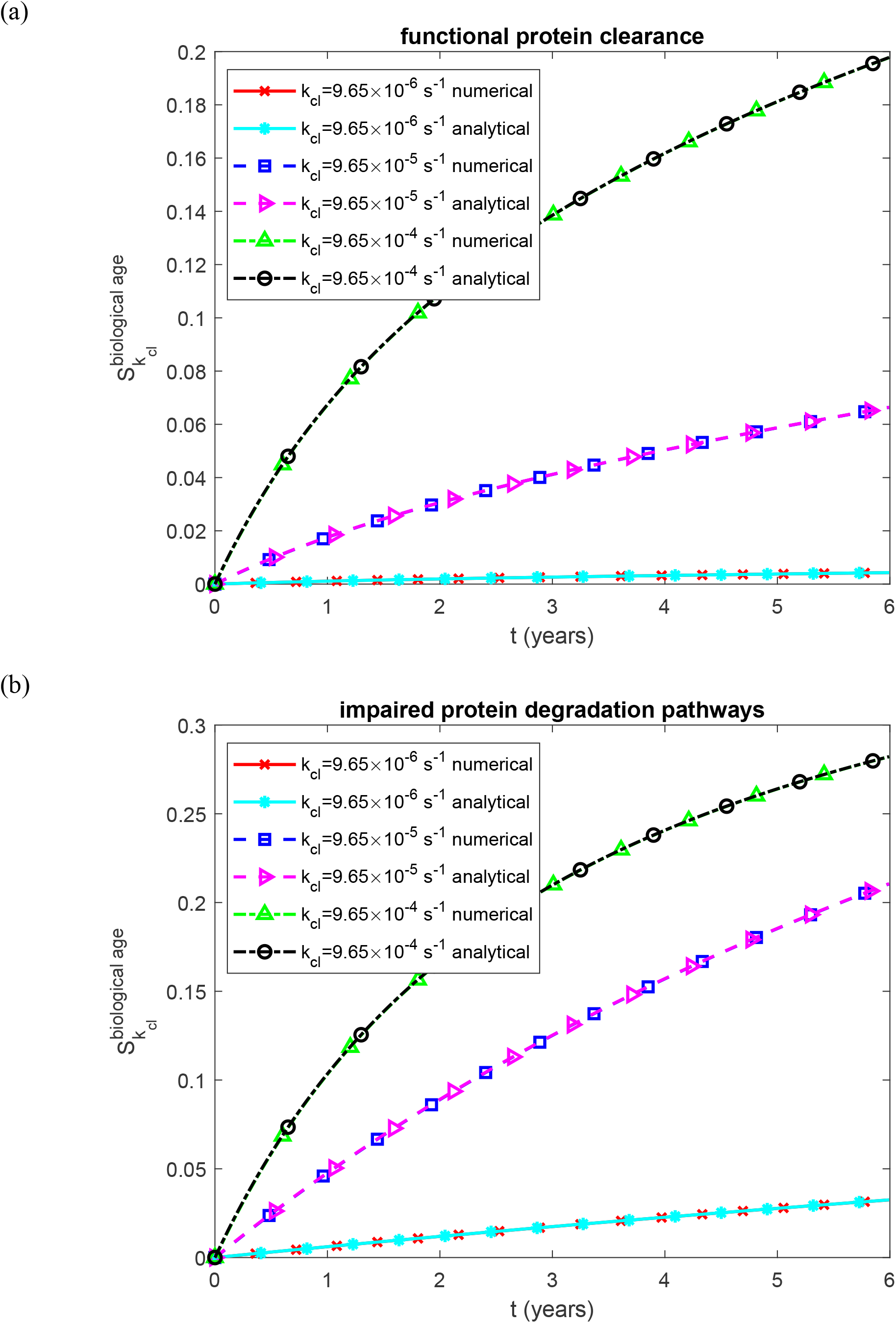
Dimensionless sensitivity of renal biological age to the rate constant for proteolytic cleavage of SAA that generates AA monomers, as a function of time. (a) Biologically relevant half-lives *T*_1/2, *A*_, *T*_1/2,*B*_, *T*_1/2,*I*_, and *T*_1/2,*S*_, representing functional protein degradation machinery. (b) Infinite half-lives *T*_1/2, *A*_, *T*_1/2,*B*_, *T*_1/2,*I*_, and *T*_1/2,*S*_, representing complete impairment of protein degradation. Each panel shows results for three values of the rate constant for proteolytic cleavage of SAA that generates AA monomers. Baseline parameters: *k*_1_ =10^−4^ s^−1^, *k*_2_ = 10^−4^ µM^−1^ s^−1^, *K*_*d*_ = 1.1789 *µ*M, *h*_*S*_ = 5.55×10^−7^ m s^−1^, *H*_*tot*_ = 30 *µ*M, *S*_*tot*_ = 8 *µ*M, and *θ*_1/2,*B*_ = 8.64 ×10^4^ s; remaining parameters as listed in Table 2.

**Fig. 8.**
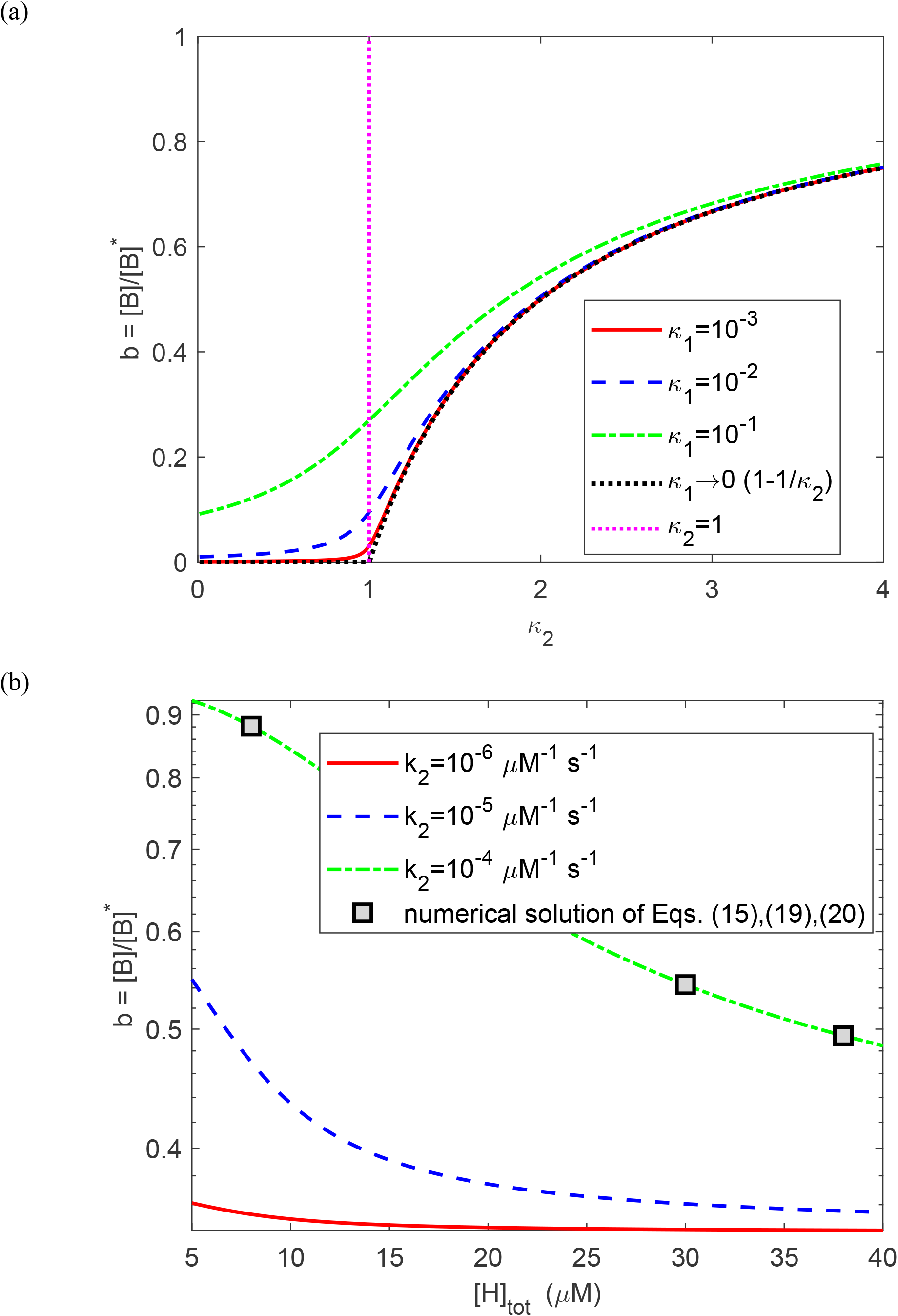
Autocatalytic threshold of the aggregation step. (a) Normalized oligomer burden *b* = [*B*]/[*B*]* as a function of the autocatalytic gain κ_2_ for three values of κ_1_, together with the limiting relation *b* = 1 − 1/κ_2_ as κ_1_ → 0. The vertical line marks the zero-primary-nucleation threshold κ_2_ = 1; finite κ_1_ smooths the transition. (b) The same quantity as a function of the effective concentration of SAA-binding sites on HDL for three values of *k*_2_ at the baseline *k*_1_ = 10^−4^ s^−1^ (κ_1_ = 0.52); symbols show the numerical solution of Eqs. (15), (19), and (20). Functional protein clearance; remaining parameters as in Table 2.

Renal biological age increases with the mass transfer coefficient *h*_*S*_ characterizing free SAA transport from plasma into the kidneys (Fig. S15). Consistent with this trend, the sensitivity of biological age to *h*_*S*_ is positive (Fig. S16). Notably, for physiologically relevant half-lives, the dimensionless sensitivity coefficient exhibits nonmonotonic behavior with respect to *h*_*S*_ : it initially increases and subsequently decreases with increasing *h*_*S*_ (Fig. S16a).

Renal biological age increases with the half-deposition time *θ*_1/2, *B*_ characterizing incorporation of free AA oligomers into fibrillar deposits (Fig. S17). Consistent with this trend, the sensitivity of biological age to *θ*_1/2, *B*_ is positive (Fig. S18). Under physiologically relevant half-life conditions, the dimensionless sensitivity magnitude exhibits nonmonotonic behavior with respect to *θ*_1/2, *B*_ : it initially increases and subsequently decreases with increasing *θ*_1/2, *B*_ (Fig. S18a).

Renal biological age increases with the equilibrium dissociation constant *K*_*d*_ characterizing SAA–HDL binding affinity (Fig. S19). This behavior reflects the inverse relationship between *K*_*d*_ and binding affinity: a small *K*_*d*_ corresponds to tight SAA–HDL association, reducing the concentration of free SAA available for renal filtration and deposition. Consistent with this trend, the sensitivity of biological age to *K*_*d*_ is positive (Fig. S20), with the sensitivity magnitude increasing monotonically with *K*_*d*_ .

Renal biological age increases with total SAA concentration (HDL-bound and free combined) (Fig. S21). For a given equilibrium dissociation constant *K*_*d*_, a higher total SAA concentration yields a disproportionately larger free SAA pool available for renal filtration and deposition, because HDL buffering saturates (Eq. (9)). Consistent with this trend, the sensitivity of biological age to total SAA concentration is positive, with its magnitude increasing monotonically with total SAA concentration (Fig. S22).

Under physiologically relevant half-life conditions of AA species, renal biological age increases with the first pseudo-elementary (nucleation) rate constant *k*_1_ of the F–W model, with this dependence becoming pronounced at larger values of *k*_1_ (Fig. S23a). Under infinite-half-life conditions, biological age exhibits negligible dependence on *k*_1_ (Fig. S23b). The sensitivity of biological age to *k*_1_ is positive under physiologically relevant conditions, increasing in magnitude with *k*_1_ (Fig. S24a). Under infinite-half-life conditions, the dimensionless sensitivity is negligibly small, on the order of 10^−6^ (Fig. S24b).

The dependence of renal biological age on the second pseudo-elementary (autocatalytic growth) rate constant *k*_2_ of the F–W model is analogous to that of *k*_1_ . Under functional protein clearance conditions, biological age increases with *k*_2_, particularly at larger values of *k* _2_ (Fig. S25a). When protein clearance is impaired, however, the dependence of biological age on *k*_2_ becomes negligible (Fig. S25b). The sensitivity of biological age to *k*_2_ is positive but small for both scenarios (Fig. S26), with the dimensionless sensitivity becoming on the order of 10^−6^ under impaired protein degradation (Fig. S26b). For both rate constants, the analytical (steady-state) sensitivity under impaired protein degradation is zero; the small numerical values arise from the initial approach of [*A*] and [*B*] to steady state, the only period in which the Finke–Watzky rate constants affect the solution when protein degradation is impaired.

The parameter sweeps and local sensitivity analysis above identify which quantities matter near the baseline operating point, but a local sensitivity ranking does not reveal whether a parameter can move the system into a different aggregation regime. Table 3 therefore provides only the first part of the picture. At baseline, the largest local sensitivities are associated with total SAA concentration and the effective concentration of SAA-binding sites on HDL. SAA–HDL affinity, the proteolytic SAA cleavage rate constant, and the oligomer half-deposition time have intermediate coefficients, whereas the mass transfer coefficient and the two Finke–Watzky rate constants have smaller coefficients. This distinction is especially important for *k*_2_: its local sensitivity is small at baseline, yet changing *k*_2_ over its plausible range can move the system across the autocatalytic threshold. This motivates the global regime analysis in Fig. 8.

Fig. 8 asks what separates nucleation-limited from supply-limited aggregation. For small κ_1_, the normalized oligomer burden *b* remains close to zero while κ_2_ < 1, then rises rapidly once κ_2_ exceeds unity and approaches the supply-limited relation *b* = 1 − 1/κ_2_ (Fig. 8a). Increasing κ_1_ smooths this transition because primary nucleation continues to generate oligomers below the autocatalytic threshold. Fig. 8b maps the same result onto the effective concentration of SAA-binding sites on HDL, [*H*]_*tot*_, a model quantity that should not be confused with a measured HDL-cholesterol concentration. At the baseline *k*_1_ = 10^−4^ s^−1^, κ_1_ = 0.52, so the transition is already smoothed. For *k*_2_ = 10^−4^ µM^−1^ s^−1^, κ_2_ = 1 is reached only at [*H*]_*tot*_ = 35.6 µM, placing the baseline system in the supply-limited regime over the range considered.

Lowering *k*_2_ to 10^−5^ µM^−1^ s^−1^ shifts the critical binding-site concentration to 6.5 µM, whereas for *k*_2_ = 10^−6^ µM^−1^ s^−1^ the system remains nucleation-limited throughout the range. The numerical solutions agree with the closed-form steady-state result to a relative error of order 10^−15^. Thus, *k*_2_ can determine which aggregation regime applies even though its local sensitivity is small at the baseline point.

Once the existence of two aggregation regimes is recognized, a natural question is whether the mean SAA concentration is sufficient to characterize inflammatory exposure. Fig. 9 shows that it is not. In these simulations, the only model input that varies in time is the total SAA concentration, [*S*]_*tot*_, which enters the model through Eq. (9) (Fig. 9a). An inflammatory flare raises [*S*]_*tot*_ from an inter-flare concentration of 0.5 µM, representing controlled disease, to a constant peak and then returns it to 0.5 µM; the onset and offset are smoothed over two days. One flare occurs each year, and the duty cycle is the fraction of the year spent in the elevated-SAA state: three months for a 25% duty cycle and about five weeks for a 10% duty cycle. Three six-year histories are constructed to have the same time-averaged total SAA concentration of 4 µM: sustained inflammation at 4 µM, annual 25%-duty-cycle flares peaking at 14.5 µM, and annual 10%-duty-cycle flares peaking at 35.5 µM. Despite their identical mean SAA, the three histories deliver very different amounts of free SAA to the kidney. Because saturation of HDL buffering makes free SAA a convex function of total SAA (Eq. (9)), free SAA falls to 0.038 µM between flares but rises to 3.4 µM and 21 µM during 25%- and 10%-duty-cycle flares, compared with a constant 0.38 µM under sustained inflammation; its six-year mean is therefore 2.2-fold and 5.2-fold higher for the two flare histories (Fig. 9b). With functional protein clearance, the time-averaged concentration of free AA oligomers in the kidney, [*B*], is 3.7-fold and 9.5-fold higher (Fig. 9c), more than the increase in free SAA, because near the autocatalytic threshold the F–W kinetics make oligomer burden a convex function of free SAA (Eq. (57)). With impaired protein degradation, time-averaged [*B*] rises 2.2-fold and 5.2-fold (Fig. 9d), in proportion to free SAA, because aggregation is then supply-limited and approximately linear in its supply; in this case the amplification comes from HDL saturation alone. These differences carry through to renal damage: six-year combined renal damage is 3.6-fold and 9.1-fold higher for the 25% and 10% duty cycles, respectively, than for sustained inflammation with functional protein clearance, and 2.2-fold and 5.2-fold higher with impaired protein degradation (Figs. 9e,f). A linear dependence of oligomer concentration on total SAA would instead give the same damage for all three histories. The model therefore predicts that the temporal pattern of SAA exposure, and not only its mean, may carry prognostic information.

**Fig. 9.**
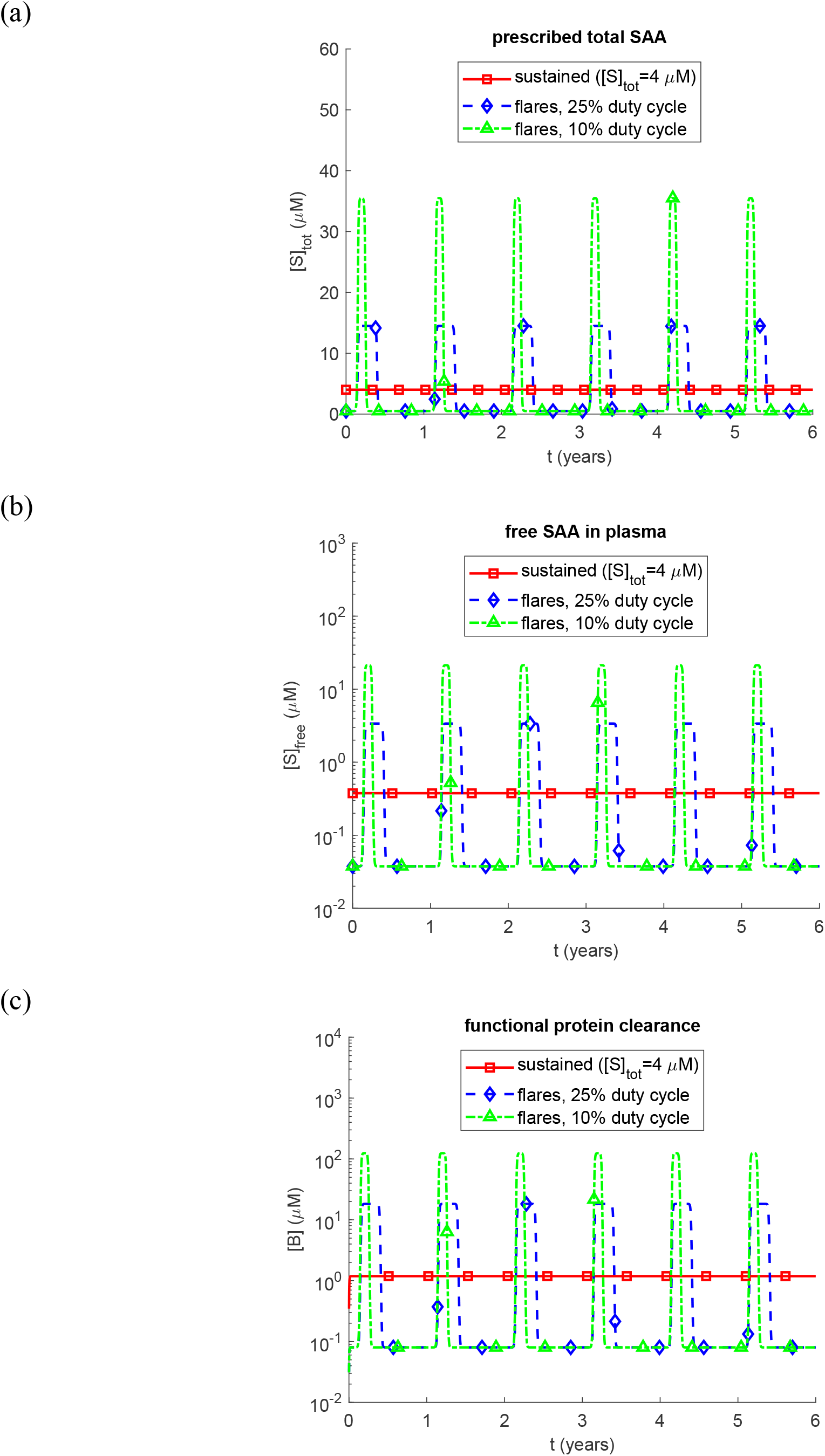
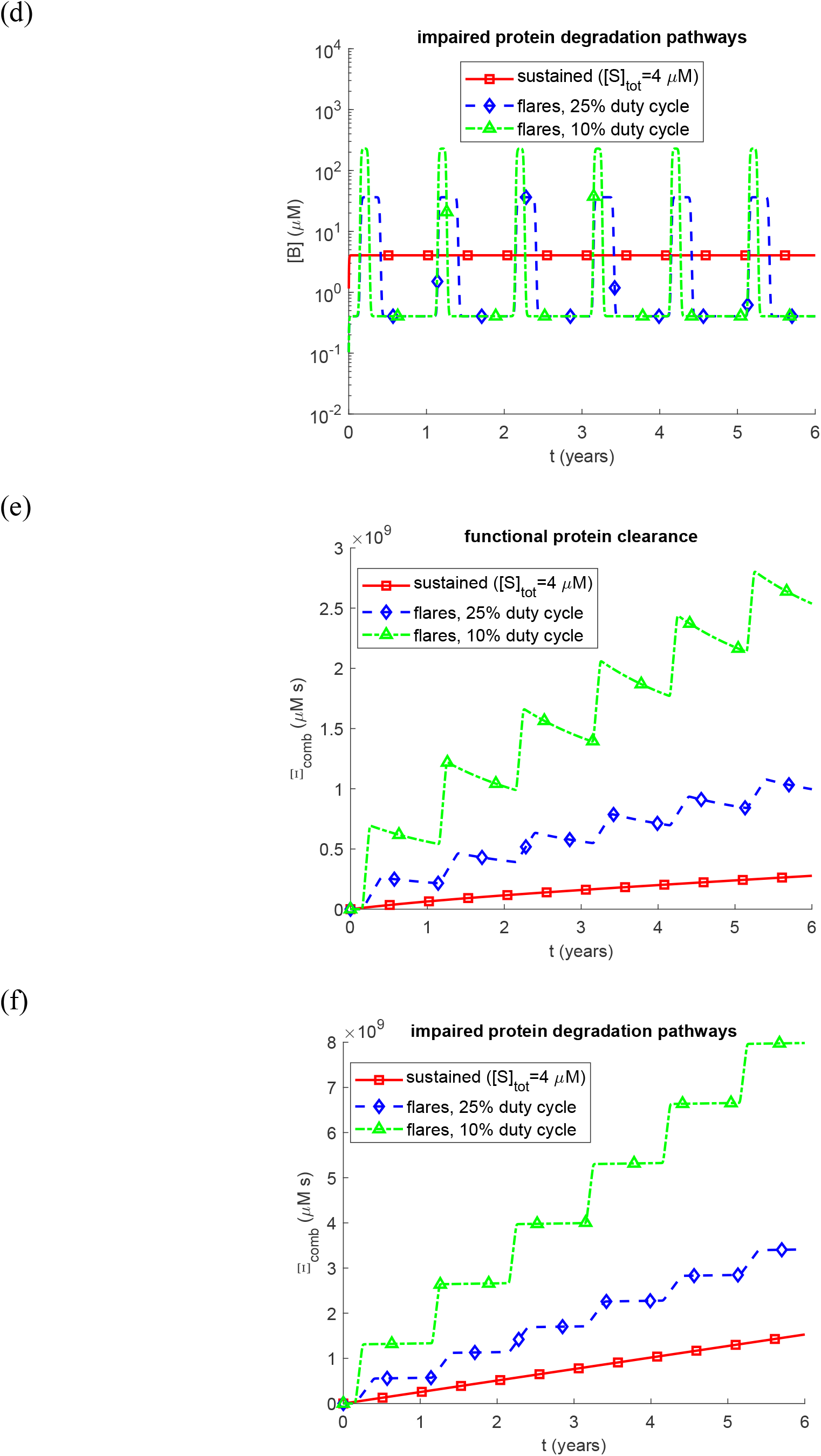
Effect of the temporal pattern of inflammation, traced from total SAA to combined renal damage, for three inflammatory histories having the same time-averaged total SAA concentration of 4 µM: sustained inflammation, annual flares with a 25% duty cycle, and annual flares with a 10% duty cycle. (a) Prescribed total SAA concentration; between flares, [*S*]_*tot*_ = 0.5 µM. (b) Free (HDL-unbound) SAA concentration in plasma, Eq. (9). (c, d) Concentration of free AA oligomers in the kidney, [*B*]. (e, f) Combined renal damage from accumulated AA oligomer cytotoxicity and insoluble fibril burden, Eq. (47). (c, e) Biologically relevant half-lives, representing functional protein degradation machinery. (d, f) Infinite half-lives, representing complete impairment of protein degradation. Panels (b)–(d) use logarithmic axes. Baseline parameters as in Fig. 2, with [*H*]_*tot*_ = 15 µM.

Having shown that the pattern of inflammation matters, Fig. S27 asks a different question: does the timing of the same inflammatory exposure matter? A single one-year episode, during which total SAA rises from 0.5 to 16 µM, is placed either early (0.5–1.5 years) or late (4.5–5.5 years) in a six-year course (Fig. S27a). With impaired protein degradation, the two episodes produce identical combined damage at six years (Fig. S27c), because in the absence of clearance both contributions depend only on cumulative oligomer exposure and are invariant to a shift of the same episode in time. With functional protein clearance, however, the late episode produces 45% more combined damage than the early episode (Fig. S27b). The accumulated toxicity of Eq. (45) is identical in the two cases, 7.4 × 10^8^ µM s; the entire difference comes from the fibrillar component, which is fourteen times larger after the late episode because fibrils deposited early have had more time to clear. The model therefore contains two distinct forms of memory: the oligomer-mediated component records cumulative exposure, whereas the fibrillar component retains information about timing through clearance. Consistent with this distinction, the instantaneous combined burden of Eq. (47) can decrease after an early episode as fibrils are removed.

Because the biological-age index of Eq. (49) follows this burden, it falls by up to 2.1 years after the early episode as the deposits formed during that episode are cleared, and at six years it is 3.7 years lower than after the late episode (64.1 versus 67.8 years); after the early episode it ends within 0.3 years of the floor of Eq. (50).

This separation between cumulative oligomer exposure and reversible fibril burden leads directly to the treatment-timing question in Fig. 10. Circulating SAA is reduced below 10 mg/L (0.83 µM), the level associated with favorable outcomes (Gillmore et al., 2001), beginning one, three, or five years after disease onset; without treatment, total SAA remains at 8 µM. Under impaired protein degradation, the ten-year renal biological-age index is 134 years without treatment and 71, 85, and 99 years when SAA suppression begins after one, three, and five years, respectively (Fig. 10b). With functional protein clearance, the corresponding values are 77 years without treatment and 62, 65, and 68 years with treatment (Fig. 10a). Earlier suppression therefore leaves a lower subsequent index in both scenarios. That ordering follows from the cumulative construction of the damage measure and is not, by itself, a unique dynamical prediction; the model instead quantifies its magnitude. Under impaired degradation, delaying treatment from one to five years increases the index by about 28 years, or approximately seven years of biological age per year of delay. This increment equals the difference between the damage-accrual rates at the untreated and target SAA concentrations and is therefore determined by HDL buffering and the aggregation regime. With functional clearance, the index rises much more slowly than chronological age once SAA is suppressed, and almost stops rising for a time when suppression begins after three or five years, because fibrils deposited before treatment are being cleared; under impaired degradation fibrils do not clear, and the index keeps rising at least as fast as chronological age. Values above BA = 100 years are extrapolations beyond the calibrated terminal-damage threshold and should not be interpreted as literal physiological ages.

The final question is whether the modeled dependence on SAA is consistent with clinical observations. Fig. 11 provides this external consistency check, but not a fully independent validation: the SAA dependence shown in the figure was not used to tune model parameters, whereas the overall biological-age scale *c* was back-calibrated from the broader untreated-survival literature. Fig. 11a shows the predicted time from disease onset to the calibrated terminal-damage threshold of BA = 100 years as a function of median circulating SAA. At the clinical 10 mg/L threshold (Gillmore et al., 2001), the predicted time to the calibrated threshold is 48 years with functional protein clearance and 34 years with impaired degradation. At 50 mg/L these times fall to 34 and 13 years, respectively, and at 100 mg/L to 17 and 5 years. The model is therefore consistent with the qualitative clinical separation (Gillmore et al., 2001): sustained low SAA is associated with stabilization or regression of amyloid burden, whereas persistently high SAA is associated with continued deposition and worsening renal function. Fig. 11b tests the strength of the dependence in another way. Between 10 and 100 mg/L, the model predicts a 1.28-fold increase in the growth rate of the biological-age index per doubling of SAA under functional clearance and a 1.70-fold increase under impaired degradation, compared with the 1.24-fold increase in the risk of progression to end-stage renal failure per doubling of SAA (Lachmann et al., 2007). These observables are not equivalent—the model describes a rate of damage accumulation, whereas the clinical study reports a hazard—so the comparison should not be interpreted as direct quantitative validation. It does, however, show agreement in direction and approximate magnitude. The comparison also depends on the calibration of *c*: over the 4–8-year untreated-survival range used to define the scale, the predicted increase per doubling of SAA spans 1.22–1.40 under functional clearance and 1.59–1.83 under impaired degradation. The clinical value of 1.24 therefore lies within the range predicted with functional clearance, although not every individual calibration reproduces it. Across the same calibration range, the predicted time to BA = 100 years at 10 mg/L remains 46–48 years with functional clearance, whereas at 50 mg/L it is 9–16 years with impaired degradation. Thus, the qualitative separation between low- and high-SAA outcomes is robust to the tested calibration uncertainty.

**Fig. 10.**
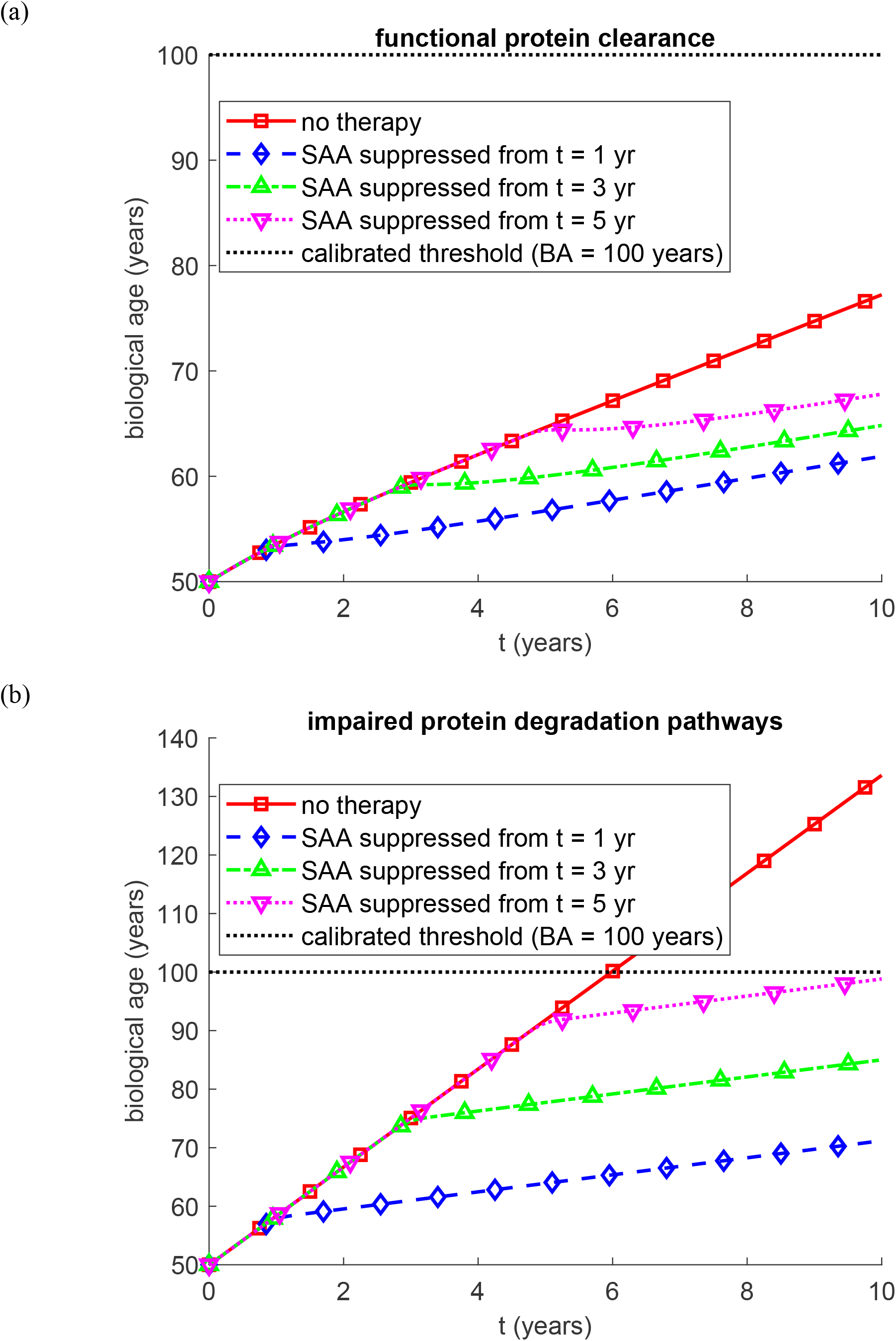
Renal biological age as a function of time when circulating SAA is suppressed below 10 mg/L (0.83 µM) starting at one, three, or five years after disease onset, compared with the untreated course at a total SAA concentration of 8 µM. (a) Biologically relevant half-lives, representing functional protein degradation machinery. (b) Infinite half-lives, representing complete impairment of protein degradation. The dotted horizontal line marks the calibrated terminal-damage threshold of BA = 100 years; values above this line are extrapolations of the index rather than literal physiological ages. Baseline parameters as in Fig. 2, with [*H*]_*tot*_ = 15 µM.

**Fig. 11.**
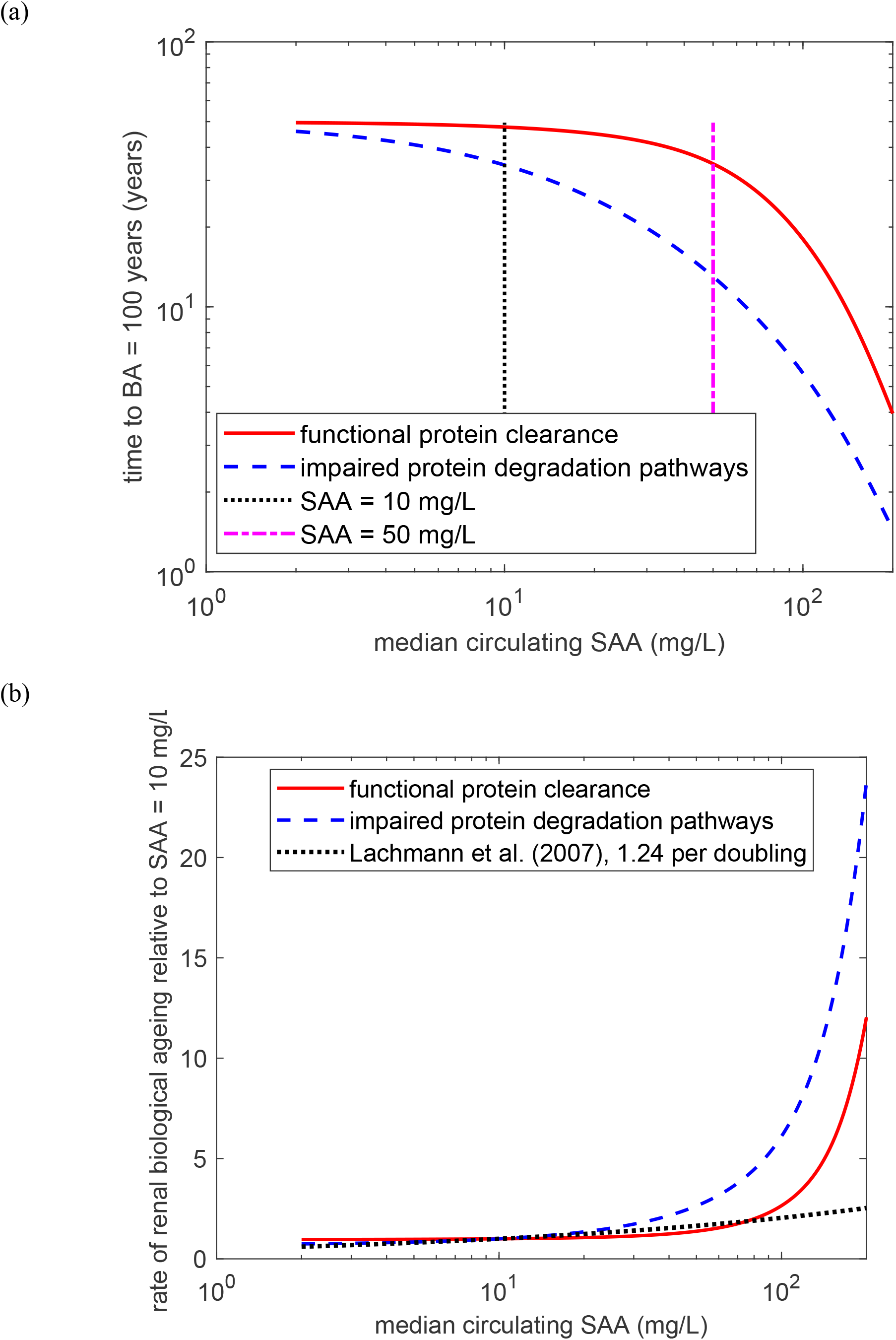
External consistency check against published clinical outcome data. The SAA dependence was not used for parameter tuning, although *c* was calibrated from the same general untreated-survival literature. (a) Predicted time from disease onset to the calibrated terminal-damage threshold of BA = 100 years as a function of the median circulating SAA concentration; vertical lines mark the clinical 10 mg/L and 50 mg/L thresholds (Gillmore et al., 2001). (b) Predicted rate of renal biological aging relative to its value at 10 mg/L, compared with the 1.24-fold increase in the risk of progression to end-stage renal failure per doubling of SAA (Lachmann et al., 2007). Baseline parameters as in Fig. 2, with [*H*]_*tot*_ = 15 µM.

Because the absolute biological-age scale is calibrated, Fig. S28 asks which conclusions remain robust when that calibration is varied. Over the reported 4–8-year range of untreated survival (Gillmore et al., 2001), (Ahbap et al., 2014), the implied value of *c* changes from 0.522 to 0.238 µM^−1^, a factor of 2.2 (Fig. S28a). The absolute magnitudes of the dimensionless sensitivity coefficients therefore change with *c* (Fig. S28b). Their signs and their ranking from most to least influential, however, remain unchanged across the entire range. The sensitivity analysis can therefore support conclusions about which parameters are most influential, whereas statements expressed as absolute numbers of years of renal biological age remain calibration-dependent.

## 4. Discussion, limitations of the model, and future directions

The model suggests that AA amyloidosis progression is governed by the dynamic equilibrium between circulating SAA and the protective buffering capacity of HDL. Within the bloodstream, HDL functions as a chaperone-like carrier for SAA, binding its N-terminal lipid-binding surface to sequester amyloidogenic segments and curtail SAA’s tendency to misfold and promote inflammation (Zhu et al., 2016).

While sufficient HDL binding capacity maintains circulating SAA in a sequestered, non-pathogenic state, chronic inflammation disrupts this protective equilibrium by simultaneously promoting SAA overproduction and reducing HDL levels (Jahangiri et al., 2009). In the model, this dual perturbation diminishes available HDL binding sites, allowing circulating SAA to overwhelm the buffering capacity of HDL and generate a pathogenic pool of free, lipid-poor SAA (Webb, 2021), (Schuchardt et al., 2019), which may contribute to renal biological aging through fibril deposition and oligomer-mediated inflammation (Simons et al., 2013).

The model further predicts that higher total SAA concentrations increase this free SAA pool, accelerating renal filtration and biological aging. This finding is consistent with the central therapeutic strategy in AA amyloidosis: aggressive suppression of the underlying inflammatory disease using anti-inflammatory and immunosuppressive agents to reduce cytokine-driven SAA synthesis. Clinical evidence indicates that sustained SAA reduction or normalization, through biologic agents, disease-modifying drugs, or targeted antimicrobial therapy, stabilizes tissue AA deposits and preserves organ function (Real de Asúa et al., 2014).

Published clinical evidence provides an external consistency check on this model prediction. A prospective study (Gillmore et al., 2001) followed 80 patients and found that amyloid deposits regressed in 25 of the 42 patients whose median circulating SAA concentration remained within the reference range (<10 mg/L), and that organ function stabilized or improved in 39 of these 42 patients. In contrast, among patients with a median SAA concentration exceeding 10 mg/L, outcomes were less favorable: amyloid burden increased and organ function deteriorated in most patients with SAA persistently above 50 mg/L. Ten-year survival was approximately 90% in patients whose median SAA concentration remained below 10 mg/L, compared with 40% in those whose median concentration exceeded 10 mg/L. A larger prospective study of 374 patients (Lachmann et al., 2007) likewise demonstrated that mortality, amyloid burden, and renal prognosis were strongly associated with the SAA concentration maintained during follow-up. In particular, the risk of progression to end-stage renal failure increased by approximately 24% for each doubling of SAA concentration, while mortality was substantially higher among patients with persistently elevated SAA. These clinical findings are consistent with the model’s prediction that sustained control of circulating SAA, rather than isolated measurements of inflammatory activity, is a major determinant of long-term amyloid burden and renal outcome. They are also consistent with the model’s proposed mechanism whereby elevated total SAA increases the pool of free, HDL-unbound SAA available for renal filtration and subsequent amyloidogenic processing. However, because these clinical studies measured circulating total SAA rather than the free, HDL-unbound fraction, they provide indirect consistency evidence rather than direct validation of the model’s specific free-SAA mechanism.

Taken together, these observations raise a more useful question than whether the model reproduces individual patient outcomes: does it reproduce the observed dependence of renal progression on circulating SAA? Fig. 11 provides a semi-quantitative check. When median SAA is held at or below 10 mg/L, the predicted time to the calibrated terminal-damage threshold of BA = 100 years exceeds the remaining life expectancy of a patient diagnosed at age 50; under impaired protein degradation, this time falls to about 13 years at 50 mg/L and 5 years at 100 mg/L. This low-versus high-SAA separation is qualitatively consistent with the published outcomes (Gillmore et al., 2001). The same trend appears in the response slope: under functional clearance, the model predicts a 1.28-fold increase in the growth rate of the biological-age index for each doubling of SAA, close to the 1.24-fold increase in end-stage renal failure risk (Lachmann et al., 2007). This agreement should not be interpreted as independent validation. No parameter was tuned specifically to reproduce the SAA-response slope or the 10/50 mg/L clinical comparison, but the absolute biological-age scale is inherited from the calibration of *c*, which maps a six-year untreated reference course to BA = 100 years using untreated-survival data drawn partly from the same clinical literature (Gillmore et al., 2001). Thus, the absolute timescale in Fig. 11a is calibrated rather than predicted. What Fig. 11 does test is the model-predicted dependence on SAA: the clinically reported response slope lies within the range predicted with functional clearance across the admissible values of *c*. Even this comparison remains non-specific, however, because the clinical studies measured total rather than free SAA and because any other mechanism under which renal outcome worsens monotonically with SAA could produce a similar trend.

That distinction between consistency and validation points directly to the next test. A direct longitudinal evaluation would follow the same patients over time with serial measurements of total SAA, HDL, and, ideally, free SAA, together with renal function or proteinuria and amyloid burden. The National Amyloidosis Centre cohorts (Gillmore et al., 2001), (Lachmann et al., 2007) already combine repeated SAA measurements with serum amyloid P component scintigraphy and renal outcomes, and related inflammatory-disease registries contain longitudinal inflammatory markers and renal endpoints. Such data would allow the central prediction of Fig. 9 to be tested directly: among patients with similar time-averaged SAA, more intermittent trajectories should be associated with faster progression. Ideally, model calibration and validation would be performed on separate patient subsets. Obtaining and analyzing such data is beyond the scope of the present study, but such an analysis is the most direct route to falsifying— or strengthening—the proposed mechanism.

Before interpreting the time-varying simulations, it is important to separate what is built into the damage metric from what emerges from the model dynamics. First, the accumulated toxicity of Eq. (45), a measure of cumulative oligomer exposure, is non-decreasing by definition, because it is the time integral of a non-negative oligomer concentration. Second, renal biological age cannot fall below the floor of Eq. (50), because that floor contains only chronological time and the accumulated oligomer toxicity. Neither property is claimed as a result; the contribution of the present work lies in representing renal injury in AA amyloidosis through cumulative oligomer exposure and quantifying the consequences. The genuinely dynamical results lie elsewhere: intermittent flares produce nonlinear amplification at fixed mean SAA (Fig. 9), the oligomeric and fibrillar components respond differently to the timing of an inflammatory episode (Fig. S27), the extent of biological-age recovery toward that floor after SAA control depends on fibril clearance (Fig. 10a), the aggregation system changes regime near κ_2_ = 1 (Fig. 8), and the progression rate depends nonlinearly on circulating SAA (Fig. 11). Importantly, the instantaneous combined burden of Eq. (47) can decrease when fibrils are cleared, so the model allows partial regression of amyloid deposits after SAA control and a corresponding partial recovery of renal biological age, consistent with clinical observations (Gillmore et al., 2001). The model therefore assigns different kinds of memory to oligomer exposure and fibril burden: clearing fibrils can reduce the current structural burden, but it does not erase oligomer exposure that has already accumulated. A fibril-directed intervention, such as the fibril-specific monoclonal antibody approach (Thorne et al., 2022), would therefore be expected to lower the fibrillar contribution without resetting the cumulative oligomer-exposure term. This distinction is biologically plausible because prefibrillar aggregation species can be strongly cytotoxic, in some cases more so than mature fibrils (Bucciantini et al., 2002). The model’s claim is thus not that all renal damage is mathematically irreversible, but that oligomer exposure and fibril deposition carry different forms of pathological memory.

With that distinction in place, the meaning of path dependence becomes more precise. Fig. 9 shows that the temporal pattern of SAA matters: intermittent flares are substantially more damaging than sustained inflammation with the same mean SAA because nonlinear HDL saturation and autocatalytic aggregation amplify peaks. Fig. S27 then asks a different question—whether the same inflammatory episode is intrinsically more damaging when it occurs early or late. The answer depends on which component is considered. The accumulated oligomer exposure is unchanged by shifting an otherwise identical episode in time, whereas the fibrillar contribution is larger after a late episode when clearance is active because there has been less time for deposited fibrils to regress. The clinically testable prediction is therefore not that timing alone determines irreversible injury, but that the shape of the SAA trajectory—particularly its peak-to-mean structure—contains information beyond the time average, while episode timing affects the residual fibril burden through clearance. The treatment simulations are consistent with this picture: earlier SAA suppression leaves a lower subsequent biological-age index, but this result should be interpreted as a comparison among model scenarios rather than as a patient-specific estimate of irreversible years lost.

A distinction between local and global behavior also resolves the apparent tension between the sensitivity analysis and the uncertainty in the aggregation kinetics. Table 3 asks which parameters matter most near the baseline operating point. There, the system lies above the autocatalytic threshold and is supply-limited, so biological age is most sensitive to total SAA concentration and HDL binding-site capacity, with SAA–HDL affinity, lysosomal processing, and the oligomer half-deposition time next in rank; the two Finke–Watzky constants have small local coefficients. Fig. 8 asks a different question: what happens if the poorly constrained aggregation parameters are varied enough to move the system into another regime? At lower *k*_2_, the model becomes nucleation-limited and the aggregation kinetics become important. Thus, a small local sensitivity to *k*_2_ at baseline does not imply that *k*_2_ is generally unimportant. Table 3 describes sensitivity within one regime, whereas Fig. 8 describes uncertainty about which regime applies.

These conclusions also make clear where the present model is strongest and where it remains provisional. The SAA–HDL module captures finite binding capacity but treats effective sites as equivalent and independent; site heterogeneity, cooperativity, and HDL remodeling are not resolved, and the number of effective sites per particle is inferred rather than measured. The aggregation module is deliberately minimal: the Finke–Watzky scheme does not distinguish oligomer sizes or toxicities, and *k*_1_ and *k*_2_ have not been measured for AA protein. The kidneys are represented as a single spatially homogeneous compartment, so regional filtration, mesangial trapping, tubular uptake, and local deposition are not resolved. These structural simplifications are compounded by parameter uncertainty: the reference HDL particle concentration comes from a cardiovascular cohort, the oligomer half-deposition time is transferred from amyloid-β plaque growth, and several quantities are inferred rather than directly measured. The uncertainty in *k*_2_ is particularly important because, as Fig. 8 shows, this parameter can change the aggregation regime. The biological-age formulation adds two further assumptions: *c* is back-calibrated to a reference terminal-damage scenario, and the index recovers only through the clearance of fibrillar deposits, so repair of oligomer-mediated injury is not represented and the floor of Eq. (50) may overstate how permanent that injury is. Finally, the clinical comparison relies on published aggregate outcomes rather than longitudinal patient-level trajectories. For these reasons, the present framework should be used to identify mechanisms, compare scenarios, and design discriminating experiments—not to guide personalized treatment. The most important next steps are AA-specific measurements of aggregation kinetics and oligomer turnover, better characterization of SAA–HDL occupancy, and longitudinal patient data that jointly track SAA, HDL, renal function, and amyloid burden in separate calibration and validation cohorts. Those measurements would determine whether the hypothesis-generating framework can be developed into a quantitatively predictive model.

## Author Contribution Statement

AVK is the sole author of this paper.

## Data Availability Statement

No experimental data were generated in this study. The complete MATLAB source code that reproduces the figures in the main text and Supplemental Materials is provided as a supplementary archive and is deposited in a public Zenodo repository (Kuznetsov, 2026b). The package comprises 22 scripts that reproduce every figure and Table 3, together with eight supporting routines and a README that lists, for every figure, the script that reproduces it.

## Ethics Statement

None.

## Abbreviations

AA: amyloid A
Aβ: amyloid-β
BA: biological age
F–W: Finke–Watzky
GFR: glomerular filtration rate
HDL: high-density lipoprotein
SAA: serum amyloid A

## Supplemental Materials

### S1. Numerical solution

The system of ordinary differential equations (15), (19), (20), and (26) with the prescribed initial conditions (32) was solved using MATLAB’s ode45 solver (R2023a; MathWorks, Natick, MA, USA). This numerical integrator implements an adaptive step-size Runge–Kutta method, making it suitable for problems requiring high numerical accuracy with reasonable computational cost. The solver was configured with stringent tolerances, setting both relative and absolute error thresholds to 10^−10^. The simulations of time-varying inflammation (Figs. 9, 10, and S27) use the same solver and tolerances with a right-hand side in which the total SAA concentration is prescribed as a function of time and the free SAA concentration follows from Eq. (9); the onsets and offsets of flares and inflammatory episodes are smoothed over a two-day interval, consistent with the one-to two-day rise and fall times of the acute-phase SAA response, which also keeps the right-hand side differentiable for the adaptive integrator. The results in Figs. 8 and 11 are obtained from the closed-form steady state of Eqs. (16), (25), and (27) and are verified against the numerical solution.

### S2. Supplementary figures

**Fig. S1.**
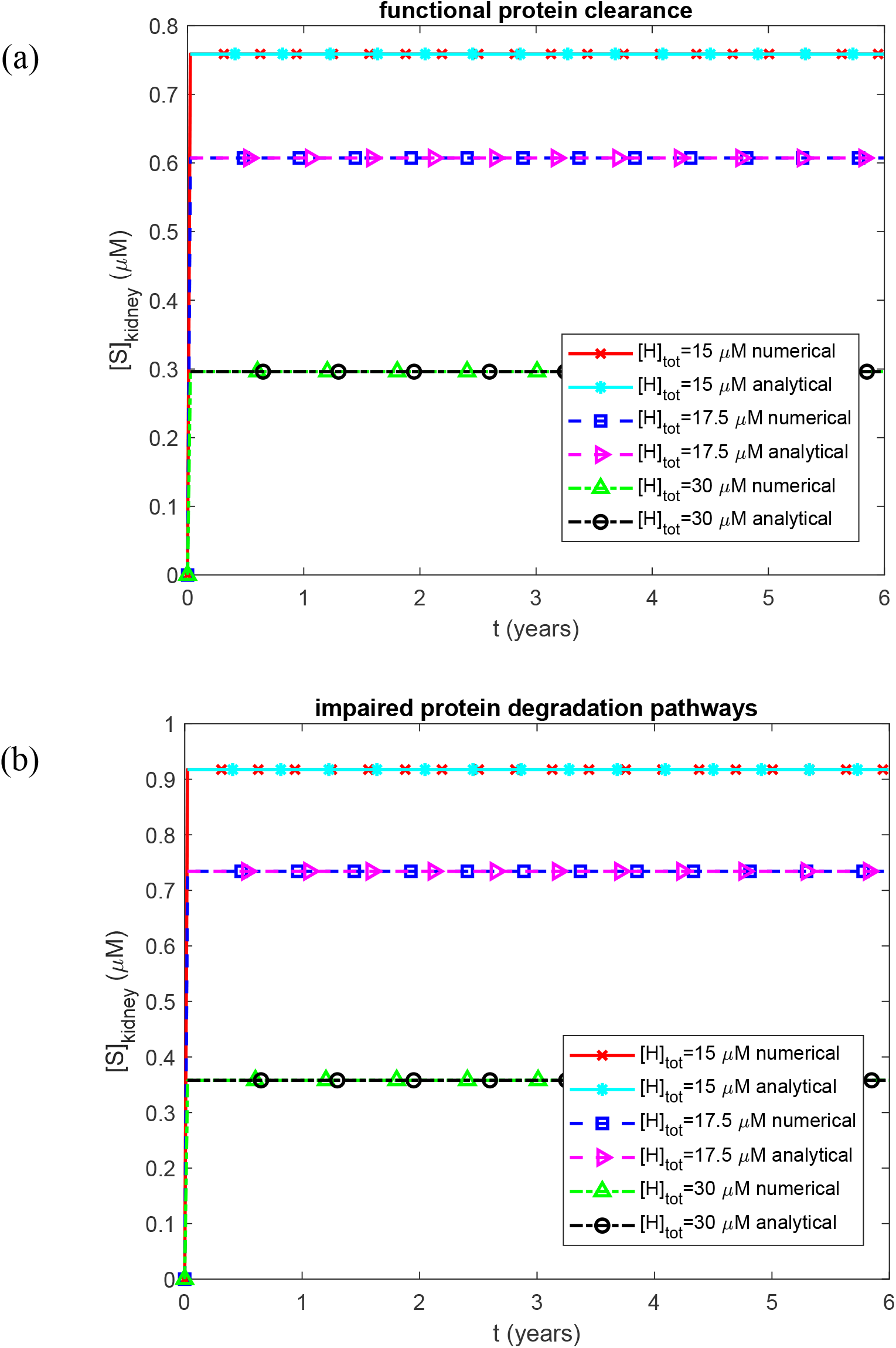
Molar concentration of SAA in the kidneys as a function of time. (a) Biologically relevant half-lives *T*_1/2, *A*_, *T*_1/2,*B*_, *T*_1/2,*I*_, and *T*_1/2,*S*_, representing functional protein degradation machinery. (b) Infinite half-lives *T*_1/2, *A*_, *T*_1/2,*B*_, *T*_1/2,*I*_, and *T*_1/2,*S*_, representing complete impairment of protein degradation. Each panel shows results for three values of the effective concentration of SAA-binding sites on HDL in plasma. Baseline parameters: *k*_1_ =10^−4^ s^−1^, *k*_2_ = 10^−4^ µM^−1^ s^−1^, *k*_*cl*_ = 9.65×10^−5^ s^−1^, *K*_*d*_ = 1.1789 *µ*M, *h*_*S*_ = 5.55×10^−7^ m s^−1^, *S*_*tot*_ = 8 *µ*M, and *θ*_1/2,*B*_ = 8.64 ×10^4^ s; remaining parameters as listed in Table 2.

**Fig. S2.**
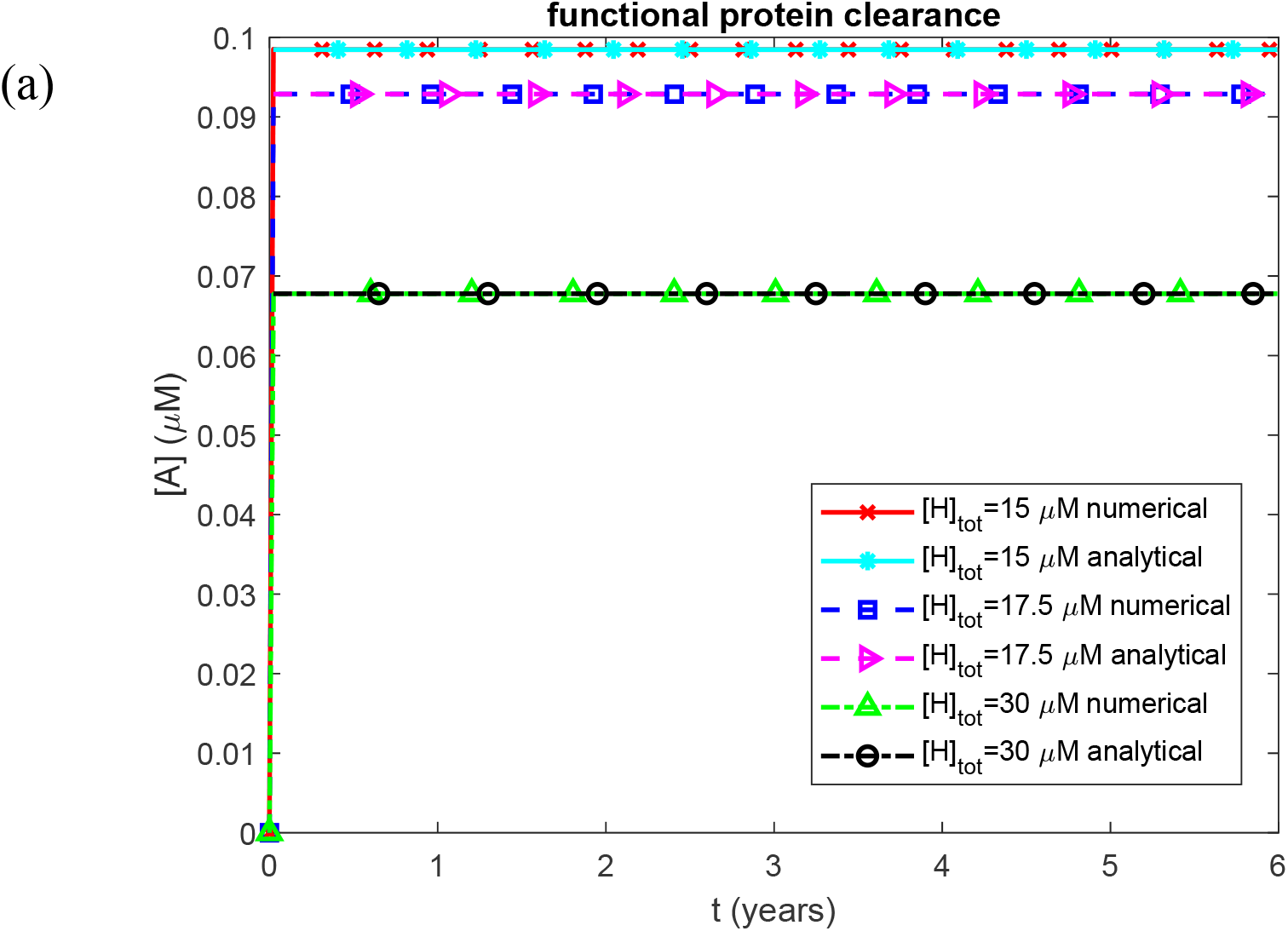
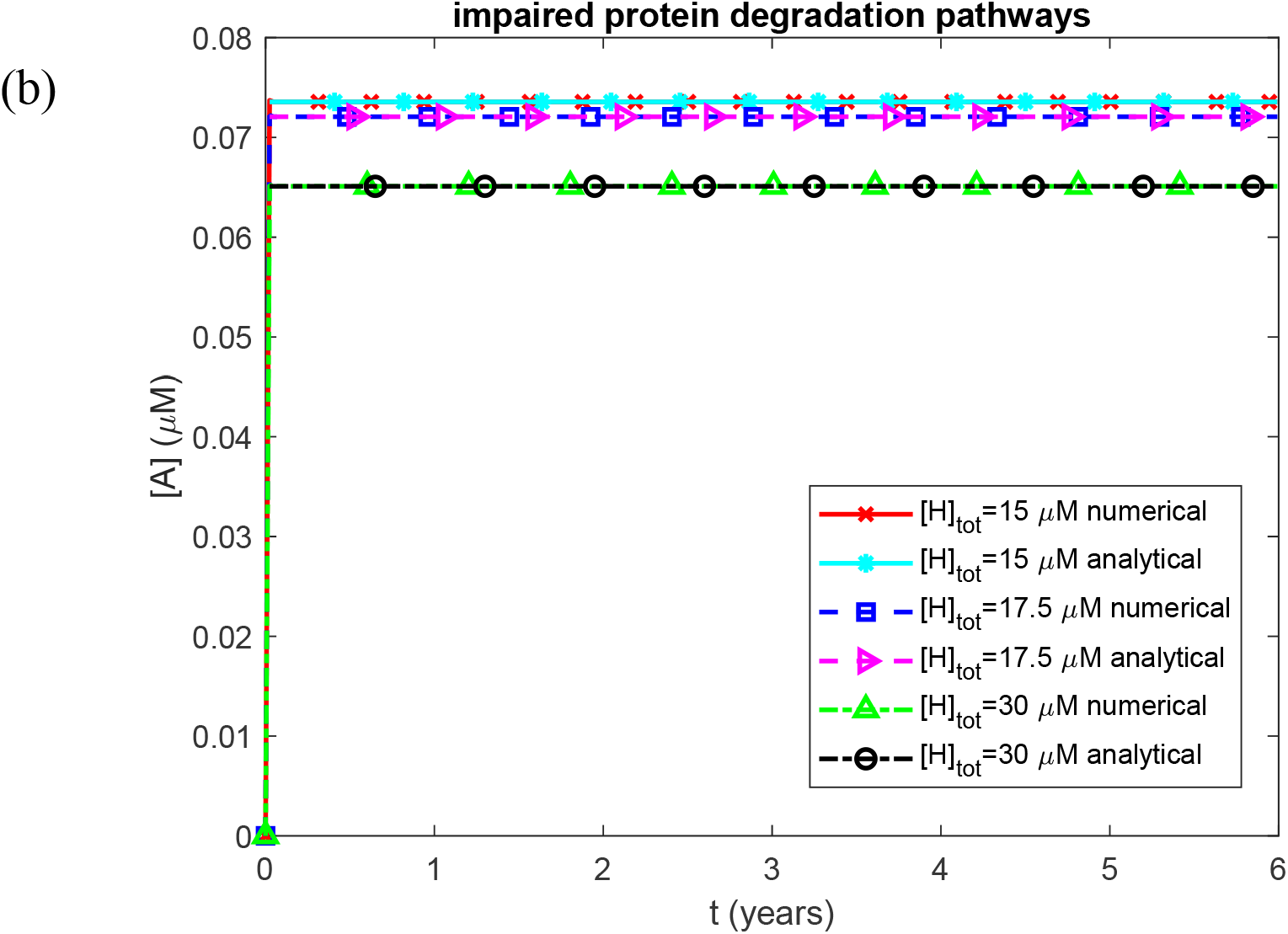
Molar concentration of AA fragments (monomers) in kidneys as a function of time. (a) Biologically relevant half-lives *T*_1/2, *A*_, *T*_1/2,*B*_, *T*_1/2,*I*_, and *T*_1/2,*S*_, representing functional protein degradation machinery. (b) Infinite half-lives *T*_1/2, *A*_, *T*_1/2,*B*_, *T*_1/2,*I*_, and *T*_1/2,*S*_, representing complete impairment of protein degradation. Each panel shows results for three values of the effective concentration of SAA-binding sites on HDL in plasma. Baseline parameters: *k*_1_ =10^−4^ s^−1^, *k*_2_ = 10^−4^ µM^−1^ s^−1^, *k*_*cl*_ = 9.65×10^−5^ s^−1^, *K*_*d*_ = 1.1789 *µ*M, *h*_*S*_ = 5.55×10^−7^ m s^−1^, *S*_*tot*_ = 8 *µ*M, and *θ*_1/2,*B*_ = 8.64 ×10^4^ s; remaining parameters as listed in Table 2.

**Fig. S3.**
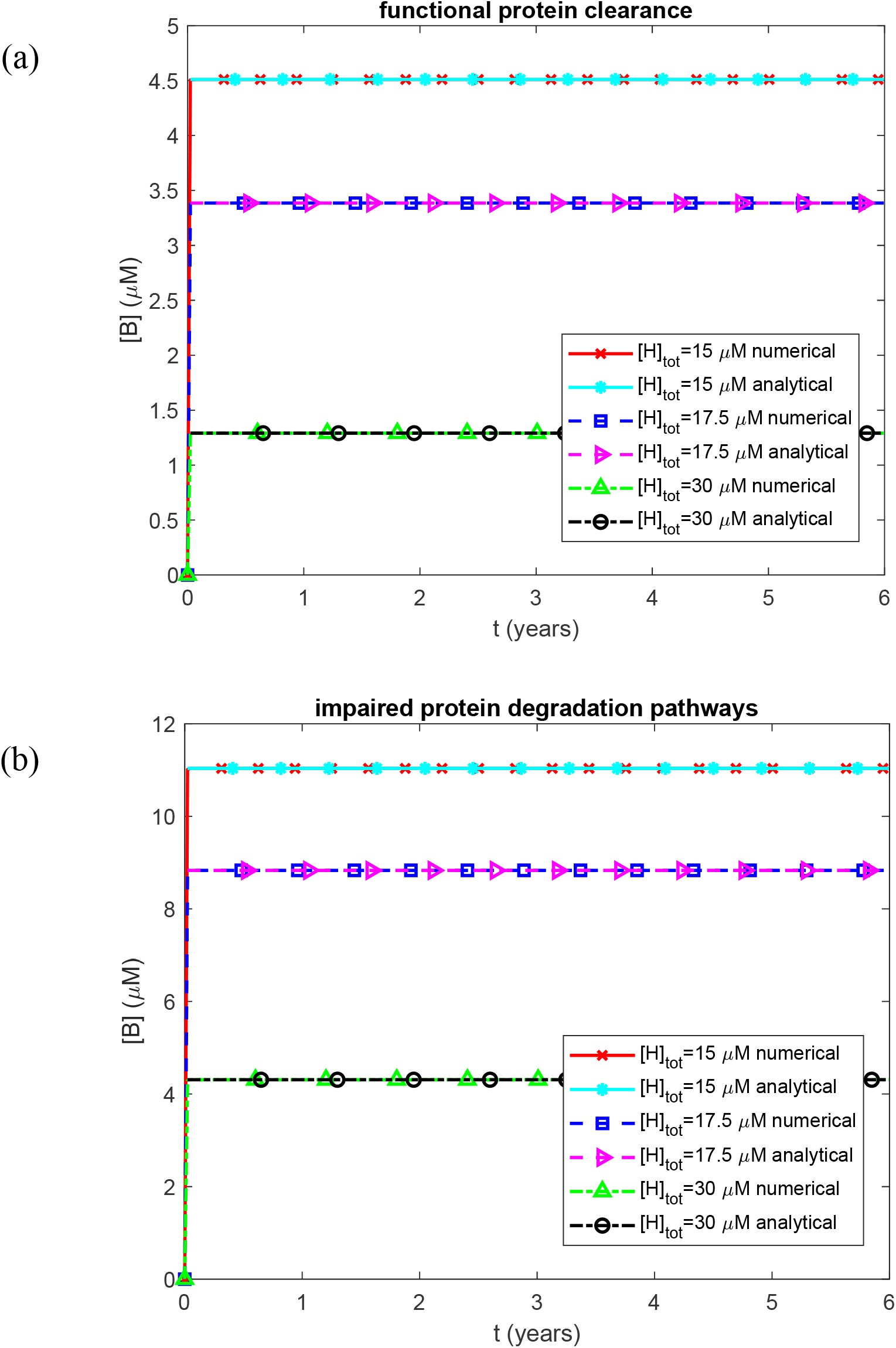
Molar concentration of free (non-deposited) AA oligomers in kidneys as a function of time. (a) Biologically relevant half-lives *T*_1/2, *A*_, *T*_1/2,*B*_, *T*_1/2,*I*_, and *T*_1/2,*S*_, representing functional protein degradation machinery. (b) Infinite half-lives *T*_1/2, *A*_, *T*_1/2,*B*_, *T*_1/2,*I*_, and *T*_1/2,*S*_, representing complete impairment of protein degradation. Each panel shows results for three values of the effective concentration of SAA-binding sites on HDL in plasma. Baseline parameters: *k*_1_ =10^−4^ s^−1^, *k*_2_ = 10^−4^ µM^−1^ s^−1^, *k*_*cl*_ = 9.65×10^−5^ s^−1^, *K*_*d*_ = 1.1789 *µ*M, *h*_*S*_ = 5.55×10^−7^ m s^−1^, *S*_*tot*_ = 8 *µ*M, and *θ*_1/2,*B*_ = 8.64 ×10^4^ s; remaining parameters as listed in Table 2.

**Fig. S4.**
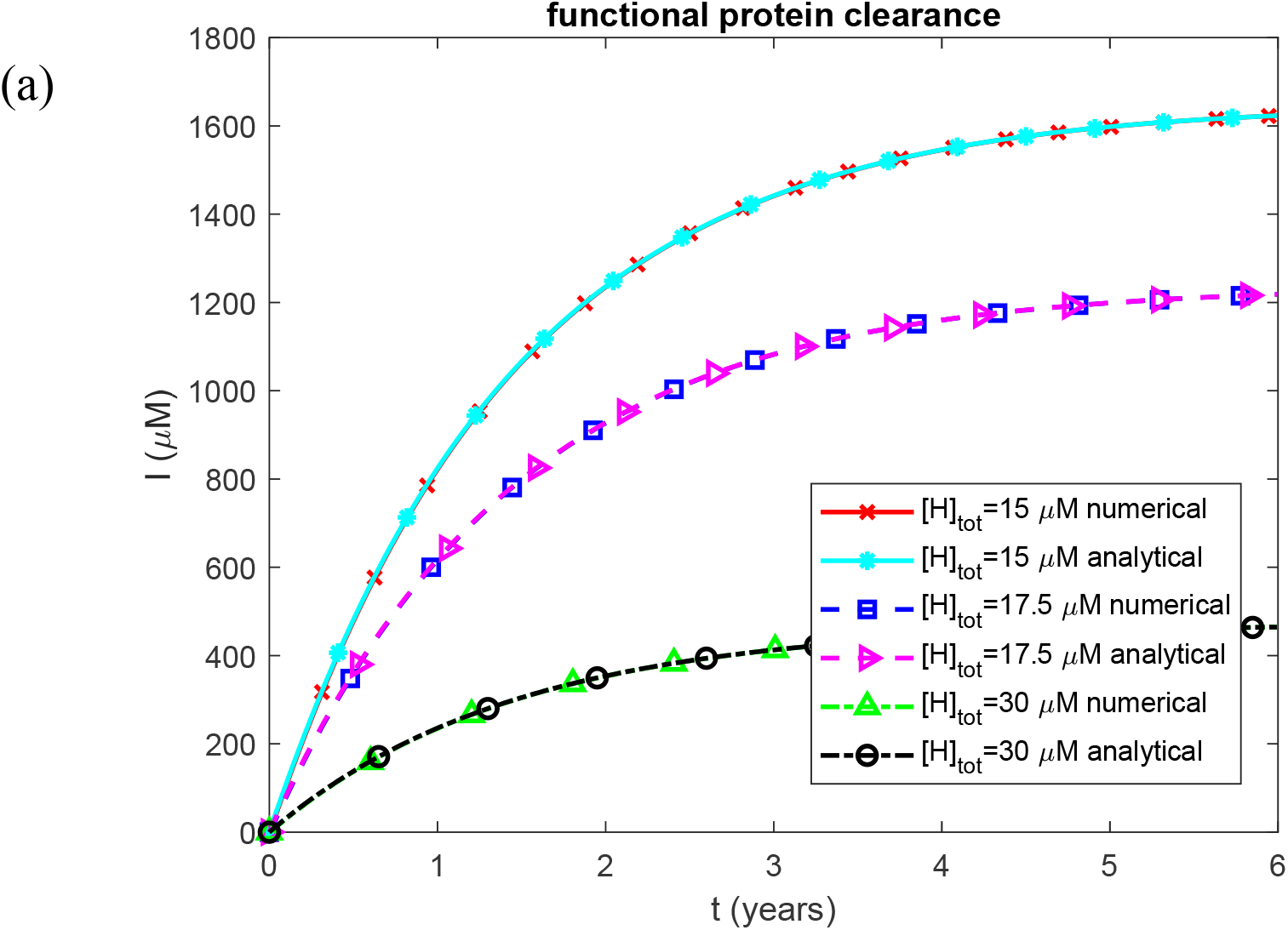
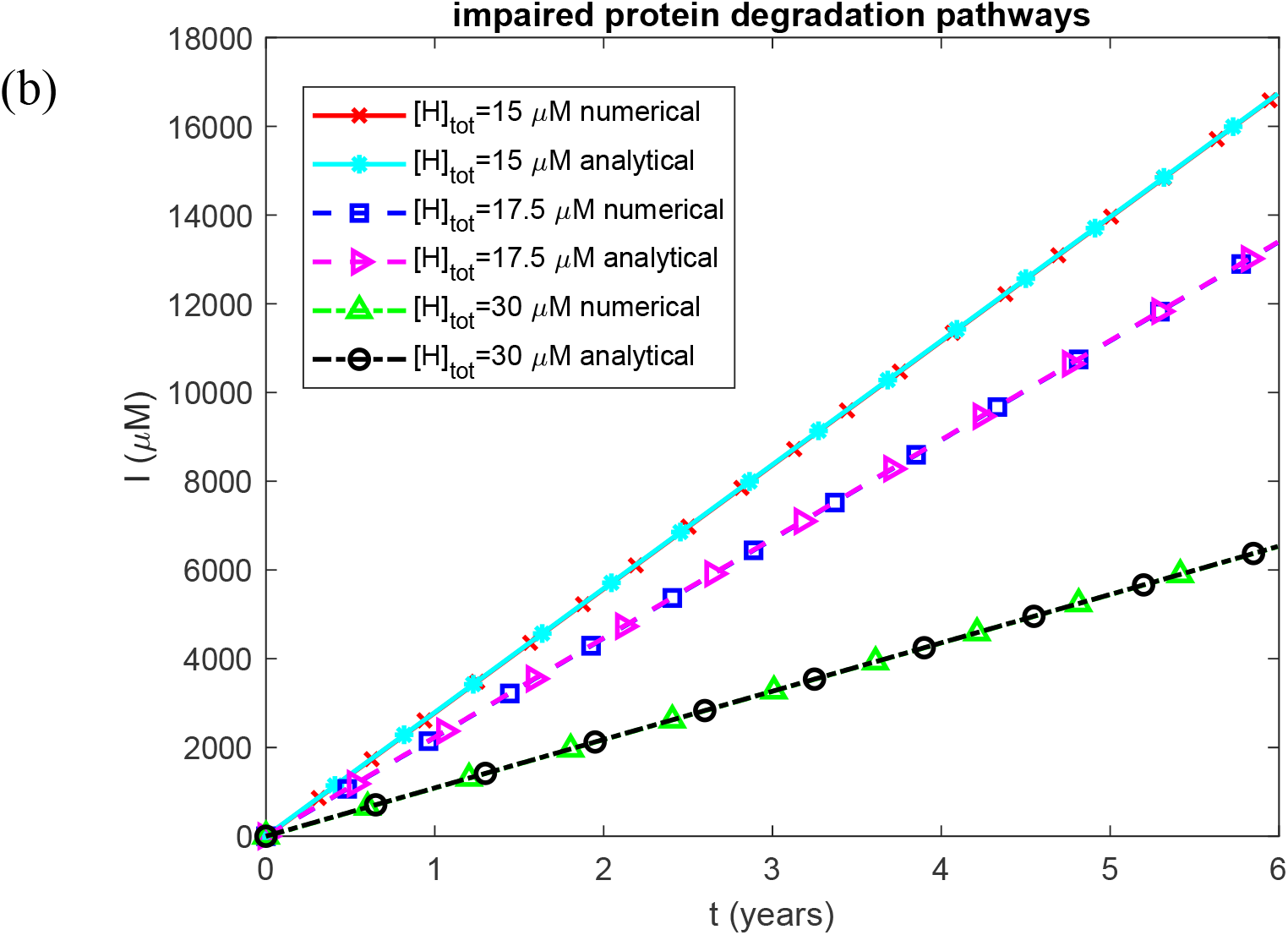
Molar concentration of AA oligomers deposited into AA amyloid fibrils in kidneys as a function of time. (a) Biologically relevant half-lives *T*_1/2, *A*_, *T*_1/2,*B*_, *T*_1/2,*I*_, and *T*_1/2,*S*_, representing functional protein degradation machinery. (b) Infinite half-lives *T*_1/2, *A*_, *T*_1/2,*B*_, *T*_1/2,*I*_, and *T*_1/2,*S*_, representing complete impairment of protein degradation. Each panel shows results for three values of the effective concentration of SAA-binding sites on HDL in plasma. Baseline parameters: *k*_1_ =10^−4^ s^−1^, *k*_2_ = 10^−4^ µM^−1^ s^−1^, *k*_*cl*_ = 9.65×10^−5^ s^−1^, *K*_*d*_ = 1.1789 *µ*M, *h*_*S*_ = 5.55×10^−7^ m s^−1^, *S*_*tot*_ = 8 *µ*M, and *θ*_1/2,*B*_ = 8.64 ×10^4^ s; remaining parameters as listed in Table 2.

**Fig. S5.**
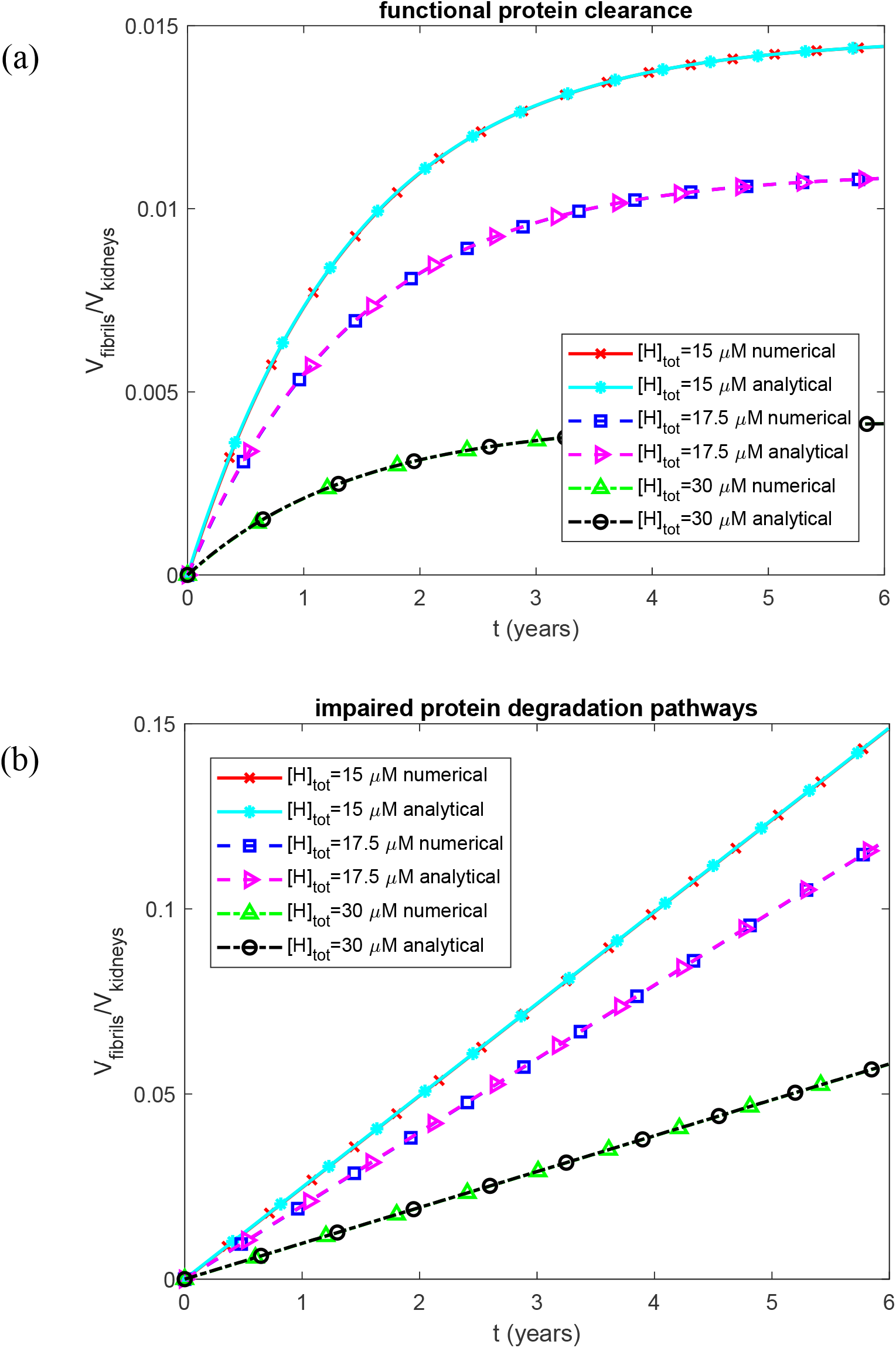
Fraction of the baseline kidney volume occupied by fibrils as a function of time. (a) Biologically relevant half-lives *T*_1/2, *A*_, *T*_1/2,*B*_, *T*_1/2,*I*_, and *T*_1/2,*S*_, representing functional protein degradation machinery. (b) Infinite half-lives *T*_1/2, *A*_, *T*_1/2,*B*_, *T*_1/2,*I*_, and *T*_1/2,*S*_, representing complete impairment of protein degradation. Each panel shows results for three values of the effective concentration of SAA-binding sites on HDL in plasma. Baseline parameters: *k*_1_ =10^−4^ s^−1^, *k*_2_ = 10^−4^ µM^−1^ s^−1^, *k*_*cl*_ = 9.65×10^−5^ s^−1^, *K*_*d*_ = 1.1789 *µ*M, *h*_*S*_ = 5.55×10^−7^ m s^−1^, *S*_*tot*_ = 8 *µ*M, and *θ*_1/2,*B*_ = 8.64 ×10^4^ s; remaining parameters as listed in Table 2.

**Fig. S6.**
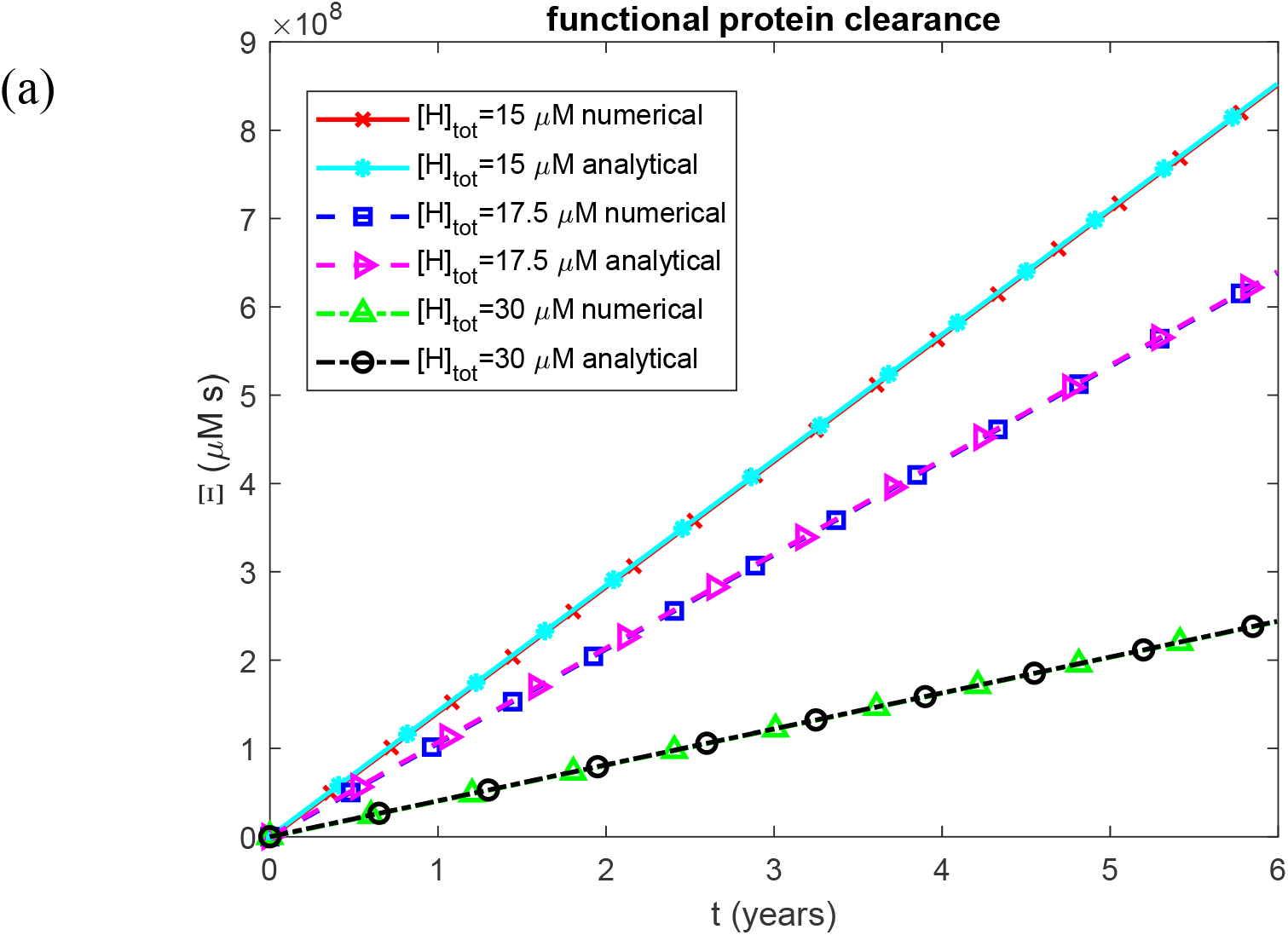
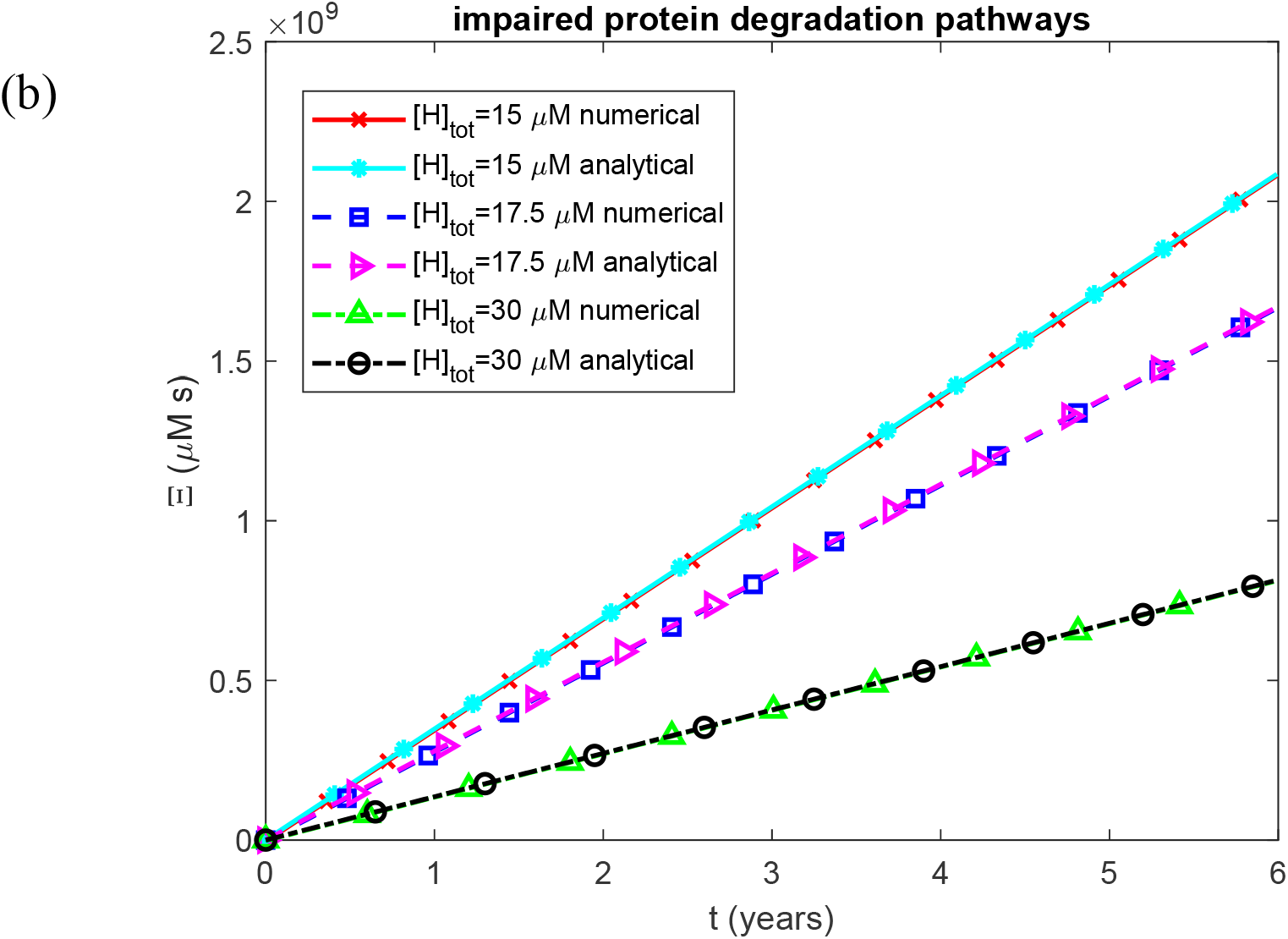
Accumulated nephrotoxicity of AA oligomers as a function of time. (a) Biologically relevant half-lives *T*_1/2, *A*_, *T*_1/2,*B*_, *T*_1/2,*I*_, and *T*_1/2,*S*_, representing functional protein degradation machinery. (b) Infinite half-lives *T*_1/2, *A*_, *T*_1/2,*B*_, *T*_1/2,*I*_, and *T*_1/2,*S*_, representing complete impairment of protein degradation. Each panel shows results for three values of the effective concentration of SAA-binding sites on HDL in plasma. Baseline parameters: *k*_1_ =10^−4^ s^−1^, *k*_2_ = 10^−4^ µM^−1^ s^−1^, *k*_*cl*_ = 9.65×10^−5^ s^−1^, *K*_*d*_ = 1.1789 *µ*M, *h*_*S*_ = 5.55×10^−7^ m s^−1^, *S*_*tot*_ = 8 *µ*M, and *θ*_1/2,*B*_ = 8.64 ×10^4^ s; remaining parameters as listed in Table 2.

**Fig. S7.**
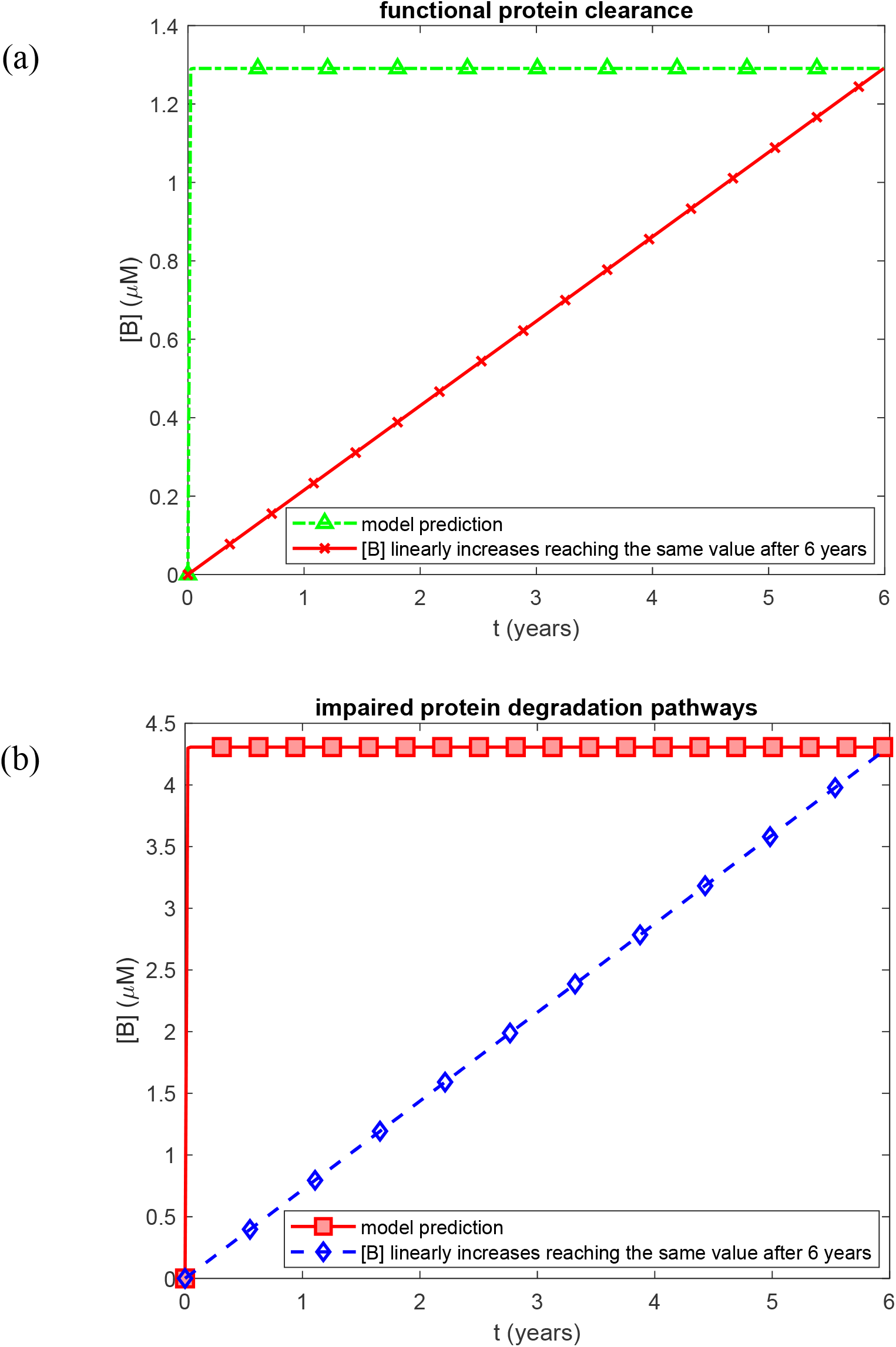
Molar concentration of free (non-deposited) AA oligomers in the kidneys as a function of time. Two cases are compared: the model prediction, in which [*B*] attains a constant steady-state value early in the disease course, and a hypothetical scenario in which [*B*] increases linearly throughout disease progression. (a) Biologically relevant half-lives *T*_1/2, *A*_, *T*_1/2,*B*_, *T*_1/2,*I*_, and *T*_1/2,*S*_, representing functional protein degradation machinery. (b) Infinite half-lives *T*_1/2, *A*_, *T*_1/2,*B*_, *T*_1/2,*I*_, and *T*_1/2,*S*_, representing complete impairment of protein degradation. Baseline parameters: *k*_1_ =10^−4^ s^−1^, *k*_2_ = 10^−4^ µM^−1^ s^−1^, *k*_*cl*_ = 9.65×10^−5^ s^−1^, *K*_*d*_ = 1.1789 *µ*M, *h*_*S*_ = 5.55×10^−7^ m s^−1^, *H*_*tot*_ = 30 *µ*M, *S*_*tot*_ = 8 *µ*M, and *θ*_1/2,*B*_ = 8.64 ×10^4^ s; remaining parameters as listed in Table 2.

**Fig. S8.**
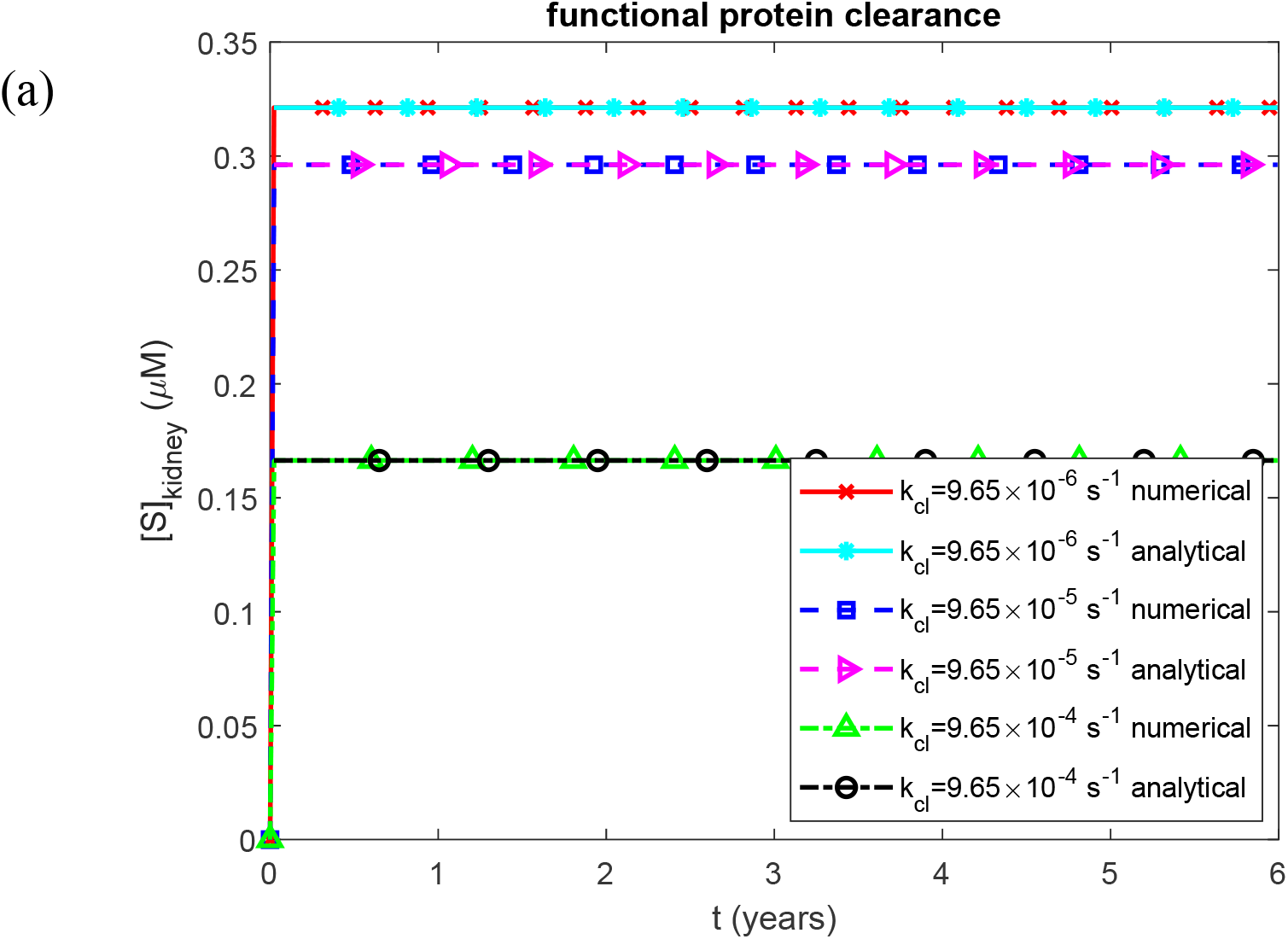
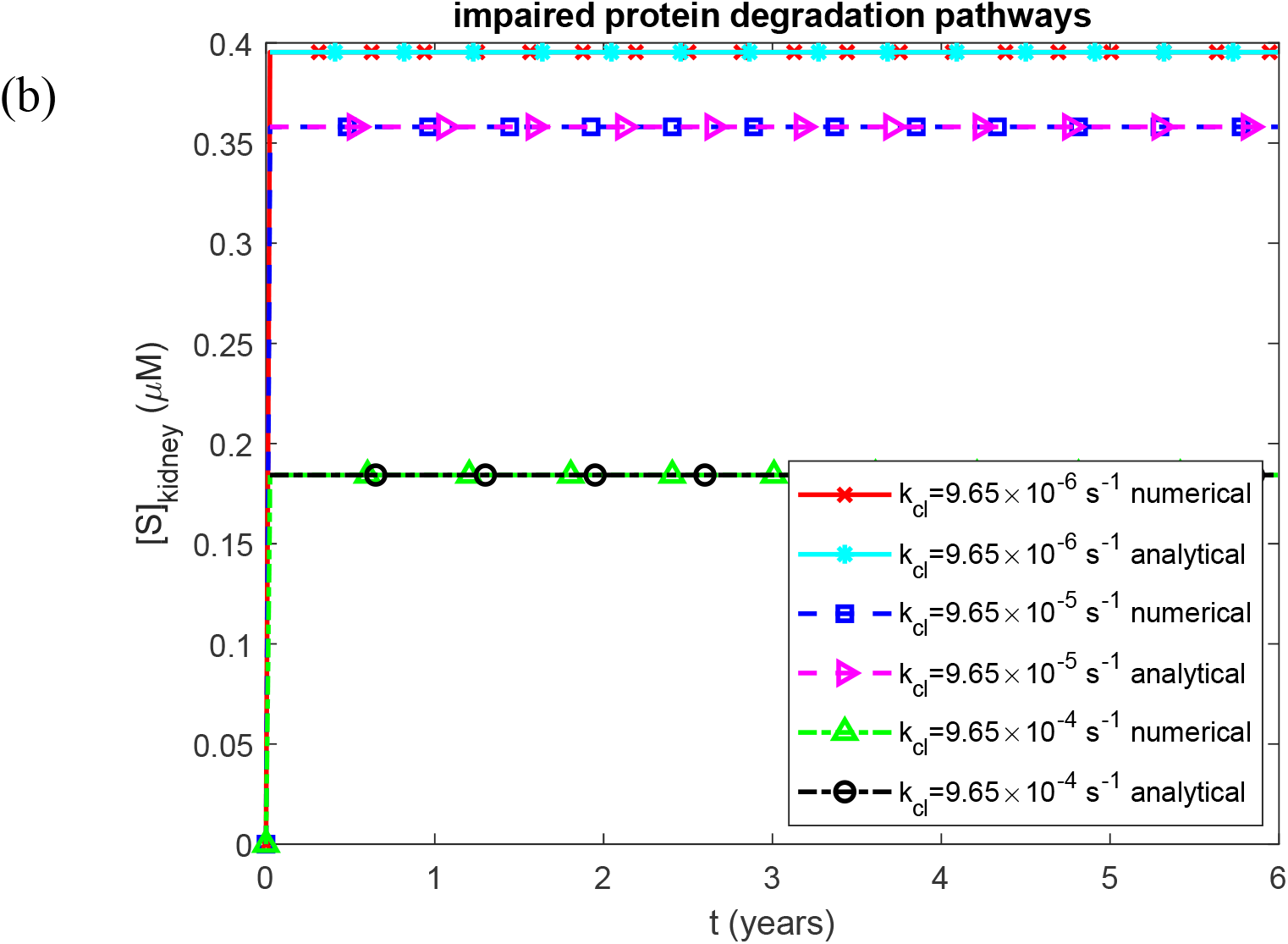
Molar concentration of SAA in the kidneys as a function of time. (a) Biologically relevant half-lives *T*_1/2, *A*_, *T*_1/2,*B*_, *T*_1/2,*I*_, and *T*_1/2,*S*_, representing functional protein degradation machinery. (b) Infinite half-lives *T*_1/2, *A*_, *T*_1/2,*B*_, *T*_1/2,*I*_, and *T*_1/2,*S*_, representing complete impairment of protein degradation. Each panel shows results for three values of the rate constant for proteolytic cleavage of SAA that generates AA monomers. Baseline parameters: *k*_1_ =10^−4^ s^−1^, *k*_2_ = 10^−4^ µM^−1^ s^−1^, *K*_*d*_ = 1.1789 *µ*M, *h*_*S*_ = 5.55×10^−7^ m s^−1^, *H*_*tot*_ = 30 *µ*M, *S*_*tot*_ = 8 *µ*M, and *θ*_1/2,*B*_ = 8.64 ×10^4^ s; remaining parameters as listed in Table 2.

**Fig. S9.**
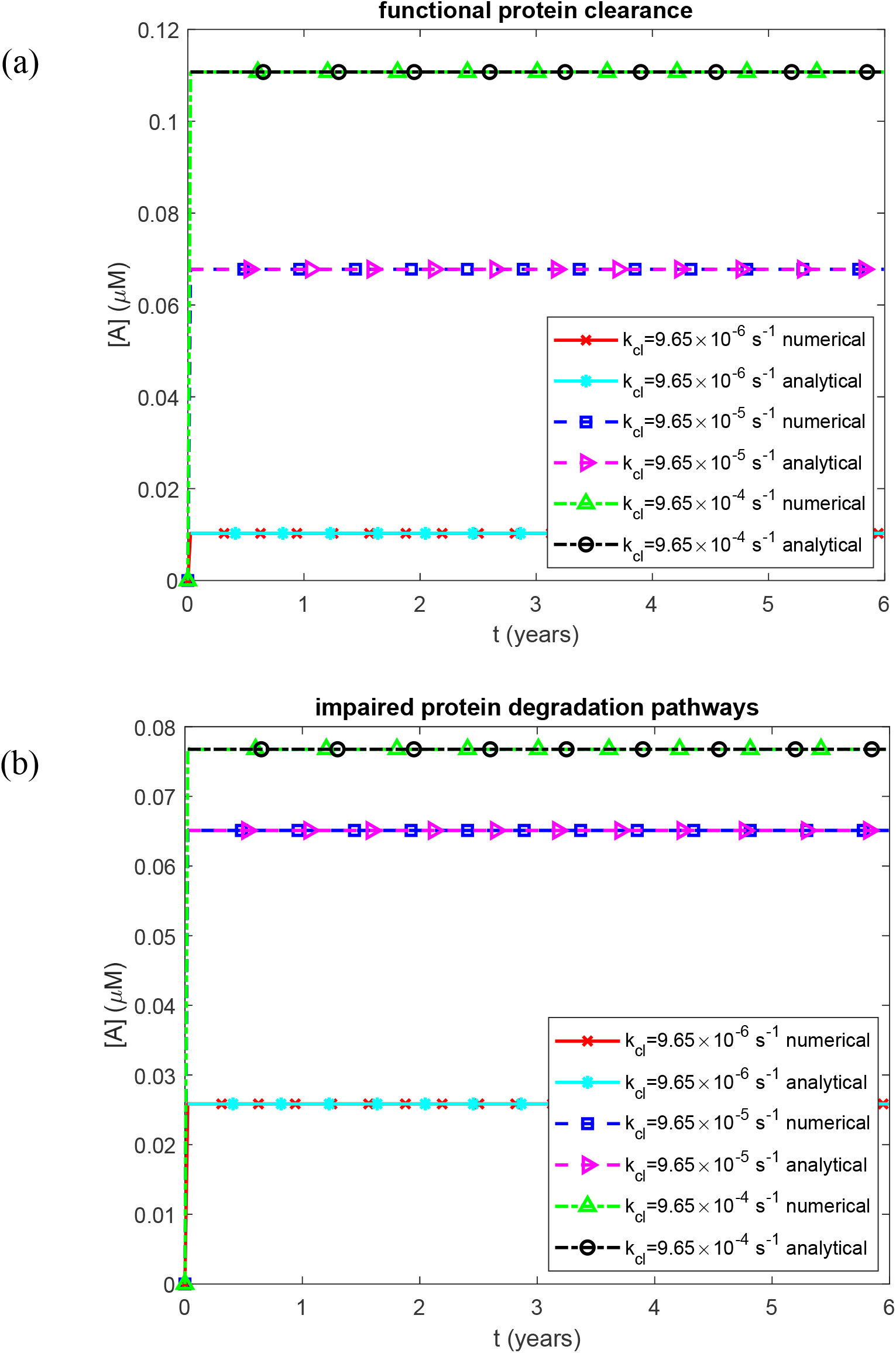
Molar concentration of AA fragments (monomers) in kidneys as a function of time. (a) Biologically relevant half-lives *T*_1/2, *A*_, *T*_1/2,*B*_, *T*_1/2,*I*_, and *T*_1/2,*S*_, representing functional protein degradation machinery. (b) Infinite half-lives *T*_1/2, *A*_, *T*_1/2,*B*_, *T*_1/2,*I*_, and *T*_1/2,*S*_, representing complete impairment of protein degradation. Each panel shows results for three values of the rate constant for proteolytic cleavage of SAA that generates AA monomers. Baseline parameters: *k*_1_ =10^−4^ s^−1^, *k*_2_ = 10^−4^ µM^−1^ s^−1^, *K*_*d*_ = 1.1789 *µ*M, *h*_*S*_ = 5.55×10^−7^ m s^−1^, *H*_*tot*_ = 30 *µ*M, *S*_*tot*_ = 8 *µ*M, and *θ*_1/2,*B*_ = 8.64 ×10^4^ s; remaining parameters as listed in Table 2.

**Fig. S10.**
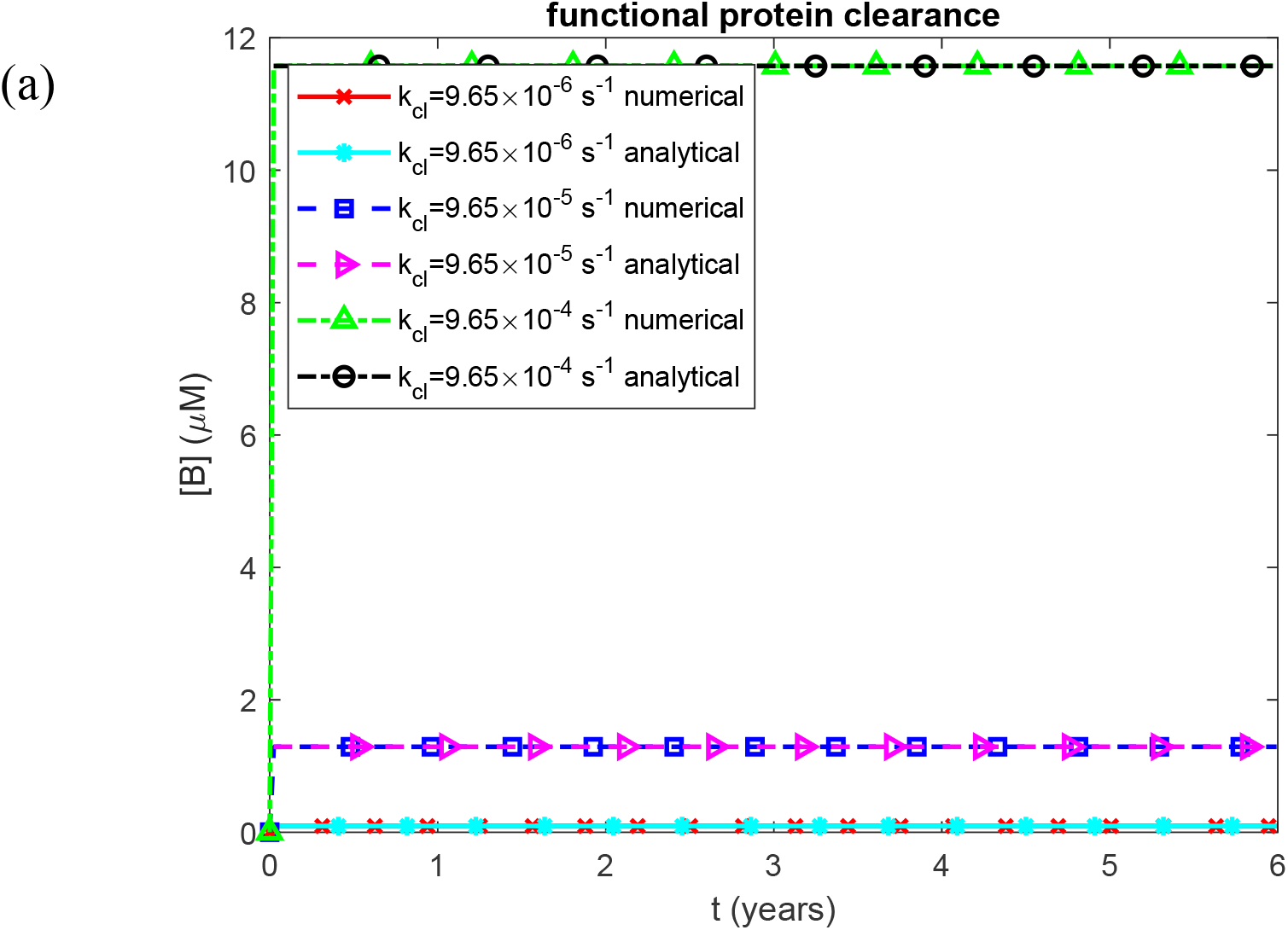
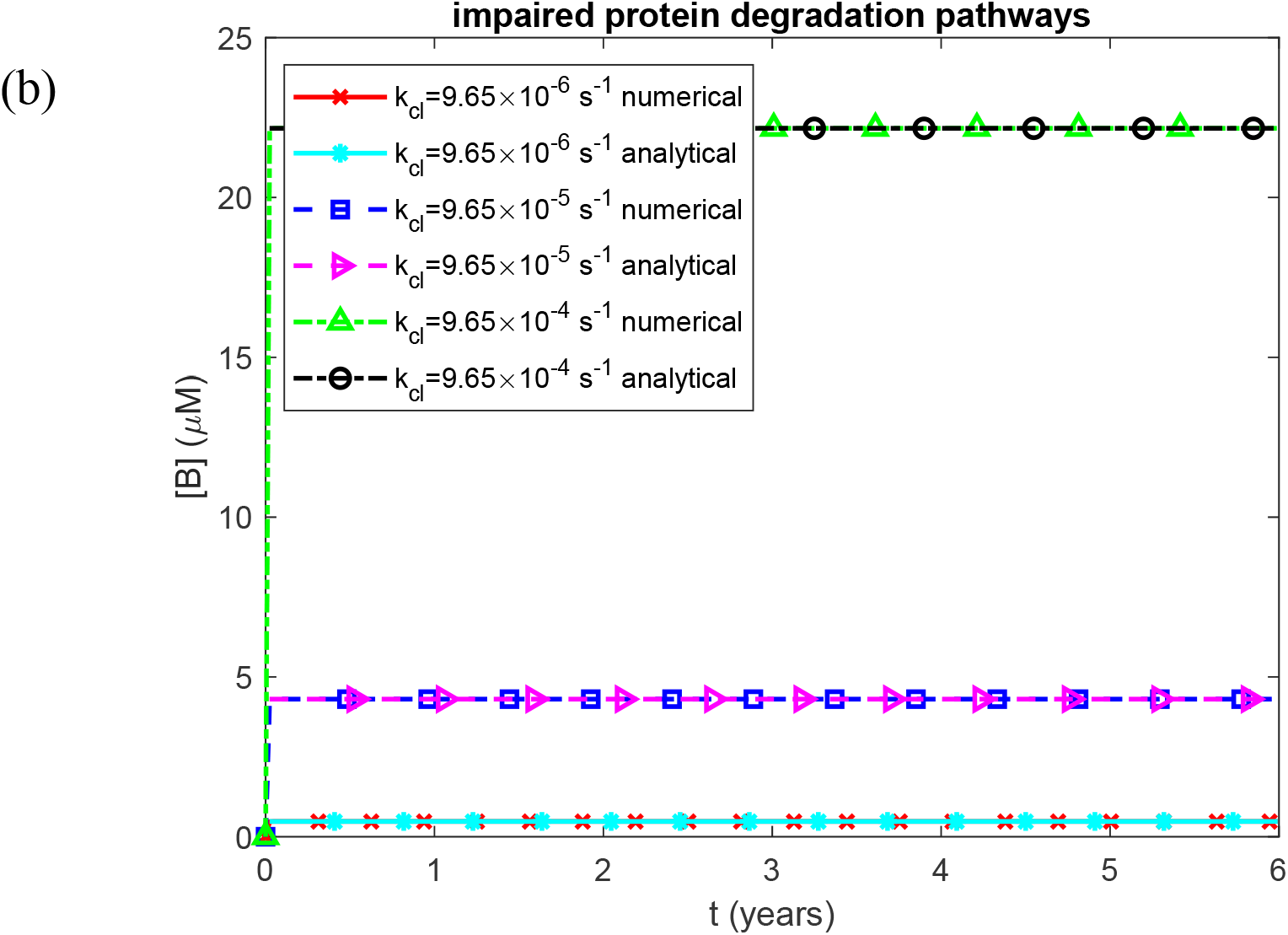
Molar concentration of free (non-deposited) AA oligomers in the kidneys as a function of time. (a) Biologically relevant half-lives *T*_1/2, *A*_, *T*_1/2,*B*_, *T*_1/2,*I*_, and *T*_1/2,*S*_, representing functional protein degradation machinery. (b) Infinite half-lives *T*_1/2, *A*_, *T*_1/2,*B*_, *T*_1/2,*I*_, and *T*_1/2,*S*_, representing complete impairment of protein degradation. Each panel shows results for three values of the rate constant for proteolytic cleavage of SAA that generates AA monomers. Baseline parameters: *k*_1_ =10^−4^ s^−1^, *k*_2_ = 10^−4^ µM^−1^ s^−1^, *K*_*d*_ = 1.1789 *µ*M, *h*_*S*_ = 5.55×10^−7^ m s^−1^, *H*_*tot*_ = 30 *µ*M, *S*_*tot*_ = 8 *µ*M, and *θ*_1/2,*B*_ = 8.64 ×10^4^ s; remaining parameters as listed in Table 2.

**Fig. S11.**
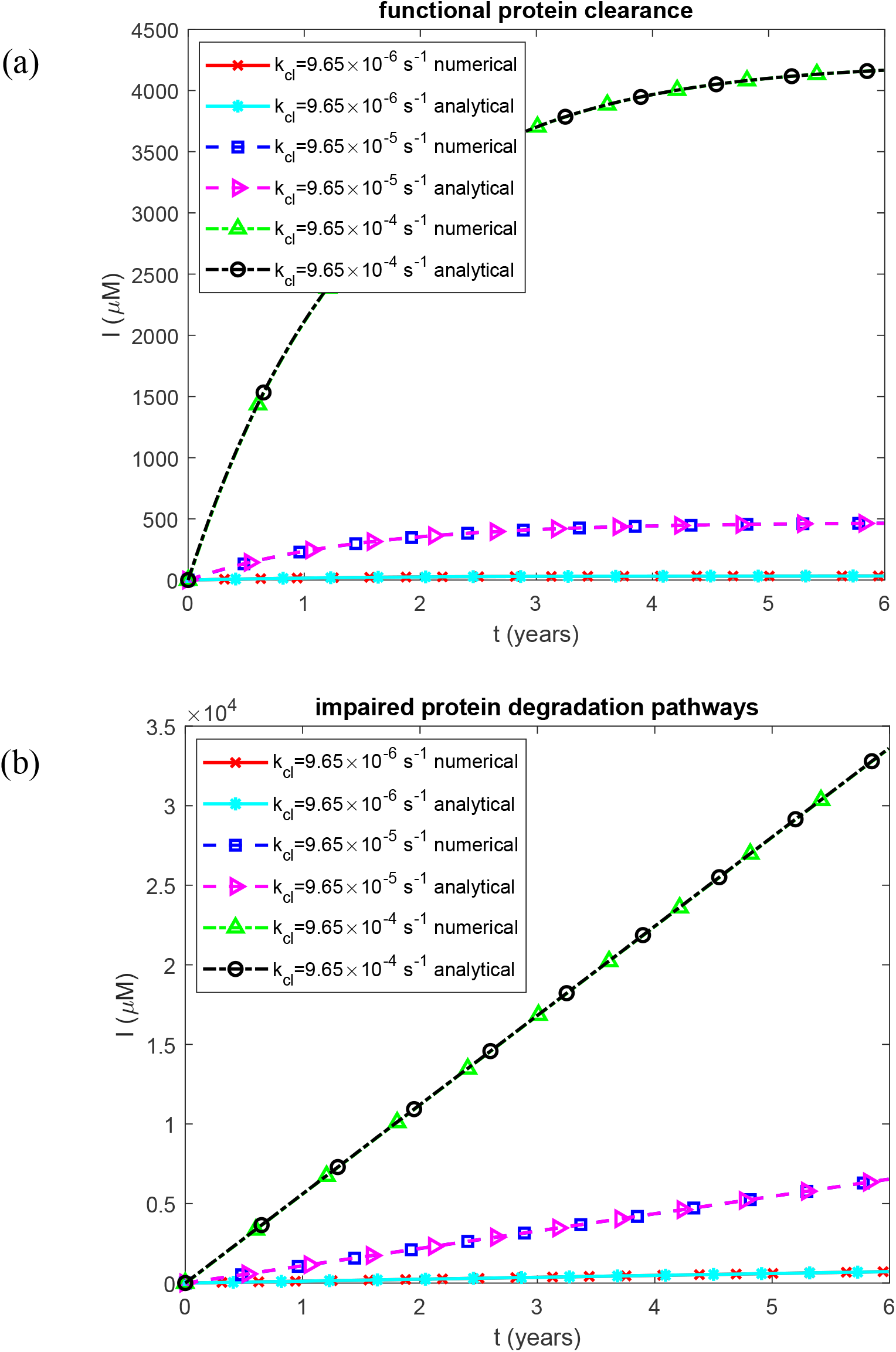
Molar concentration of AA oligomers deposited into AA amyloid fibrils in kidneys as a function of time. (a) Biologically relevant half-lives *T*_1/2, *A*_, *T*_1/2,*B*_, *T*_1/2,*I*_, and *T*_1/2,*S*_, representing functional protein degradation machinery. (b) Infinite half-lives *T*_1/2, *A*_, *T*_1/2,*B*_, *T*_1/2,*I*_, and *T*_1/2,*S*_, representing complete impairment of protein degradation. Each panel shows results for three values of the rate constant for proteolytic cleavage of SAA that generates AA monomers. Baseline parameters: *k*_1_ =10^−4^ s^−1^, *k*_2_ = 10^−4^ µM^−1^ s^−1^, *K*_*d*_ = 1.1789 *µ*M, *h*_*S*_ = 5.55×10^−7^ m s^−1^, *H*_*tot*_ = 30 *µ*M, *S*_*tot*_ = 8 *µ*M, and *θ*_1/2,*B*_ = 8.64 ×10^4^ s; remaining parameters as listed in Table 2.

**Fig. S12.**
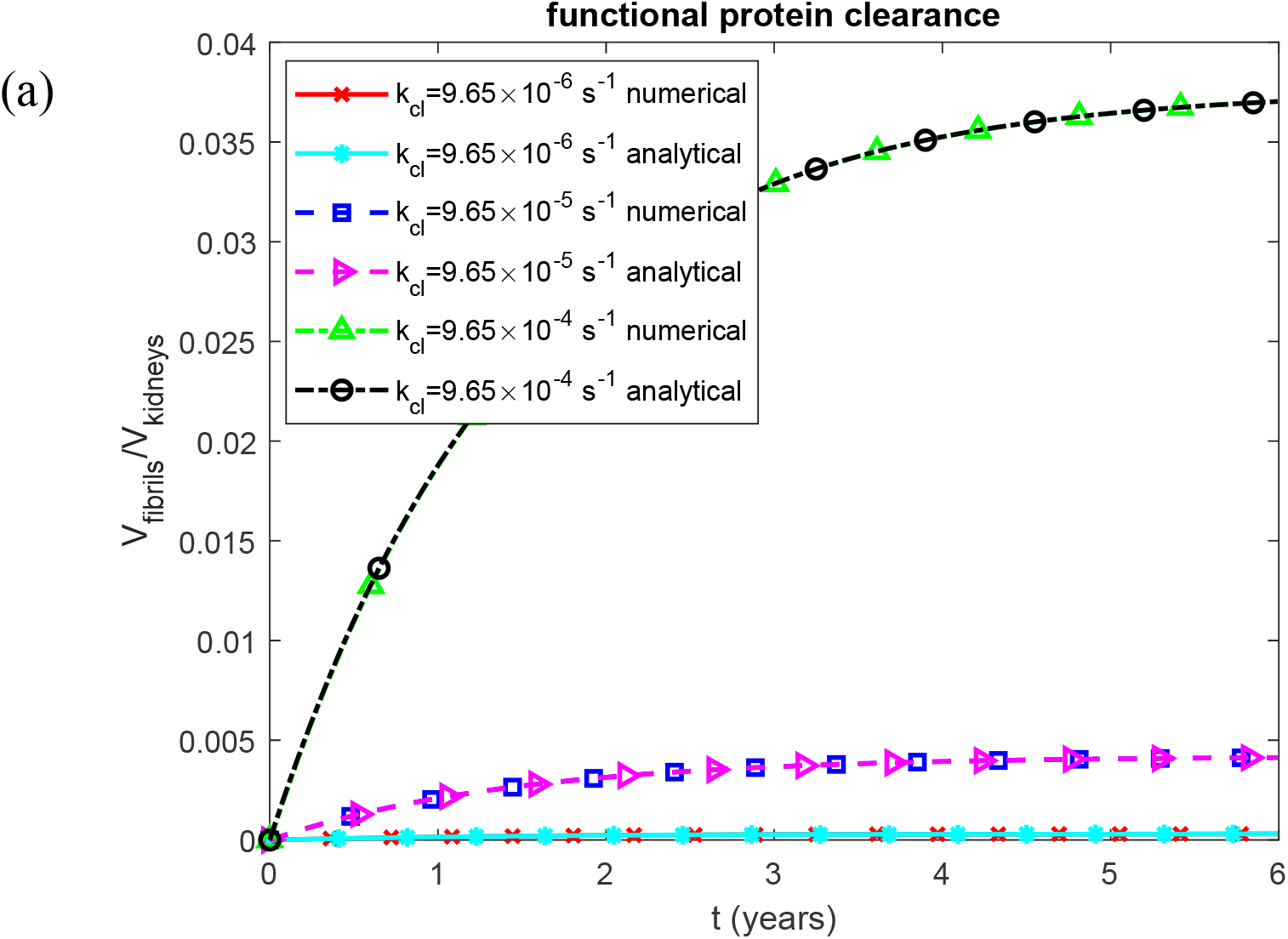
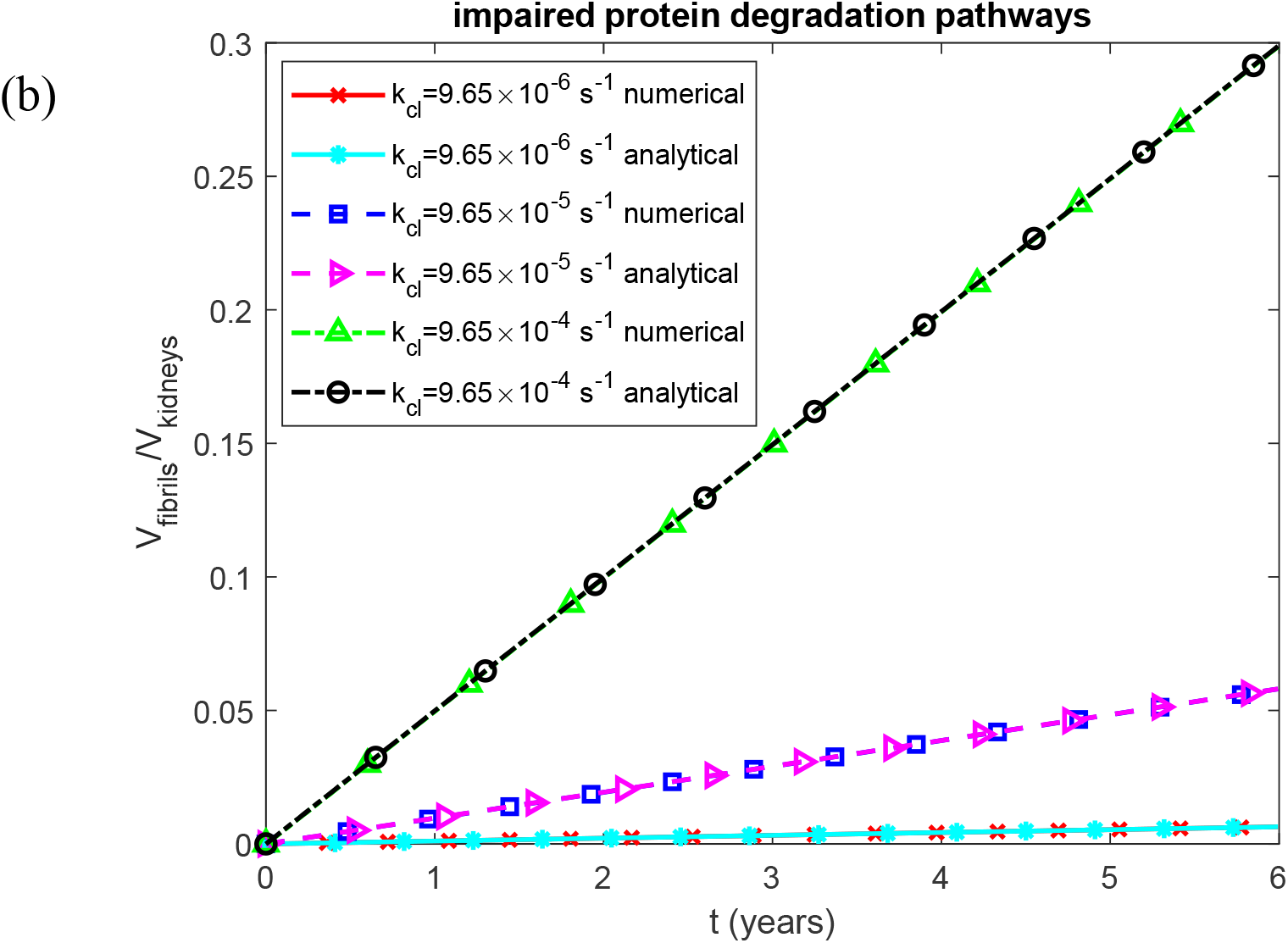
Fraction of the baseline kidney volume occupied by fibrils as a function of time. (a) Biologically relevant half-lives *T*_1/2, *A*_, *T*_1/2,*B*_, *T*_1/2,*I*_, and *T*_1/2,*S*_, representing functional protein degradation machinery. (b) Infinite half-lives *T*_1/2, *A*_, *T*_1/2,*B*_, *T*_1/2,*I*_, and *T*_1/2,*S*_, representing complete impairment of protein degradation. Each panel shows results for three values of the rate constant for proteolytic cleavage of SAA that generates AA monomers. Baseline parameters: *k*_1_ =10^−4^ s^−1^, *k*_2_ = 10^−4^ µM^−1^ s^−1^, *K*_*d*_ = 1.1789 *µ*M, *h*_*S*_ = 5.55×10^−7^ m s^−1^, *H*_*tot*_ = 30 *µ*M, *S*_*tot*_ = 8 *µ*M, and *θ*_1/2,*B*_ = 8.64 ×10^4^ s; remaining parameters as listed in Table 2.

**Fig. S13.**
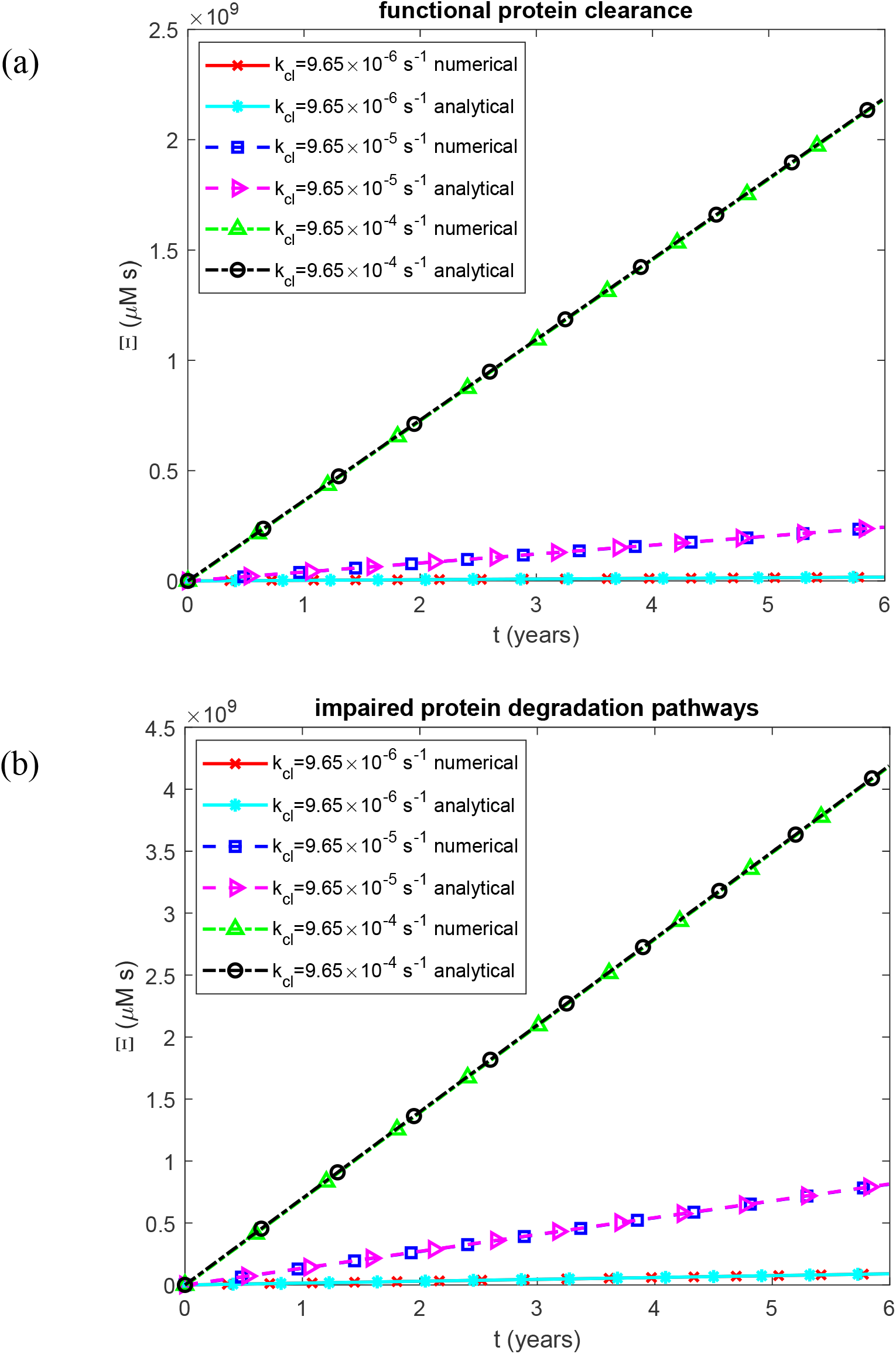
Accumulated nephrotoxicity of AA oligomers as a function of time. (a) Biologically relevant half-lives *T*_1/2, *A*_, *T*_1/2,*B*_, *T*_1/2,*I*_, and *T*_1/2,*S*_, representing functional protein degradation machinery. (b) Infinite half-lives *T*_1/2, *A*_, *T*_1/2,*B*_, *T*_1/2,*I*_, and *T*_1/2,*S*_, representing complete impairment of protein degradation. Each panel shows results for three values of the rate constant for proteolytic cleavage of SAA that generates AA monomers. Baseline parameters: *k*_1_ =10^−4^ s^−1^, *k*_2_ = 10^−4^ µM^−1^ s^−1^, *K*_*d*_ = 1.1789 *µ*M, *h*_*S*_ = 5.55×10^−7^ m s^−1^, *H*_*tot*_ = 30 *µ*M, *S*_*tot*_ = 8 *µ*M, and *θ*_1/2,*B*_ = 8.64 ×10^4^ s; remaining parameters as listed in Table 2.

**Fig. S14.**
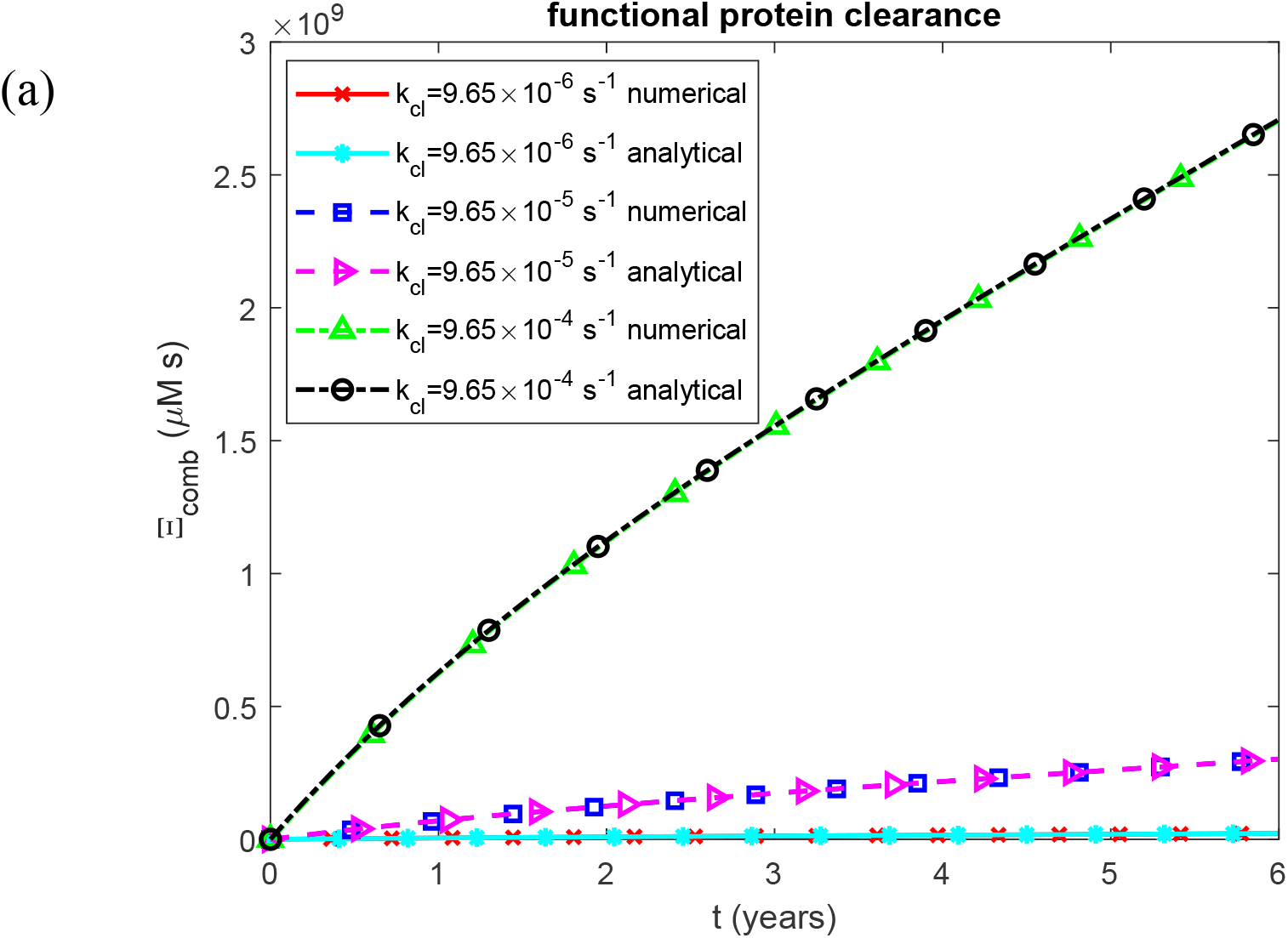
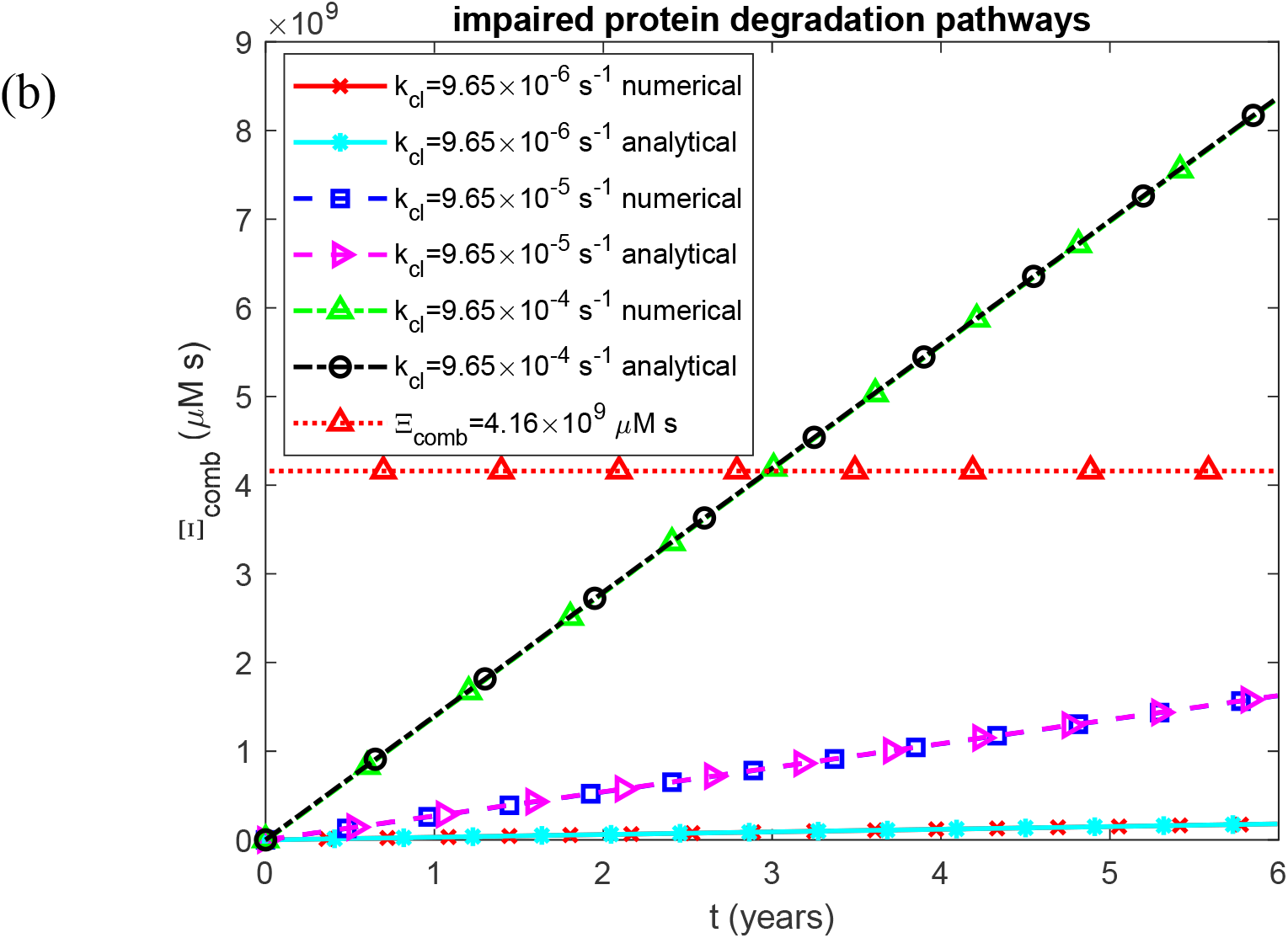
Combined renal damage from accumulated AA oligomer cytotoxicity and insoluble fibril burden as a function of time. (a) Biologically relevant half-lives *T*_1/2, *A*_, *T*_1/2,*B*_, *T*_1/2,*I*_, and *T*_1/2,*S*_, representing functional protein degradation machinery. (b) Infinite half-lives *T*_1/2, *A*_, *T*_1/2,*B*_, *T*_1/2,*I*_, and *T*_1/2,*S*_, representing complete impairment of protein degradation. Each panel shows results for three values of the rate constant for proteolytic cleavage of SAA that generates AA monomers. The horizontal line in (b) marks *Ξ*_*comb*_ = 4.16 × 10^9^ µM s, the combined damage that corresponds to the calibrated terminal-damage threshold of BA = 100 years (Section 2.3). Baseline parameters: *k*_1_ =10^−4^ s^−1^, *k*_2_ = 10^−4^ µM^−1^ s^−1^, *K*_*d*_ = 1.1789 *µ*M, *h*_*S*_ = 5.55×10^−7^ m s^−1^, *H*_*tot*_ = 30 *µ*M, *S*_*tot*_ = 8 *µ*M, and *θ*_1/2,*B*_ = 8.64 ×10^4^ s; remaining parameters as listed in Table 2.

**Fig. S15.**
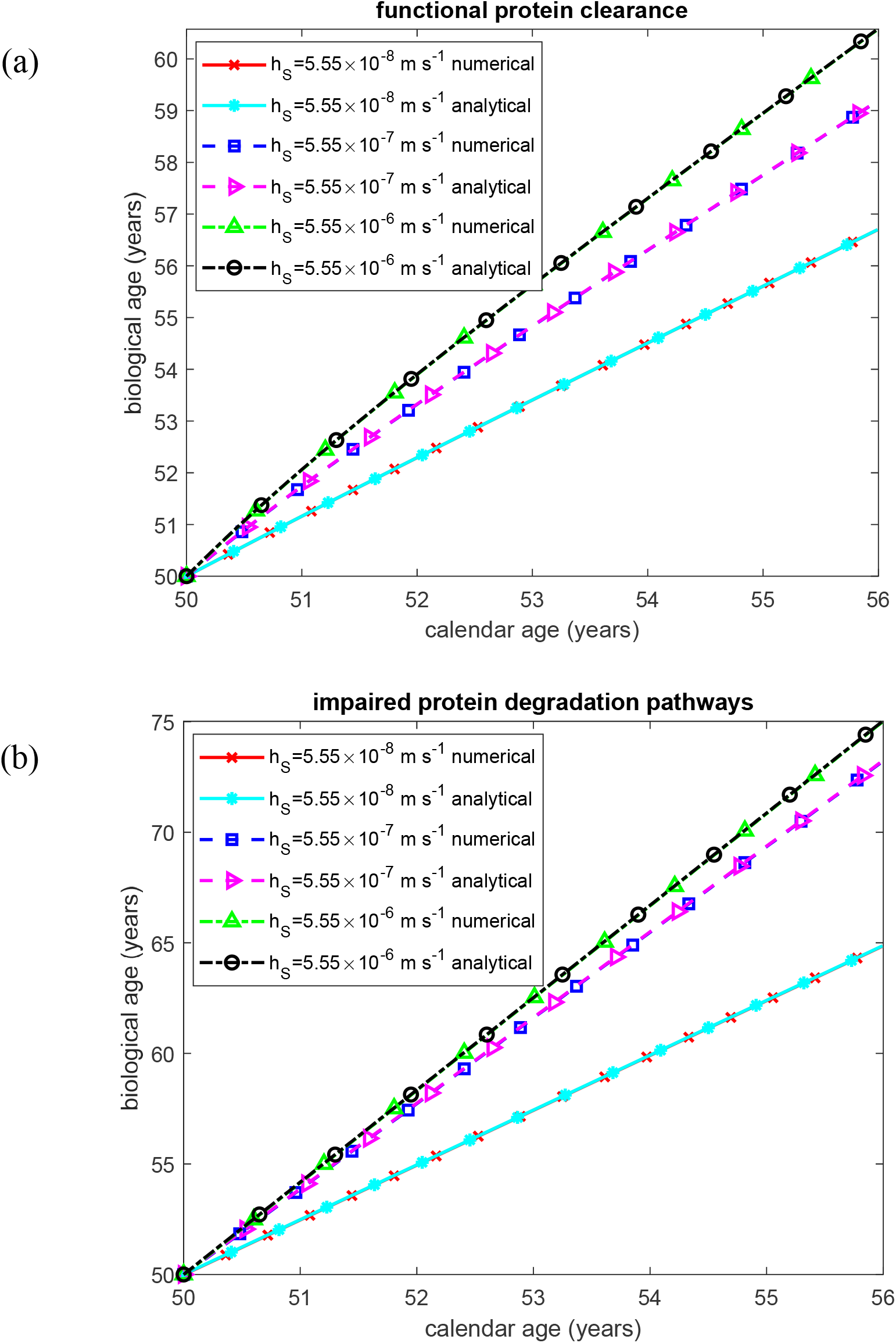
Biological age of the kidney as a function of time. (a) Biologically relevant half-lives *T*_1/2, *A*_, *T*_1/2,*B*_, *T*_1/2,*I*_, and *T*_1/2,*S*_, representing functional protein degradation machinery. (b) Infinite half-lives *T*_1/2, *A*_, *T*_1/2,*B*_, *T*_1/2,*I*_, and *T*_1/2,*S*_, representing complete impairment of protein degradation. Each panel shows results for three values of the mass transfer coefficient characterizing free SAA transport from blood plasma into the kidneys. Baseline parameters: *k*_1_ =10^−4^ s^−1^, *k*_2_ = 10^−4^ µM^−1^ s^−1^, *k*_*cl*_ = 9.65×10^−5^ s^−1^, *K*_*d*_ = 1.1789 *µ*M, *H*_*tot*_ = 30 *µ*M, *S*_*tot*_ = 8 *µ*M, and *θ*_1/2,*B*_ = 8.64 ×10^4^ s; remaining parameters as listed in Table 2.

**Fig. S16.**
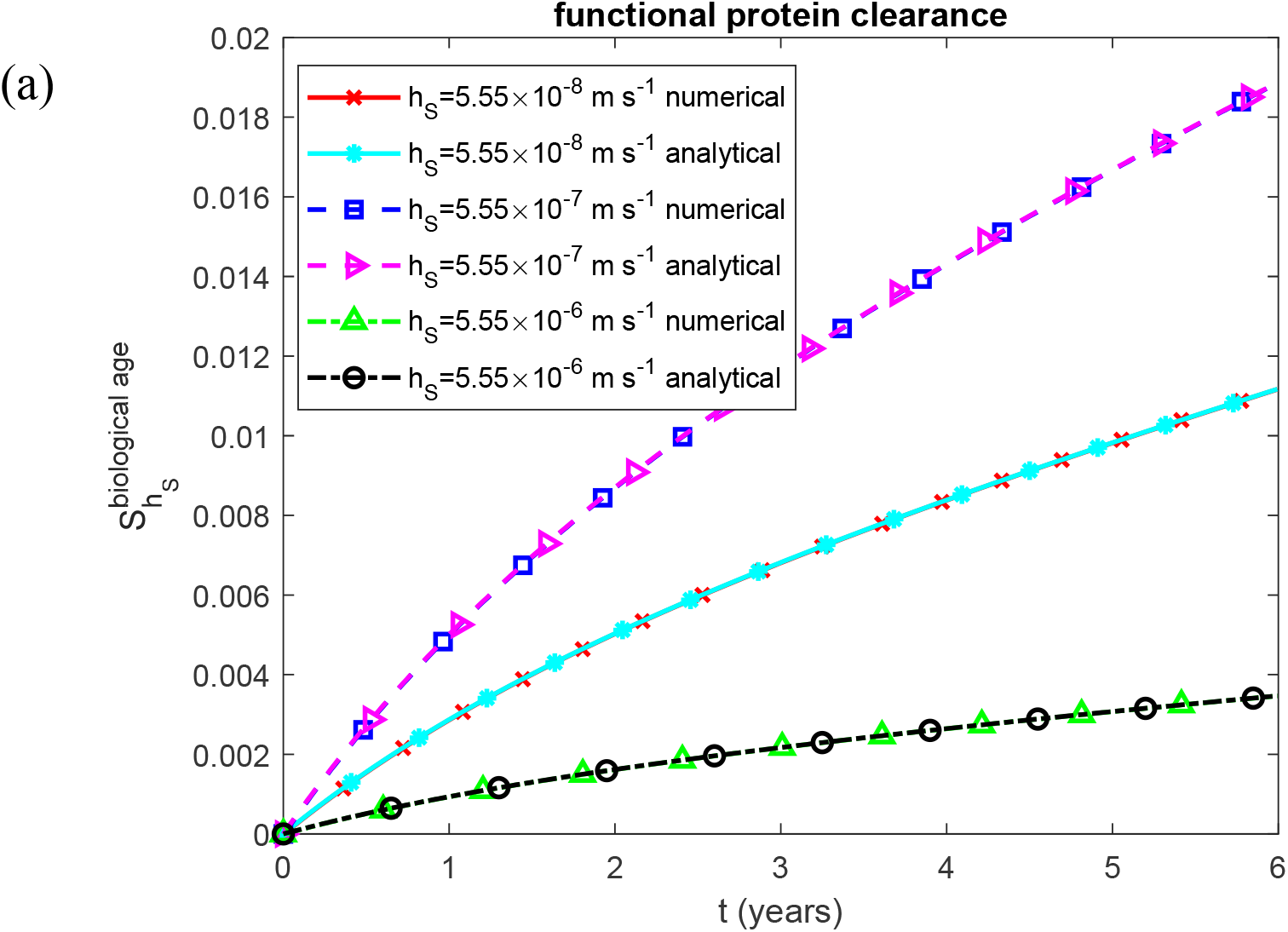
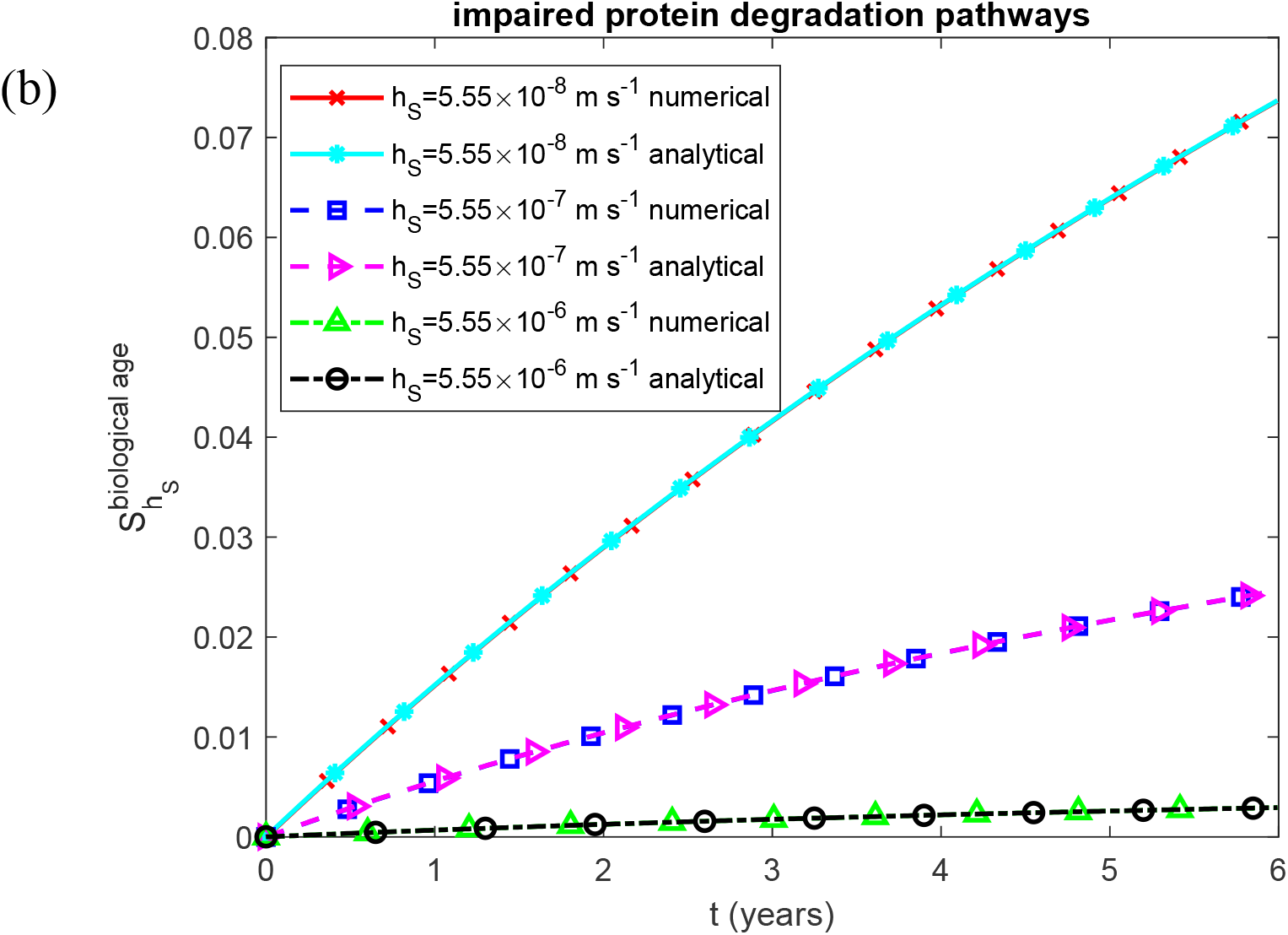
Dimensionless sensitivity of renal biological age to the mass transfer coefficient for free SAA transport from blood plasma into the kidneys, as a function of time. (a) Biologically relevant half-lives *T*_1/2, *A*_, *T*_1/2,*B*_, *T*_1/2,*I*_, and *T*_1/2,*S*_, representing functional protein degradation machinery. (b) Infinite half-lives *T*_1/2, *A*_, *T*_1/2,*B*_, *T*_1/2,*I*_, and *T*_1/2,*S*_, representing complete impairment of protein degradation. Each panel shows results for three values of the mass transfer coefficient characterizing free SAA transport from blood plasma into the kidneys. Baseline parameters: *k*_1_ =10^−4^ s^−1^, *k*_2_ = 10^−4^ µM^−1^ s^−1^, *k*_*cl*_ = 9.65×10^−5^ s^−1^, *K*_*d*_ = 1.1789 *µ*M, *H*_*tot*_ = 30 *µ*M, *S*_*tot*_ = 8 *µ*M, and *θ*_1/2,*B*_ = 8.64 ×10^4^ s; remaining parameters as listed in Table 2.

**Fig. S17.**
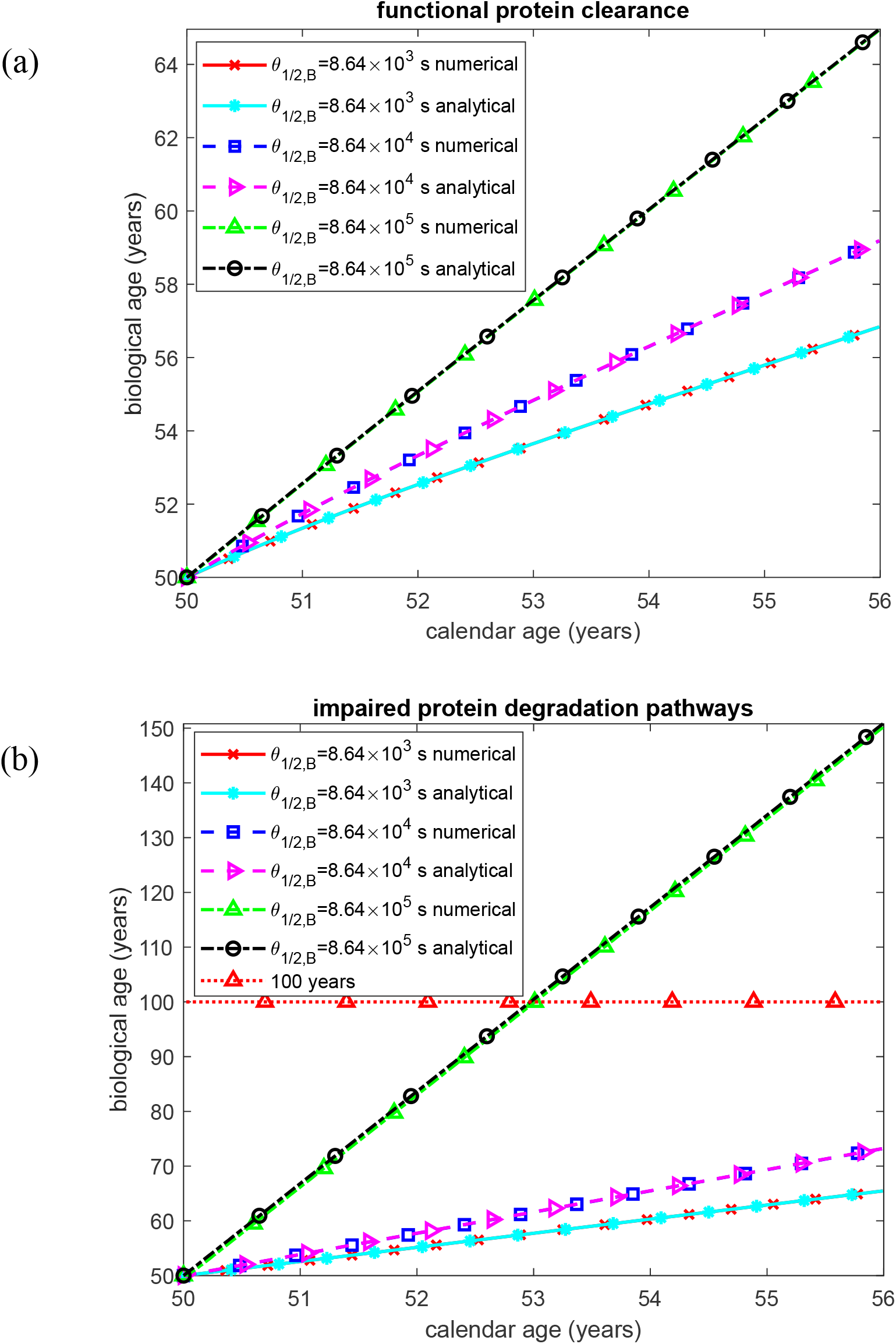
Biological age of the kidney as a function of time. (a) Biologically relevant half-lives *T*_1/2, *A*_, *T*_1/2,*B*_, *T*_1/2,*I*_, and *T*_1/2,*S*_, representing functional protein degradation machinery. (b) Infinite half-lives *T*_1/2, *A*_, *T*_1/2,*B*_, *T*_1/2,*I*_, and *T*_1/2,*S*_, representing complete impairment of protein degradation. Each panel shows results for three values of the half-deposition time characterizing the incorporation of free AA oligomers into fibrillar deposits. The horizontal line in (b) marks the calibrated terminal-damage threshold of BA = 100 years. Baseline parameters: *k*_1_ =10^−4^ s^−1^, *k*_2_ = 10^−4^ µM^−1^ s^−1^, *k*_*cl*_ = 9.65×10^−5^ s^−1^, *K*_*d*_ = 1.1789 *µ*M, *h*_*S*_ = 5.55×10^−7^ m s^−1^, *H*_*tot*_ = 30 *µ*M, and *S*_*tot*_ = 8 *µ*M ; remaining parameters as listed in Table 2.

**Fig. S18.**
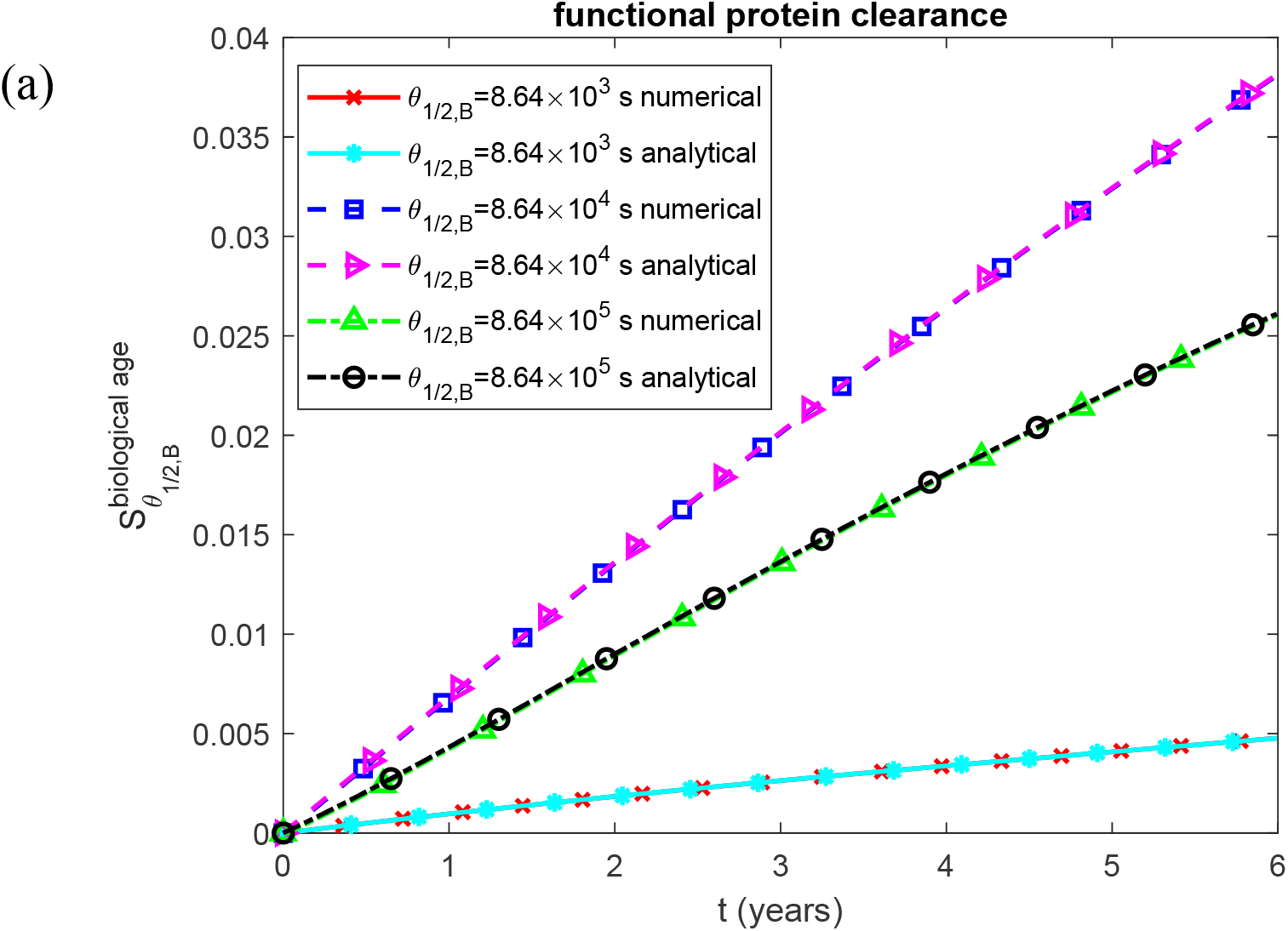
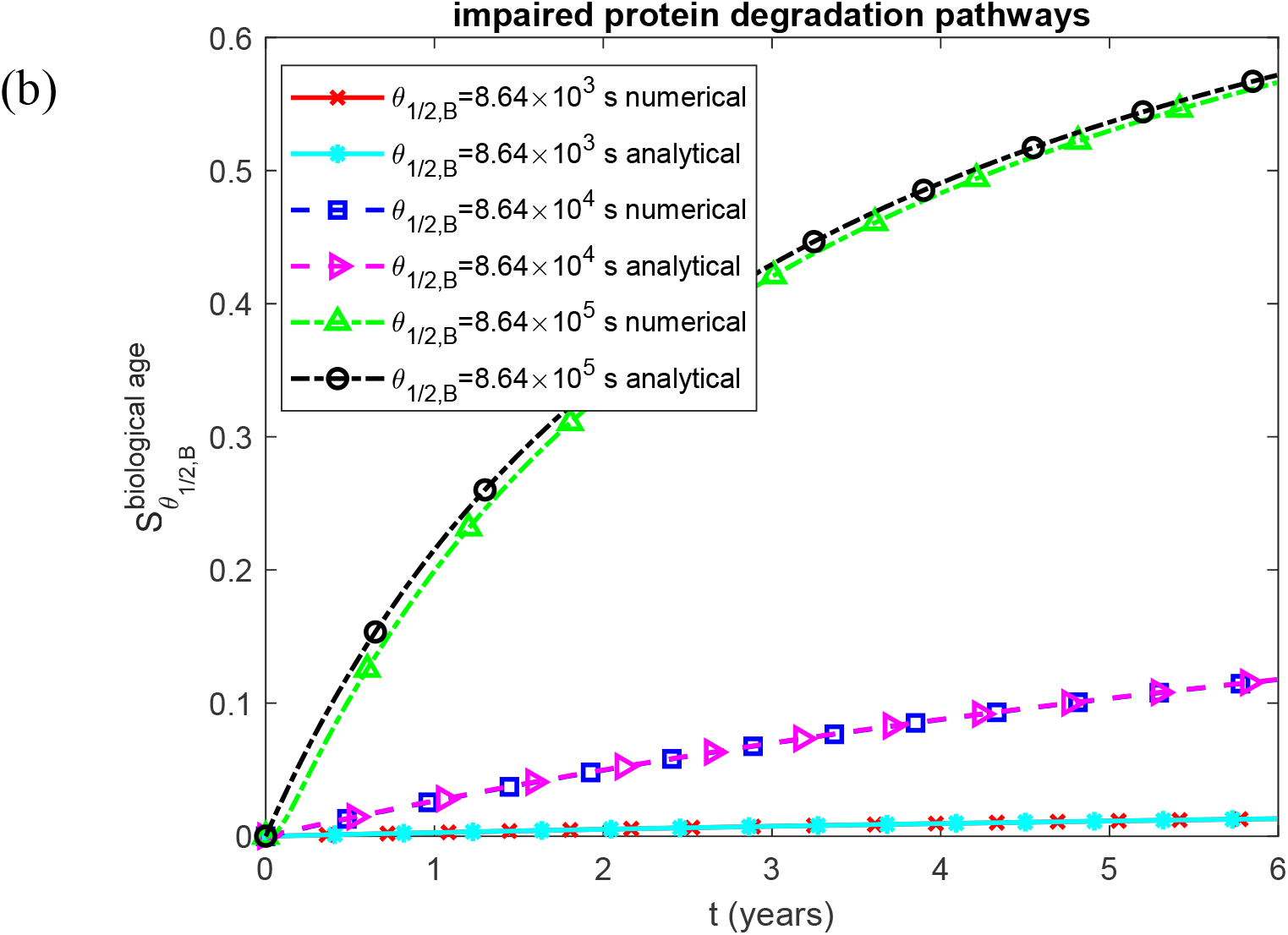
Dimensionless sensitivity of renal biological age to the half-deposition time characterizing the incorporation of free AA oligomers into fibrillar deposits, as a function of time. (a) Biologically relevant half-lives *T*_1/2, *A*_, *T*_1/2,*B*_, *T*_1/2,*I*_, and *T*_1/2,*S*_, representing functional protein degradation machinery. (b) Infinite half-lives *T*_1/2, *A*_, *T*_1/2,*B*_, *T*_1/2,*I*_, and *T*_1/2,*S*_, representing complete impairment of protein degradation. Each panel shows results for three values of the half-deposition time characterizing the incorporation of free AA oligomers into fibrillar deposits. Baseline parameters: *k*_1_ =10^−4^ s^−1^, *k*_2_ = 10^−4^ µM^−1^ s^−1^, *k*_*cl*_ = 9.65×10^−5^ s^−1^, *K*_*d*_ = 1.1789 *µ*M, *h*_*S*_ = 5.55×10^−7^ m s^−1^, *H*_*tot*_ = 30 *µ*M, and *S*_*tot*_ = 8 *µ*M ; remaining parameters as listed in Table 2.

**Fig. S19.**
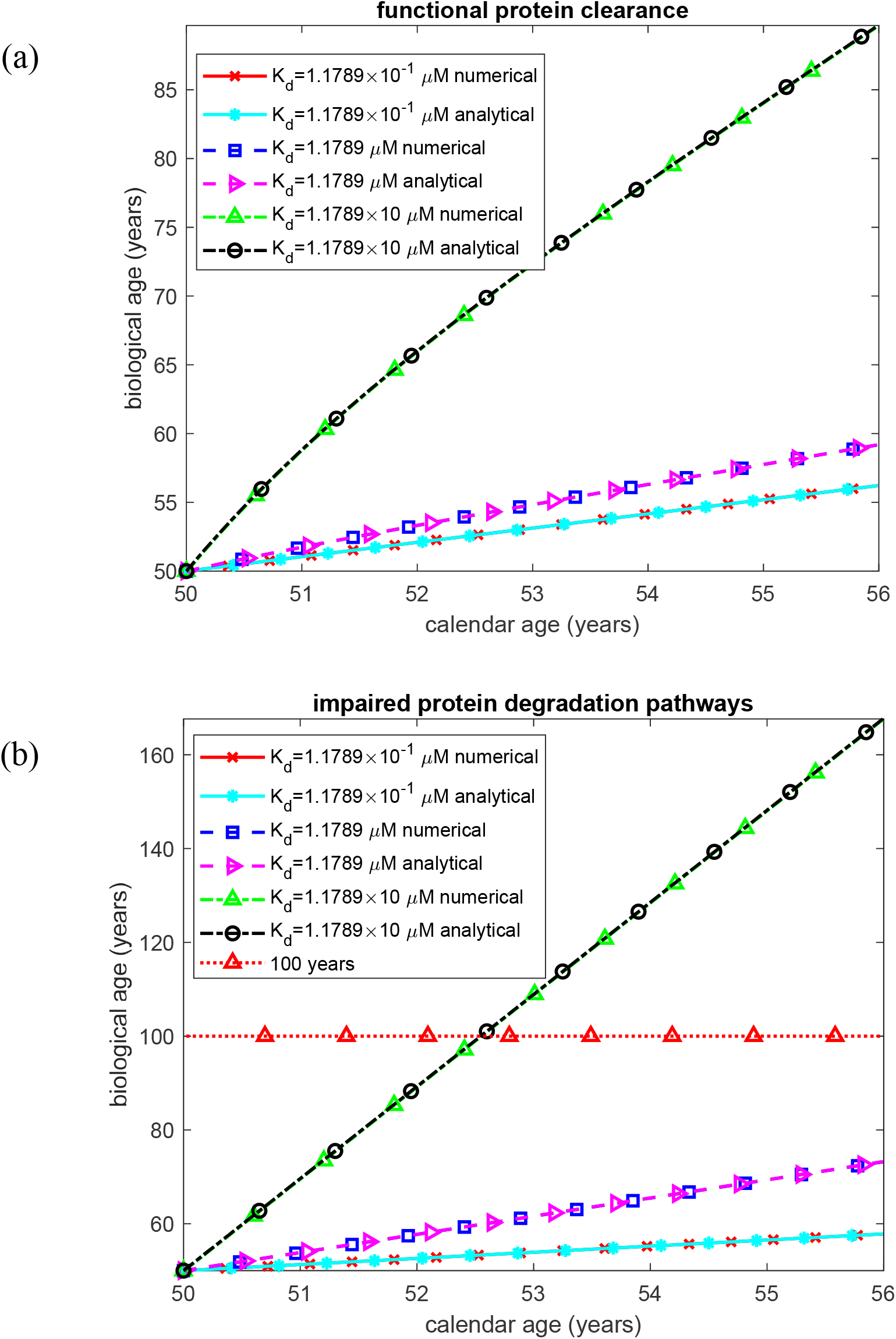
Biological age of the kidney as a function of time. (a) Biologically relevant half-lives *T*_1/2, *A*_, *T*_1/2,*B*_, *T*_1/2,*I*_, and *T*_1/2,*S*_, representing functional protein degradation machinery. (b) Infinite half-lives *T*_1/2, *A*_, *T*_1/2,*B*_, *T*_1/2,*I*_, and *T*_1/2,*S*_, representing complete impairment of protein degradation. Each panel shows results for three values of the equilibrium dissociation constant characterizing the affinity of SAA for HDL. The horizontal line in (b) marks the calibrated terminal-damage threshold of BA = 100 years. Baseline parameters: *k*_1_ =10^−4^ s^−1^, *k*_2_ = 10^−4^ µM^−1^ s^−1^, *k*_*cl*_ = 9.65×10^−5^ s^−1^, *h*_*S*_ = 5.55×10^−7^ m s^−1^, *H*_*tot*_ = 30 *µ*M, *S*_*tot*_ = 8 *µ*M, and *θ*_1/2, *B*_ = 8.64 ×10^4^ s; remaining parameters as listed in Table 2.

**Fig. S20.**
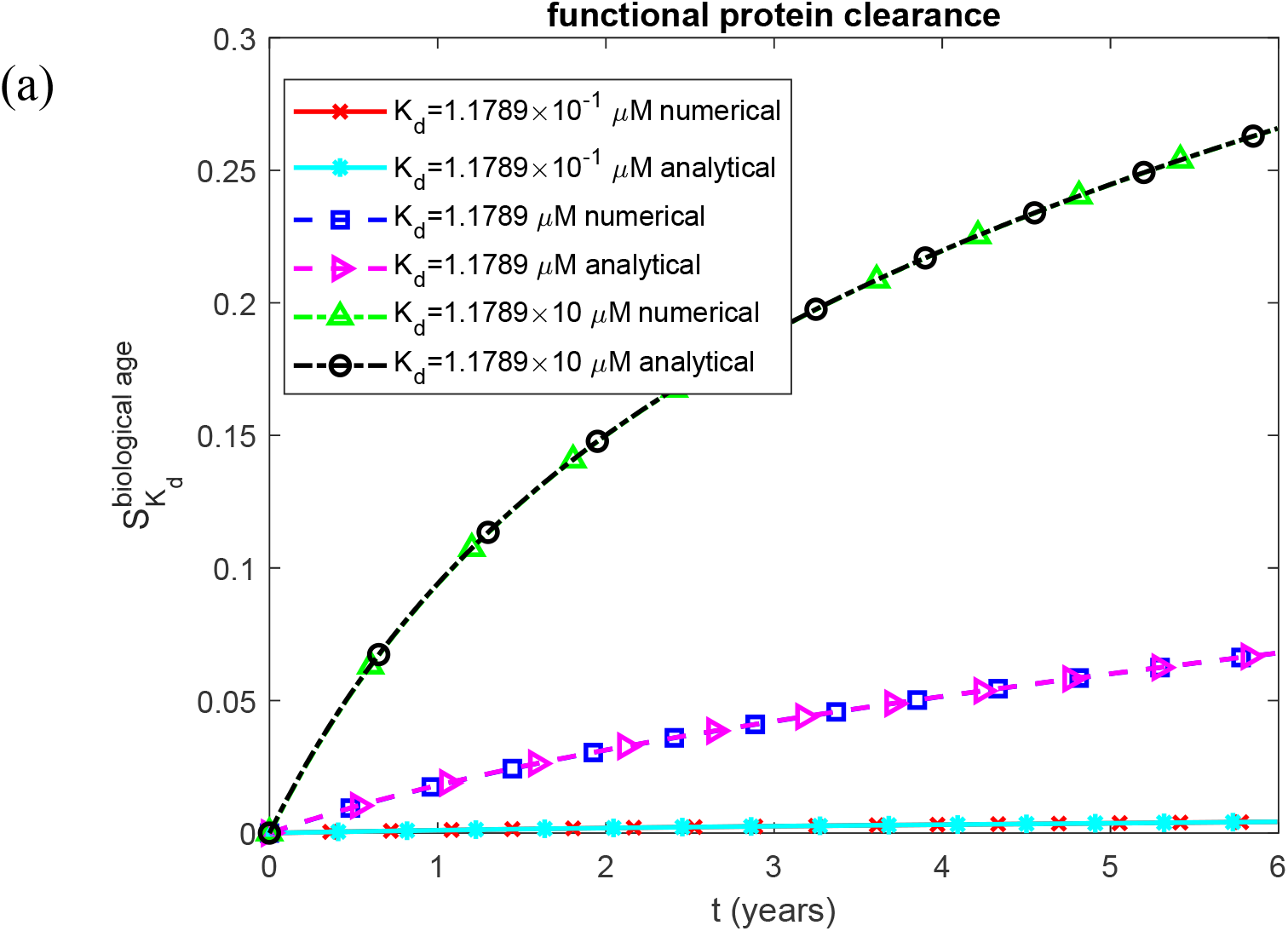
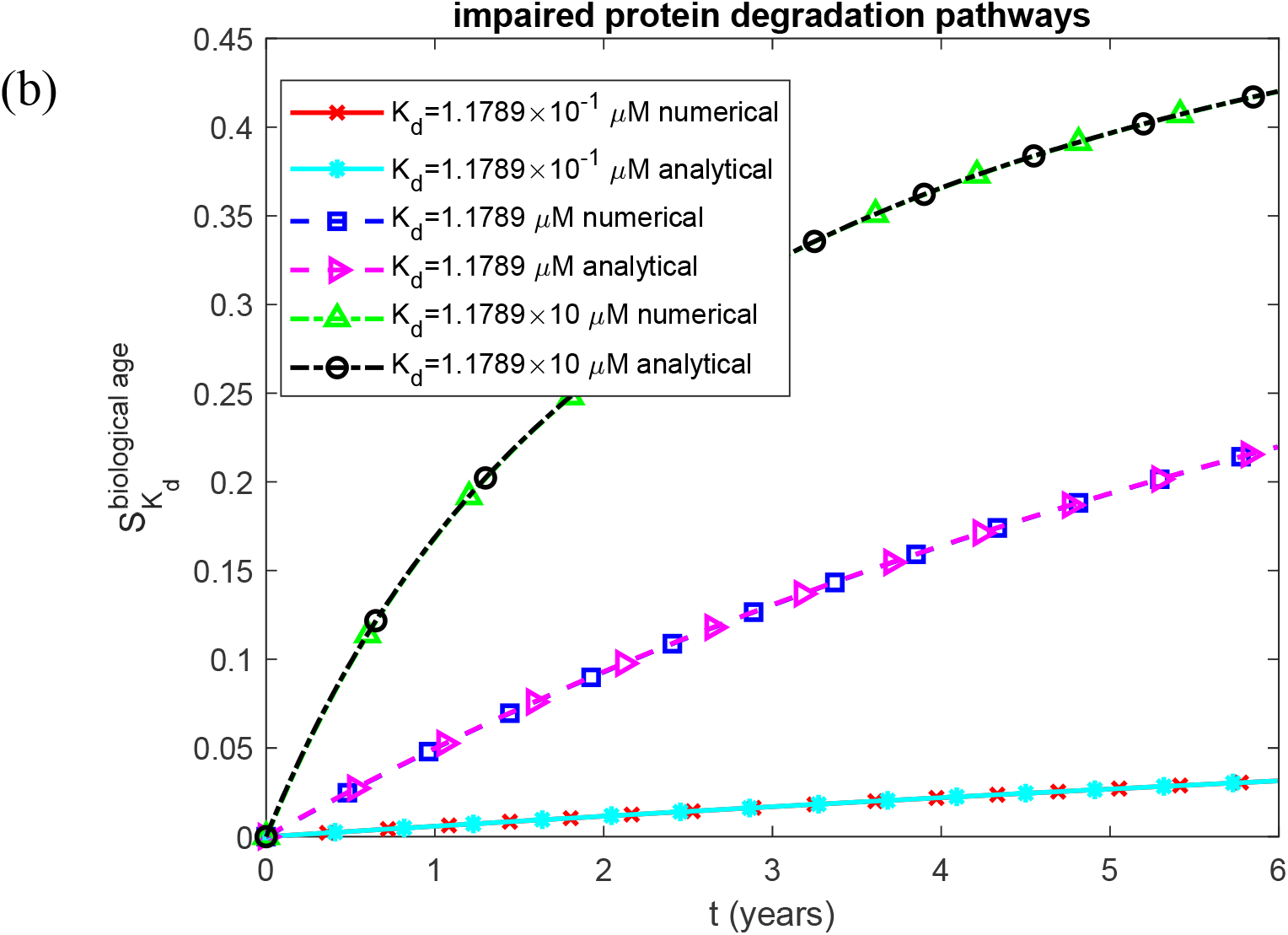
Dimensionless sensitivity of renal biological age to the equilibrium dissociation constant characterizing the affinity of SAA for HDL, as a function of time. (a) Biologically relevant half-lives *T*_1/2, *A*_, *T*_1/2,*B*_, *T*_1/2,*I*_, and *T*_1/2,*S*_, representing functional protein degradation machinery. (b) Infinite half-lives *T*_1/2, *A*_, *T*_1/2,*B*_, *T*_1/2,*I*_, and *T*_1/2,*S*_, representing complete impairment of protein degradation. Each panel shows results for three values of the equilibrium dissociation constant characterizing the affinity of SAA for HDL. Baseline parameters: *k*_1_ =10^−4^ s^−1^, *k*_2_ = 10^−4^ µM^−1^ s^−1^, *k*_*cl*_ = 9.65×10^−5^ s^−1^, *h*_*S*_ = 5.55×10^−7^ m s^−1^, *H*_*tot*_ = 30 *µ*M, *S*_*tot*_ = 8 *µ*M, and *θ*_1/2, *B*_ = 8.64 ×10^4^ s; remaining parameters as listed in Table 2.

**Fig. S21.**
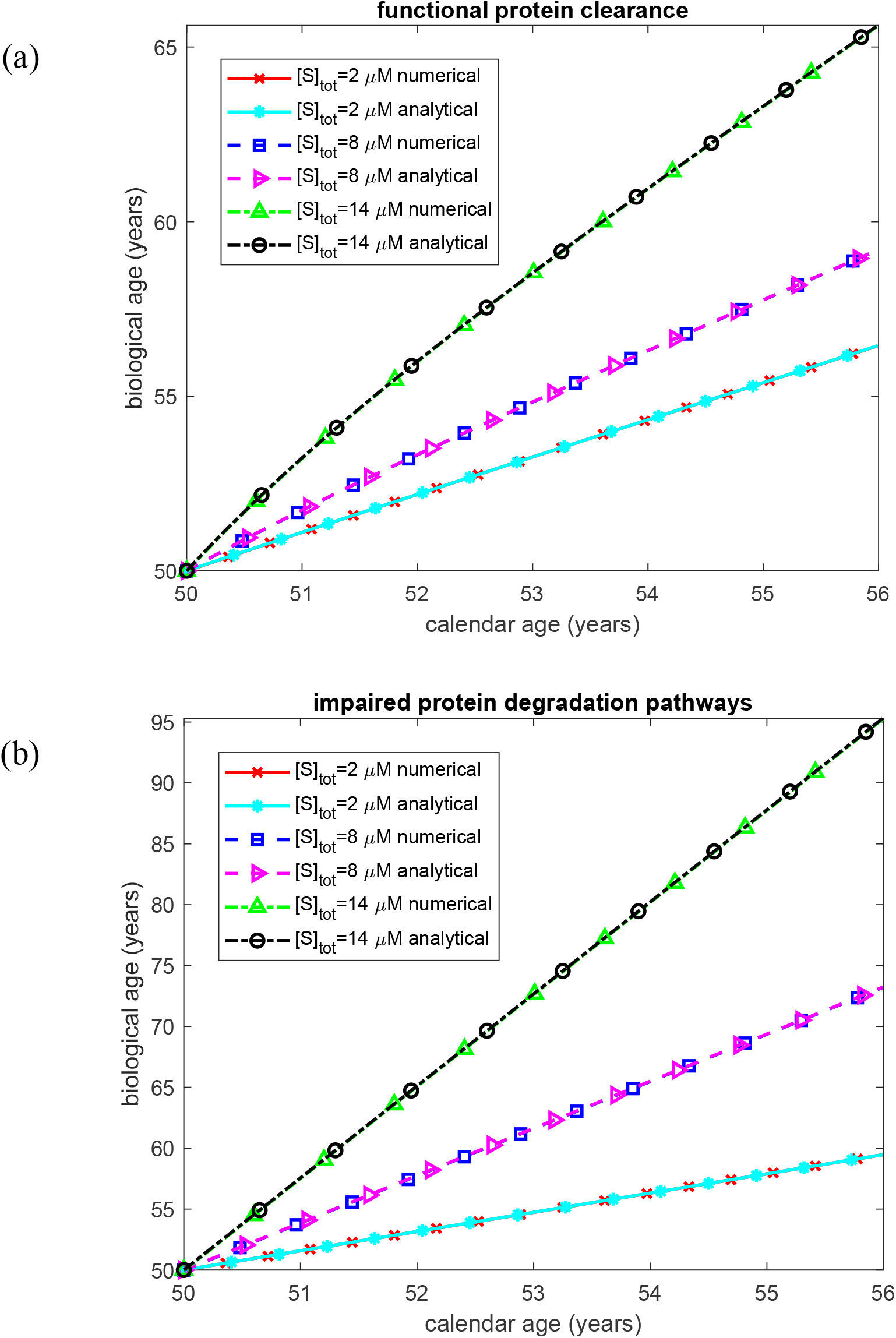
Biological age of the kidney as a function of time. (a) Biologically relevant half-lives *T*_1/2, *A*_, *T*_1/2,*B*_, *T*_1/2,*I*_, and *T*_1/2,*S*_, representing functional protein degradation machinery. (b) Infinite half-lives *T*_1/2, *A*_, *T*_1/2,*B*_, *T*_1/2,*I*_, and *T*_1/2,*S*_, representing complete impairment of protein degradation. Each panel shows results for three values of the total SAA concentration (HDL-bound plus free) under chronic inflammation. Baseline parameters: *k*_1_ =10^−4^ s^−1^, *k*_2_ = 10^−4^ µM^−1^ s^−1^, *k*_*cl*_ = 9.65×10^−5^ s^−1^, *K*_*d*_ = 1.1789 *µ*M, *h*_*S*_ = 5.55×10^−7^ m s^−1^, *H*_*tot*_ = 30 *µ*M, and *θ*_1/2, *B*_ = 8.64 ×10^4^ s; remaining parameters as listed in Table 2.

**Fig. S22.**
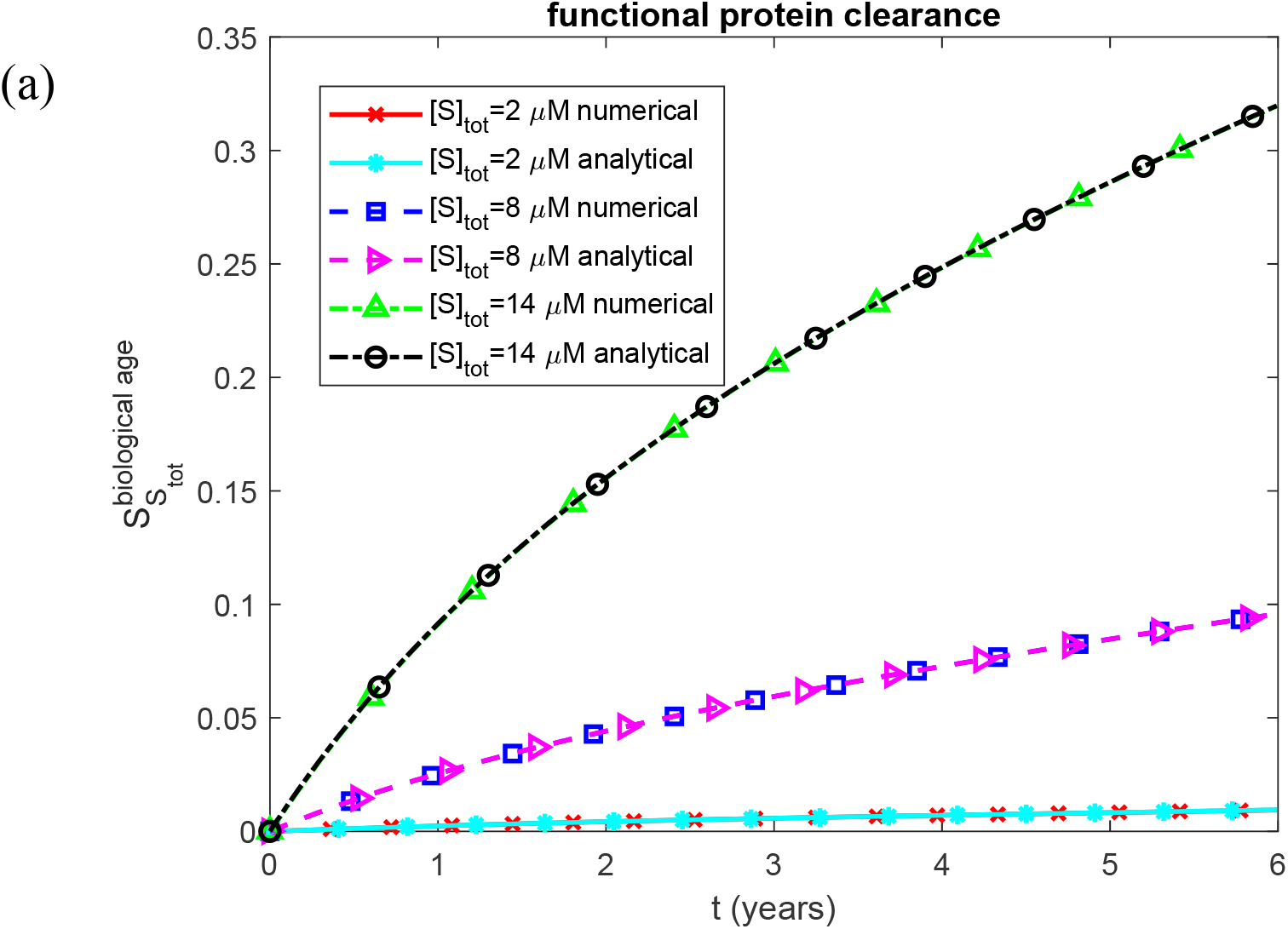
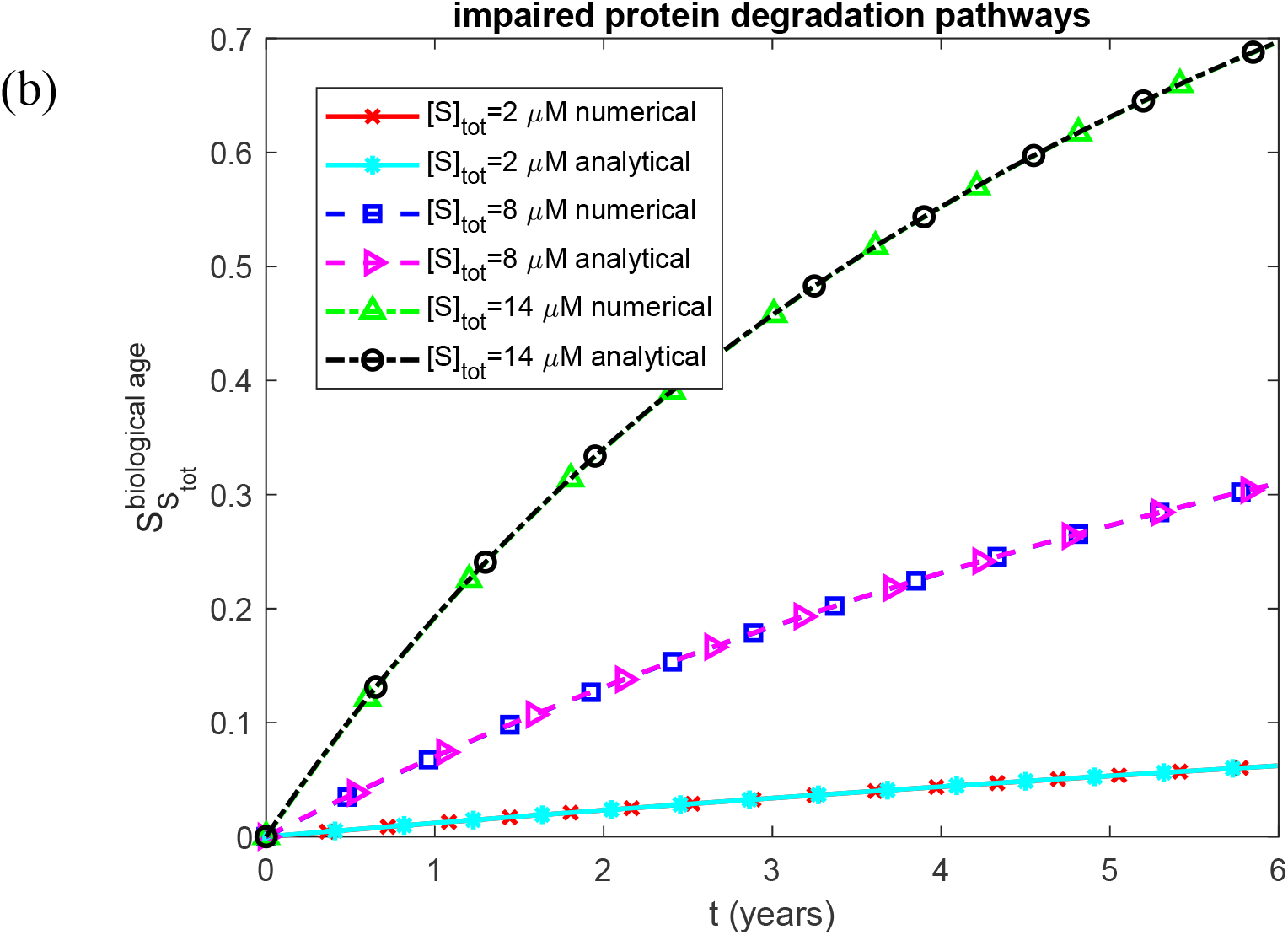
Dimensionless sensitivity of renal biological age to the total SAA concentration (HDL-bound plus free) under chronic inflammation, as a function of time. (a) Biologically relevant half-lives *T*_1/2, *A*_, *T*_1/2,*B*_, *T*_1/2,*I*_, and *T*_1/2,*S*_, representing functional protein degradation machinery. (b) Infinite half-lives *T*_1/2, *A*_, *T*_1/2,*B*_, *T*_1/2,*I*_, and *T*_1/2,*S*_, representing complete impairment of protein degradation. Each panel shows results for three values of the total SAA concentration (HDL-bound plus free) under chronic inflammation. Baseline parameters: *k*_1_ =10^−4^ s^−1^, *k*_2_ = 10^−4^ µM^−1^ s^−1^, *k*_*cl*_ = 9.65×10^−5^ s^−1^, *K*_*d*_ = 1.1789 *µ*M, *h*_*S*_ = 5.55×10^−7^ m s^−1^, *H*_*tot*_ = 30 *µ*M, and *θ*_1/2, *B*_ = 8.64 ×10^4^ s; remaining parameters as listed in Table 2.

**Fig. S23.**
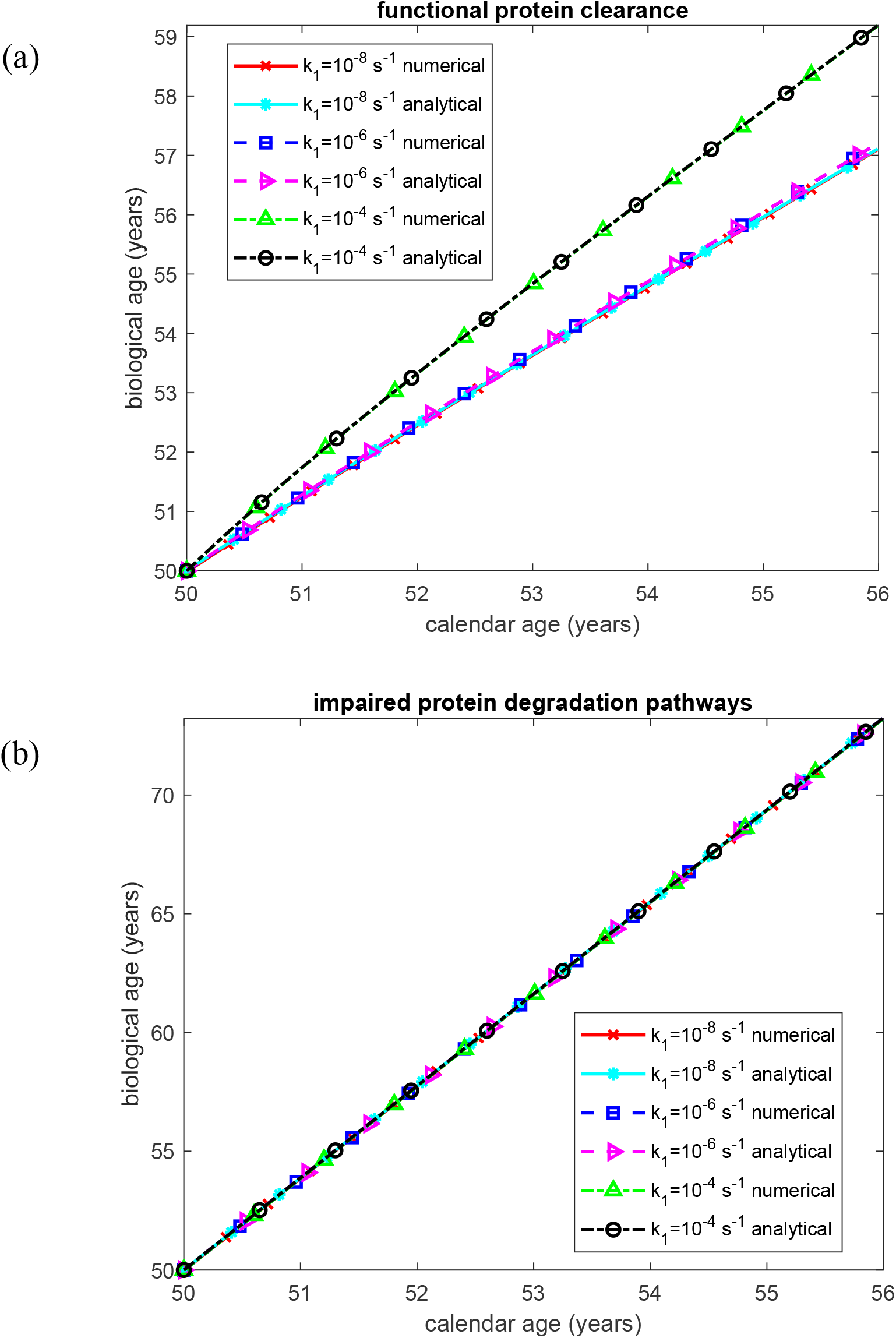
Biological age of the kidney as a function of time. (a) Biologically relevant half-lives *T*_1/2, *A*_, *T*_1/2,*B*_, *T*_1/2,*I*_, and *T*_1/2,*S*_, representing functional protein degradation machinery. (b) Infinite half-lives *T*_1/2, *A*_, *T*_1/2,*B*_, *T*_1/2,*I*_, and *T*_1/2,*S*_, representing complete impairment of protein degradation. Each panel shows results for three values of the rate constant for the first pseudo-elementary (nucleation) step in the F–W model of AA oligomer formation. Baseline parameters: *k*_2_ = 10^−4^ µM^−1^ s^−1^, *k*_*cl*_ = 9.65×10^−5^ s^−1^, *K*_*d*_ = 1.1789 *µ*M, *h*_*S*_ = 5.55×10^−7^ m s^−1^, *H*_*tot*_ = 30 *µ*M, *S*_*tot*_ = 8 *µ*M, and *θ*_1/2,*B*_ = 8.64 ×10^4^ s; remaining parameters as listed in Table 2.

**Fig. S24.**
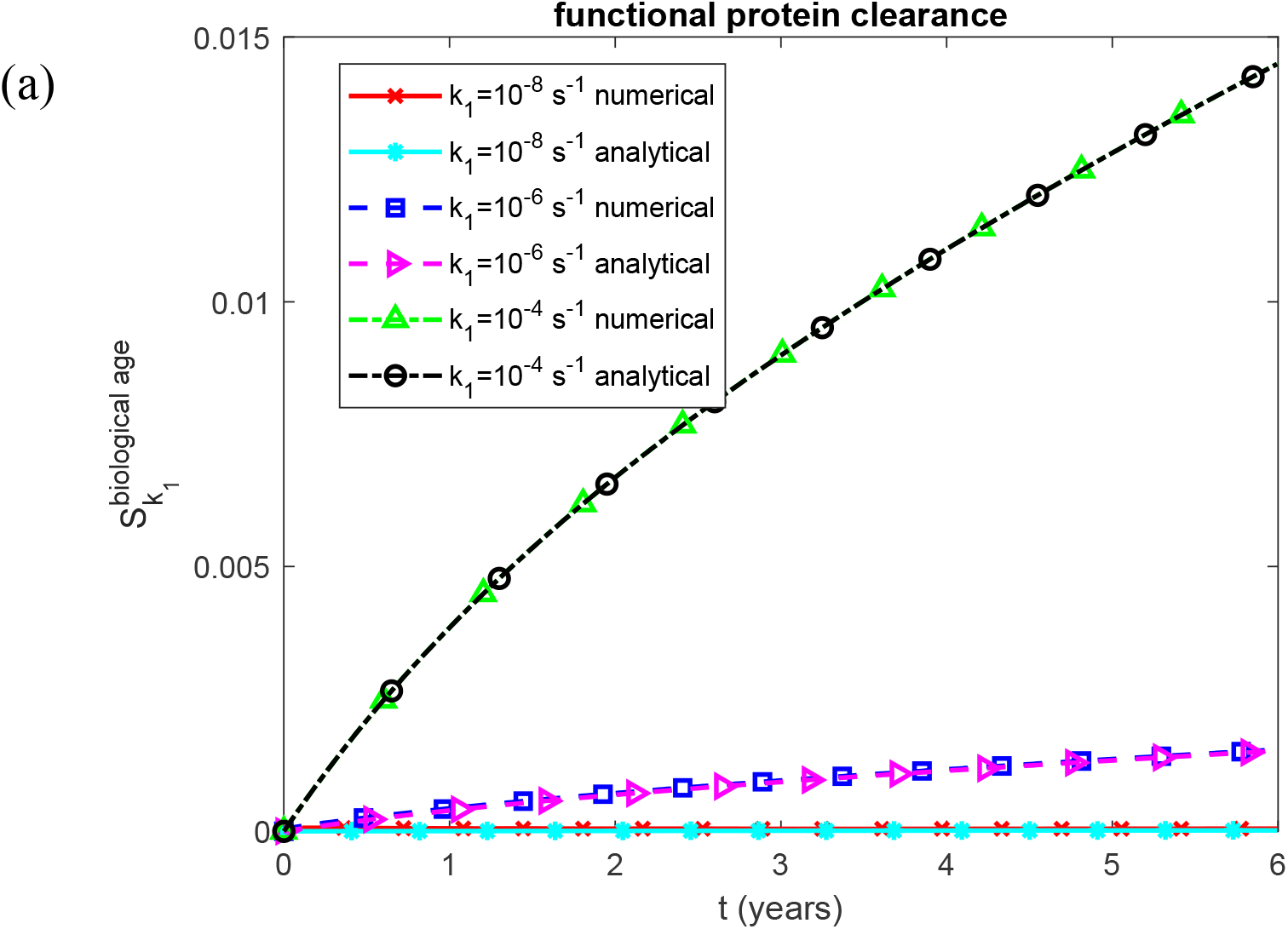
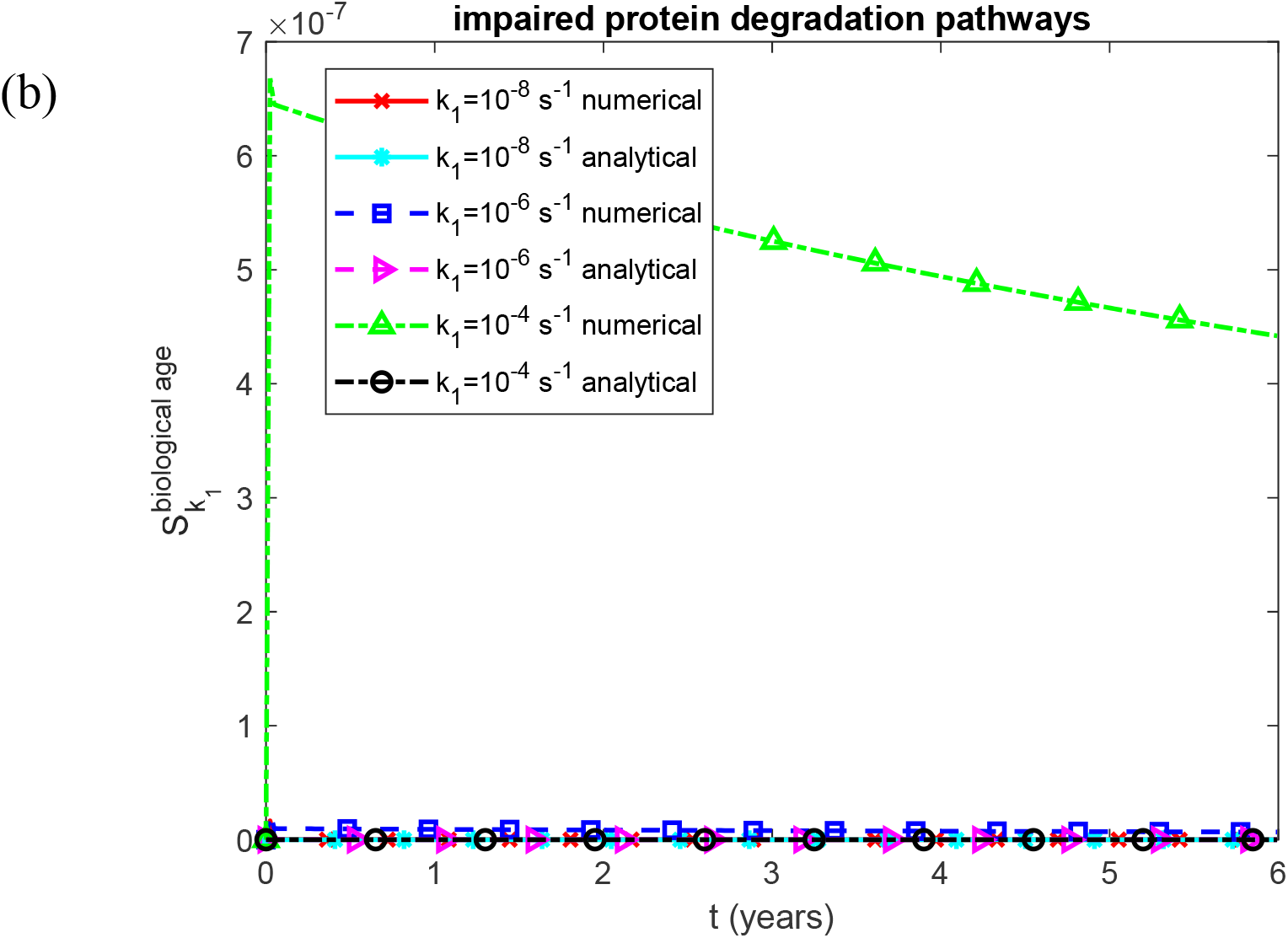
Dimensionless sensitivity of renal biological age to the rate constant for the first pseudo-elementary (nucleation) step in the F–W model of AA oligomer formation, as a function of time. (a) Biologically relevant half-lives *T*_1/2, *A*_, *T*_1/2,*B*_, *T*_1/2,*I*_, and *T*_1/2,*S*_, representing functional protein degradation machinery. (b) Infinite half-lives *T*_1/2, *A*_, *T*_1/2,*B*_, *T*_1/2,*I*_, and *T*_1/2,*S*_, representing complete impairment of protein degradation. Each panel shows results for three values of the rate constant for the first pseudo-elementary (nucleation) step in the F–W model of AA oligomer formation. Baseline parameters: *k*_2_ = 10^−4^ µM^−1^ s^−1^, *k*_*cl*_ = 9.65×10^−5^ s^−1^, *K*_*d*_ = 1.1789 *µ*M, *h*_*S*_ = 5.55×10^−7^ m s^−1^, *H*_*tot*_ = 30 *µ*M, *S*_*tot*_ = 8 *µ*M, and *θ*_1/2,*B*_ = 8.64 ×10^4^ s; remaining parameters as listed in Table 2.

**Fig. S25.**
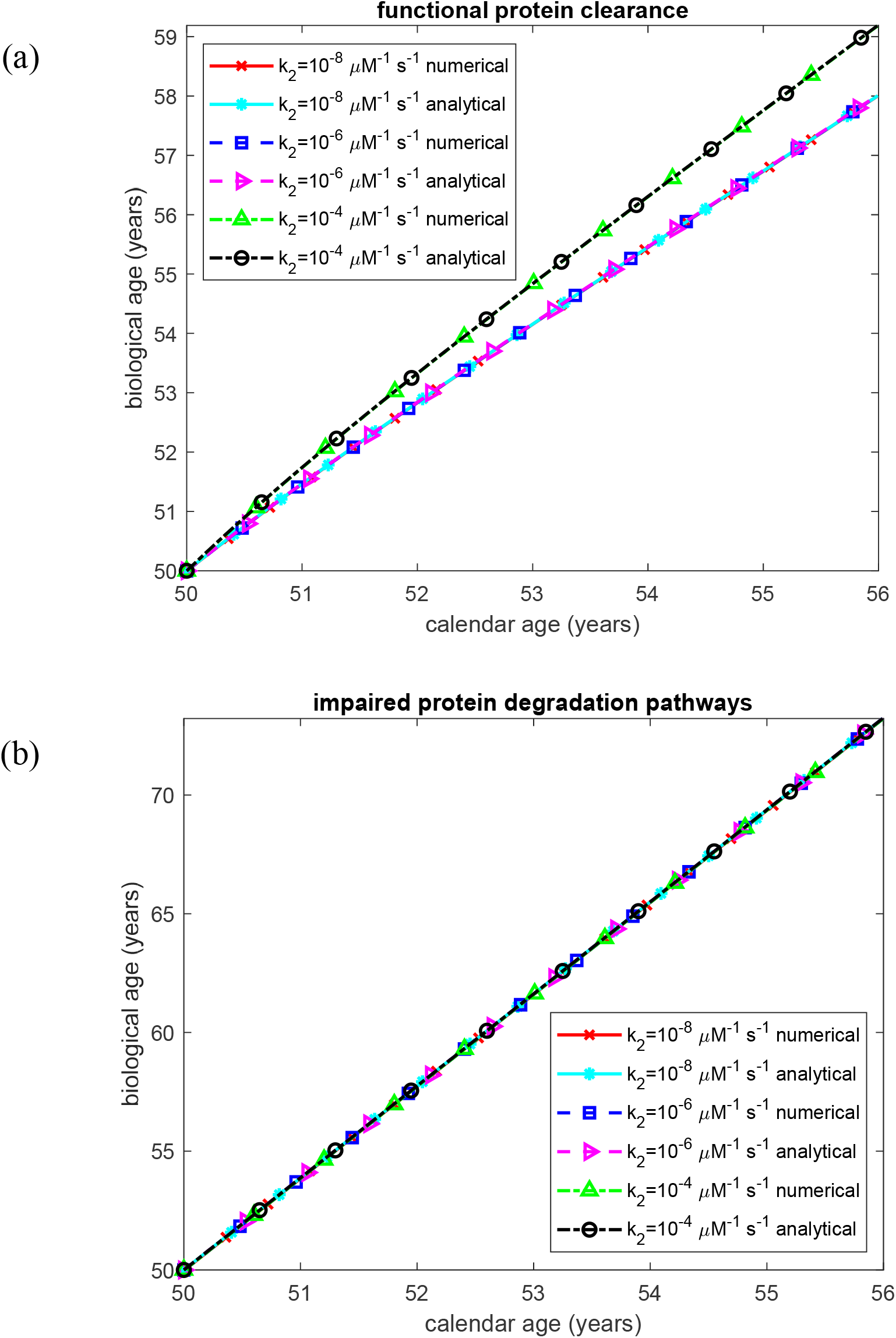
Biological age of the kidney as a function of time. (a) Biologically relevant half-lives *T*_1/2, *A*_, *T*_1/2,*B*_, *T*_1/2,*I*_, and *T*_1/2,*S*_, representing functional protein degradation machinery. (b) Infinite half-lives *T*_1/2, *A*_, *T*_1/2,*B*_, *T*_1/2,*I*_, and *T*_1/2,*S*_, representing complete impairment of protein degradation. Each panel shows results for three values of the rate constant for the second pseudo-elementary (autocatalytic growth) step in the F–W model of AA oligomer formation. Baseline parameters: *k*_1_ =10^−4^ s^−1^, *k*_*cl*_ = 9.65×10^−5^ s^−1^, *K*_*d*_ = 1.1789 *µ*M, *h*_*S*_ = 5.55×10^−7^ m s^−1^, *H*_*tot*_ = 30 *µ*M, *S*_*tot*_ = 8 *µ*M, and *θ*_1/2,*B*_ = 8.64 ×10^4^ s; remaining parameters as listed in Table 2.

**Fig. S26.**
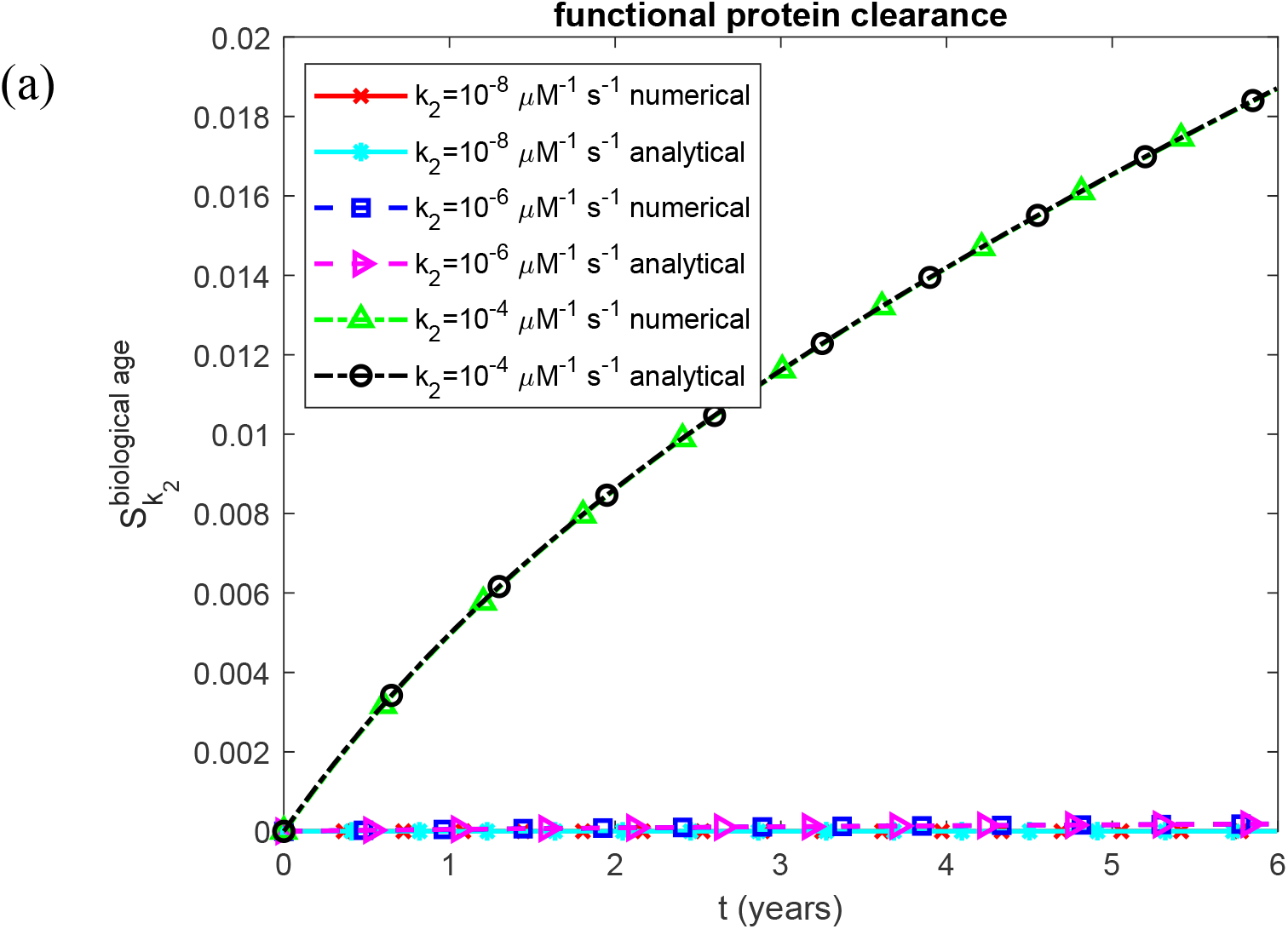
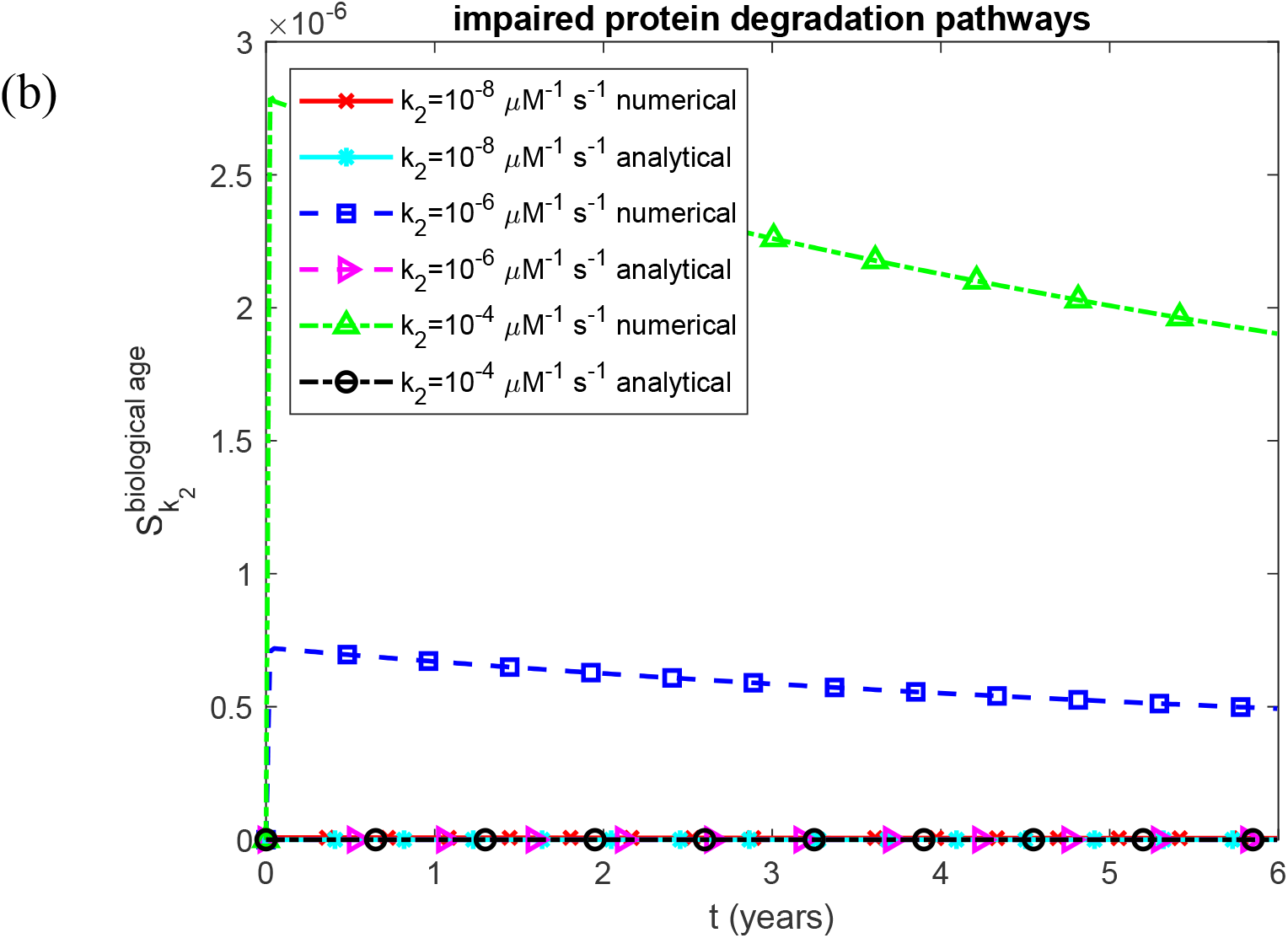
Dimensionless sensitivity of renal biological age to the rate constant for the second pseudo-elementary (autocatalytic growth) step in the F–W model of AA oligomer formation, as a function of time. (a) Biologically relevant half-lives *T*_1/2, *A*_, *T*_1/2,*B*_, *T*_1/2,*I*_, and *T*_1/2,*S*_, representing functional protein degradation machinery. (b) Infinite half-lives *T*_1/2, *A*_, *T*_1/2,*B*_, *T*_1/2,*I*_, and *T*_1/2,*S*_, representing complete impairment of protein degradation. Each panel shows results for three values of the rate constant for the second pseudo-elementary (autocatalytic growth) step in the F–W model of AA oligomer formation. Baseline parameters: *k*_1_ =10^−4^ s^−1^, *k*_*cl*_ = 9.65×10^−5^ s^−1^, *K*_*d*_ = 1.1789 *µ*M, *h*_*S*_ = 5.55×10^−7^ m s^−1^, *H*_*tot*_ = 30 *µ*M, *S*_*tot*_ = 8 *µ*M, and *θ*_1/2,*B*_ = 8.64 ×10^4^ s; remaining parameters as listed in Table 2.

**Fig. S27.**
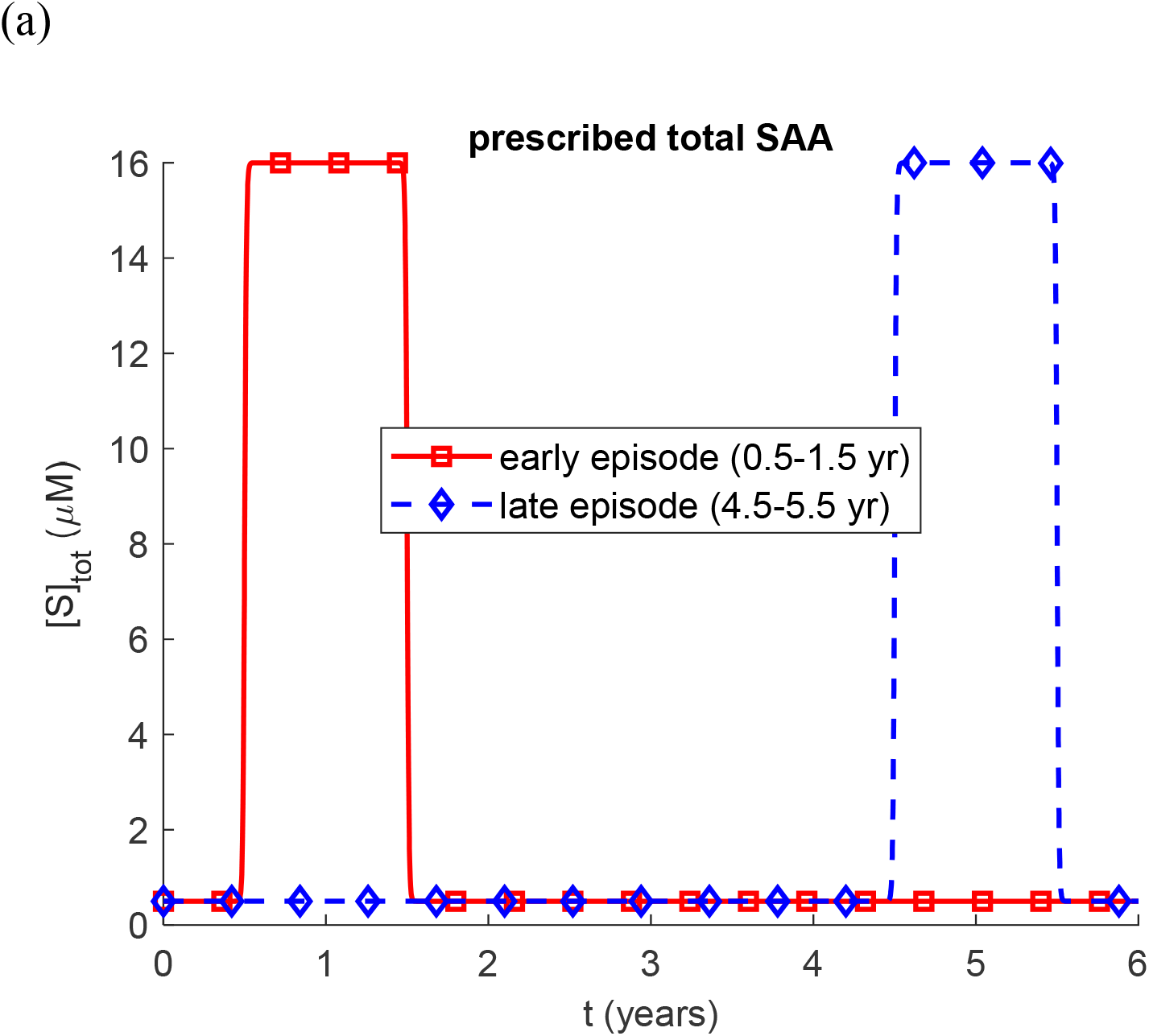
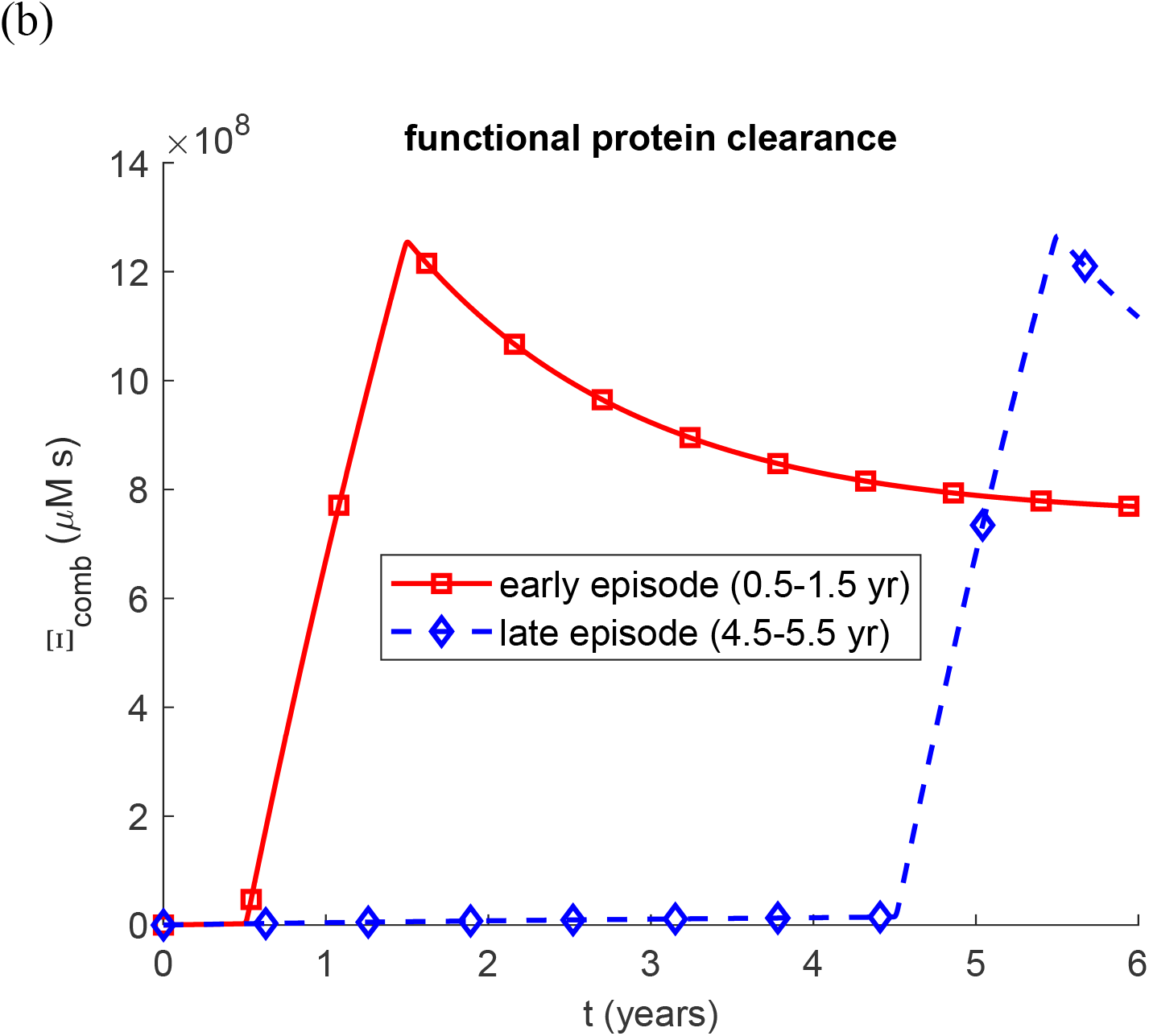
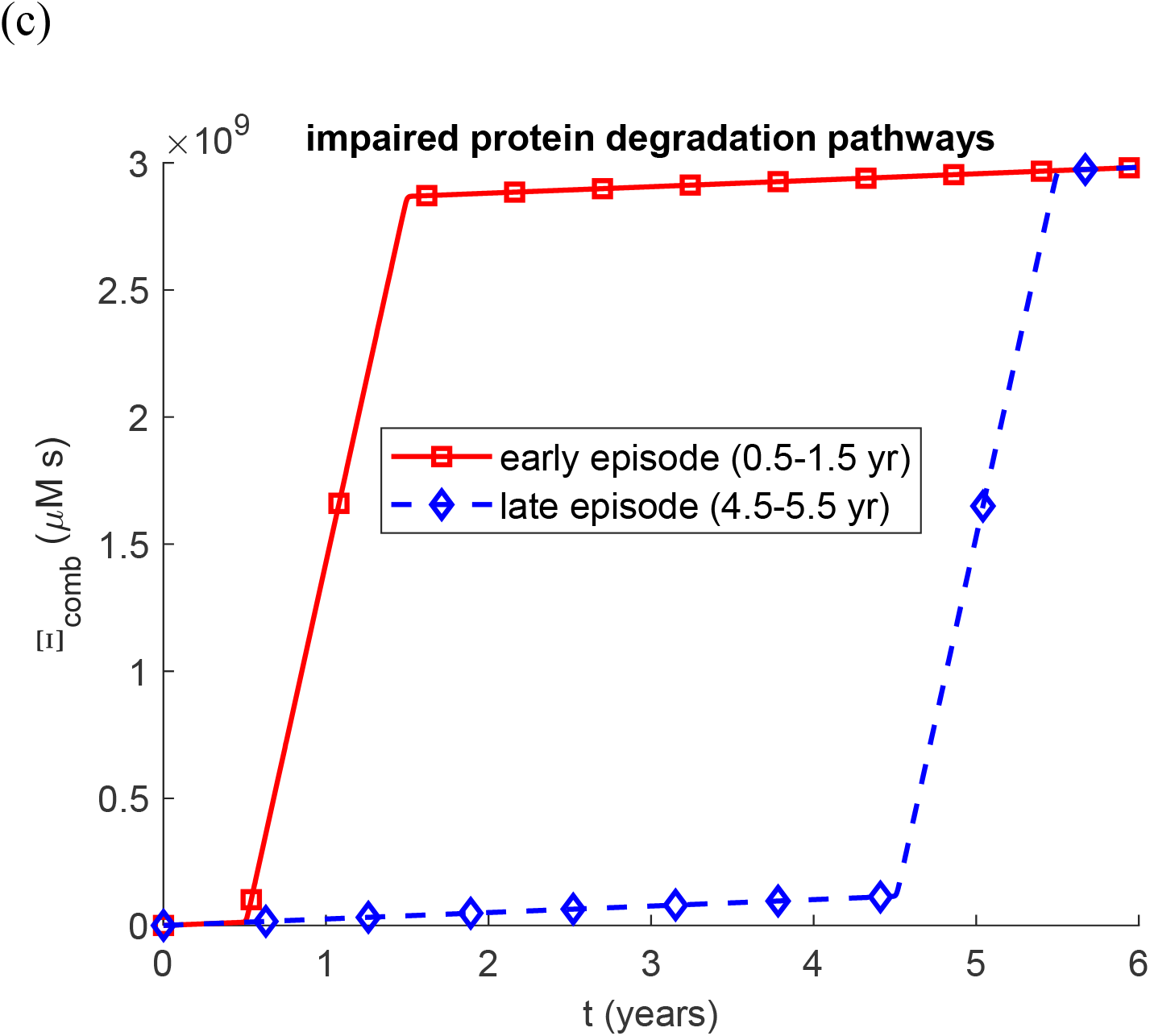
Combined renal damage for two scenarios in which an inflammatory episode of identical amplitude (peak total SAA 16 µM) and identical duration (one year) occurs either early (0.5–1.5 years) or late (4.5–5.5 years) in a six-year disease course. (a) Prescribed total SAA concentration; outside the episode, [*S*]_*tot*_ = 0.5 µM. (b) Biologically relevant half-lives, representing functional protein degradation machinery; the late episode produces 45% more combined damage at six years, the difference residing entirely in the fibrillar term, and the criterion is seen to decrease after the early episode as fibrils are cleared. (c) Infinite half-lives, representing complete impairment of protein degradation; the two scenarios then reach exactly the same combined damage at six years, because without clearance each contribution is a running total of oligomer exposure, and a total accumulated over the fixed six-year window does not depend on when within that window the episode occurs. Baseline parameters as in Fig. 2, with [*H*]_*tot*_ = 15 µM.

**Fig. S28.**
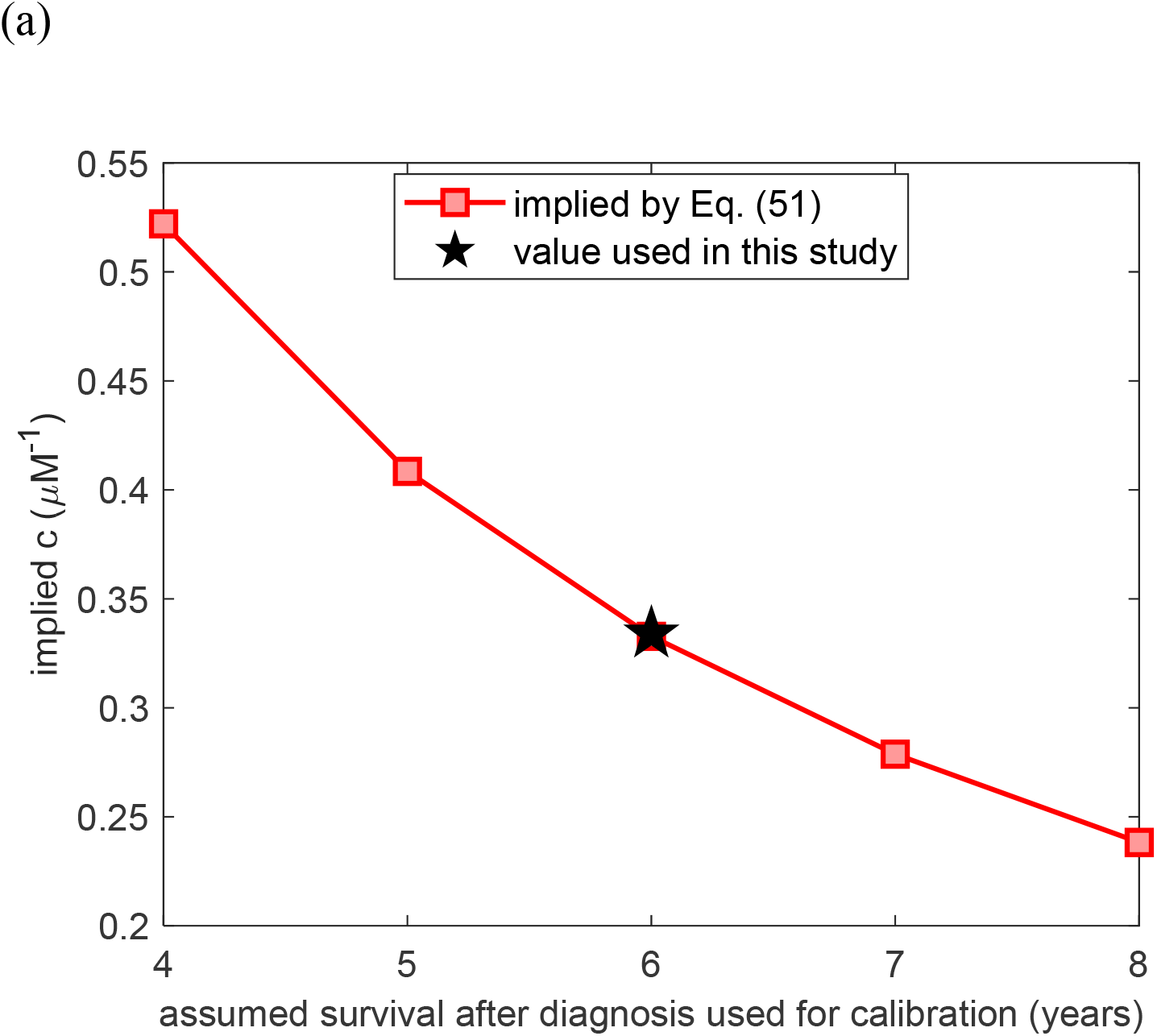
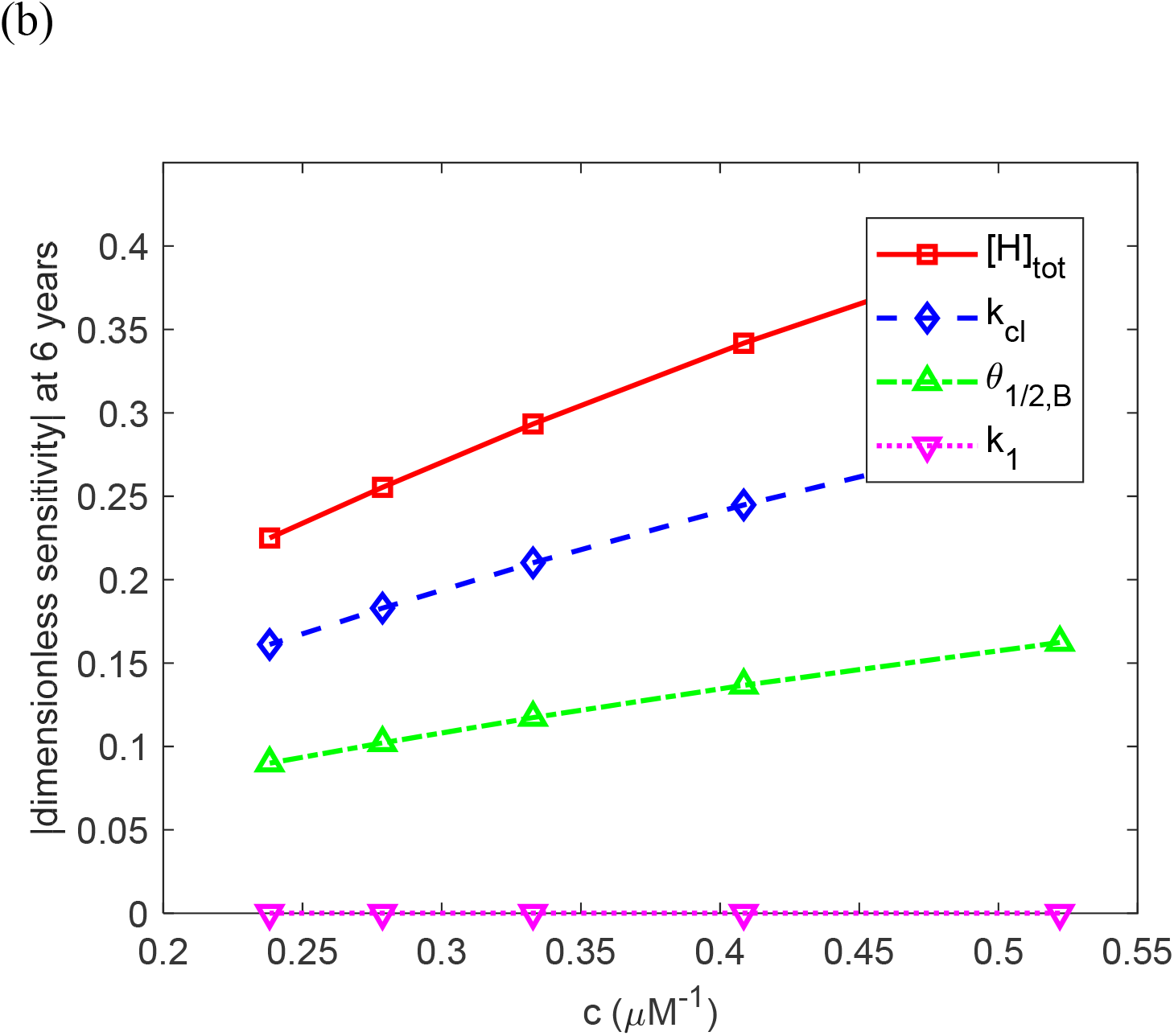
Dependence of the biological-age scale on the calibration of *c*. (a) Value of *c* implied by Eq. (51) as a function of the survival time after diagnosis assumed in the calibration scenario, over the reported 4–8-year range (Gillmore et al., 2001), (Ahbap et al., 2014); the star marks the value used in this study. (b) Magnitude of the dimensionless sensitivity of renal biological age at six years to four representative parameters, as a function of *c*. The magnitudes scale with *c*, but the signs of the coefficients, and the order of the parameters from most to least influential, are unchanged over the whole range.

